# Consistent Induction of Broadly Neutralizing HIV Antibodies by a Novel Two-Step Mechanism Informs Immunogen Design

**DOI:** 10.1101/2025.10.06.680687

**Authors:** Ashwin N. Skelly, Harry B. Gristick, Hui Li, Edem Gavor, Andrew J. Connell, Edward F. Kreider, Lorie Marchitto, Michael P. Hogarty, Maddy L. Newby, Joel D. Allen, Weimin Liu, Anthony P. West, Kasirajan Ayyanathan, Mary S. Campion, Kaitlyn Winters, Colette G. Gordon, Rebecca A. Osbaldeston, Macy J. Akeley, Yingying Li, Ajay Singh, Kendra Cruickshank, Younghoon Park, Chengyan Zhao, Xuduo Li, Khaled Amereh, Elizabeth Van Itallie, John W. Carey, Amie Albertus, Andrew T. DeLaitsch, Jennifer R. Keeffe, Melinda G. Lituchy, Agnes A. Walsh, Daniel J. Morris, Rumi Habib, Frederic Bibollet-Ruche, Nitesh Mishra, Gabriel Avillion, Nicholas S. Koranda, Samantha J. Plante, Christian L. Martella, Jinery Lora, Eric J. D. Wang, Mark G. Lewis, Malcolm A. Martin, Michel C. Nussenzweig, Michael S. Seaman, Darrell J. Irvine, Kevin J. Wiehe, Barton F. Haynes, Kshitij Wagh, Bette T. Korber, Raiees Andrabi, Max Crispin, Drew Weissman, Pamela J. Bjorkman, Beatrice H. Hahn, George M. Shaw

## Abstract

A major obstacle confronting HIV-1 vaccine and cure research is the lack of an outbred animal model for rapid and consistent induction of broadly neutralizing antibodies (bNAbs). We designed an epitope-focused simian-human immunodeficiency virus (SHIV.5MUT) that elicited broad and potent V3-glycan-targeted antibodies within a year of infection in 14 of 22 macaques compared with 0 of 14 control animals. SHIV.5MUT elicited bNAbs by a novel two-step mechanism, inducing an initial wave of V1-directed antibodies that selected for Envs with shortened, hypoglycosylated V1 loops, which in turn primed V3-glycan bNAb precursors. Rhesus bNAbs were immunogenetically and structurally diverse, closely resembling human V3-glycan bNAbs. Env-bNAb coevolution revealed a diverse repertoire of bNAb precursors and the Env variants that matured them, yielding a molecular blueprint for vaccine design.

## Main Text

Prophylactic administration of HIV-1 broadly neutralizing antibodies (bNAbs) to naïve rhesus macaques reliably protects them from heterologous simian-human immunodeficiency virus (SHIV) challenge (*1–4*). Similarly, in humans the bNAb VRC01 protects against infection by sensitive viral strains (*5, 6*). When given therapeutically to SHIV-infected monkeys or people living with HIV, bNAbs reduce plasma virus load and delay viral rebound after antiretroviral treatment interruption, in rare instances leading to a “functional cure” (*7–11*). These findings have led to intense efforts to elicit bNAbs by vaccination. While there has been progress, consistent induction of bNAbs in the plasma at titers that approach clinically protective thresholds has not been achieved. A major obstacle thus far has been the lack of an outbred animal model wherein potent bNAbs can be reproducibly elicited and their underlying developmental pathways elucidated and replicated. Such a model, if available, could serve as both a blueprint and a benchmark for iterative vaccine design. Here, we report the development of an epitope-focused SHIV strain (SHIV.5MUT) that rapidly induces V3-glycan bNAbs in the majority of infected monkeys with breadth, potency, and kinetics desirable in a human vaccine.

The V3-glycan patch on the HIV-1 envelope (Env) glycoprotein is one of the most frequently targeted bNAb sites in chronic human infection (*12*). This epitope includes the N332 and N301 glycans and the ^324^GDIR_327_ peptide motif at the base of the V3 loop, which contributes to coreceptor binding (*13–16*). V3-glycan antibodies are among the most potent bNAb classes (*17, 18*), often achieving geometric mean titers (GMT) of <0.1 µg/mL IC_50_ (half-maximal inhibitory concentration) against large panels of heterologous tier-2 viruses, with the broadest members neutralizing up to 65% of circulating strains (*15, 17–22*). Unlike bNAbs targeting other Env epitopes, such as the CD4 binding site (CD4bs) and the trimer apex (V2), V3-glycan bNAbs are structurally and immunogenetically diverse, exhibiting widely varying angles of approach and less reliance on uncommon features like long heavy chain complementarity-determining region 3 (CDRH3) loops and restricted V_H_ or D gene segment usage (*16–18*). Thus, V3-glycan bNAbs arise from a less restricted pool of germline precursors and may be easier to elicit by vaccination in human populations, which have heterogeneous immunoglobulin repertoires. Additionally, compared to other bNAb classes, V3-glycan bNAbs tend to require less extensive somatic hypermutation to acquire neutralization breadth (*15, 18, 19*), suggesting they could be more rapidly matured once primed. Altogether, these features make V3-glycan bNAbs a promising vaccine target.

The most common strategy for eliciting V3-glycan bNAbs thus far has been to use reverse vaccinology approaches, in which the germline precursor of a mature bNAb is inferred and used as a template to design immunogens that selectively bind to it (*18, 23*). However, such “lineage-based” strategies are limited by a paucity of high-confidence inferences of V3-glycan bNAb unmutated common ancestors (UCAs), which require longitudinal B cell receptor (BCR) sequencing datasets that are often not available. Inferred germlines (iGLs) have been used in their stead, but these typically retain most of their mature CDRH3 sequence which comprises non-templated V-D and D-J junctions. This is particularly problematic when targeting V3-glycan bNAb lineages, which generally bind in a CDRH3-dominant manner (*13–16*). Additionally, engineering an immunogen to maximize affinity for a single bNAb UCA/iGL does not ensure cross-reactivity with other bNAb precursors that target the same epitope. Despite these limitations, advances have been made using immunogens specifically engineered to engage precursors of the human V3-glycan bNAbs PGT121 (*24–27*), DH270 (*28, 29*), and BG18 (*30–32*). Sequential immunization in immunoglobulin knockin mouse models has provided proof-of-principle that these types of lineages can be primed and boosted *in vivo* (*25, 28, 32*), though consistent V3-glycan bNAb induction resulting in protective plasma titers has not yet been achieved in outbred animals (*27, 31*).

As an alternative to these lineage-based designs, we describe here a lineage-agnostic, epitope-focused approach to eliciting V3-glycan bNAbs. Previously, we hypothesized that immunizing with BG505.N332 Env variants lacking N-glycans at residues N133/N137/N156 (RC1) (*26*) or N133/N137 (11MUTB) (*24*) might selectively activate V3-glycan bNAb precursors, and that further boosting with more native-like immunogens, including 5MUT and a cocktail of wildtype Envs, would promote their maturation to breadth (*27*). This approach was only partially successful, eliciting antibodies that exhibited weak cross-neutralization and bound heterologous viruses with open or occluded-open Env conformations (*27, 33, 34*). Additionally, structural analyses suggested that while the initial humoral response was focused to the V3 region, it became increasingly off-target with subsequent boosting (*27*). In the current study, we substituted the boosting immunogen (5MUT) with an infectious SHIV bearing the corresponding Env. We reasoned that infection with SHIV.5MUT might better mature RC1/11MUTB-primed responses by acting as an “evolving immunogen” given its persistent replication, high antigenic load, and ability to coevolve with the humoral response. Surprisingly, we found that regardless of prior vaccination, 14 of 22 (64%) SHIV.5MUT-infected macaques developed potent V3-glycan bNAb responses within one year of infection in contrast to 0 of 14 (0%) control animals infected with SHIV.BG505.N332, despite differing by only four residues in the V1 region of Env (p<0.0001, Fisher’s exact test). bNAb induction occurred via a two-step mechanism in which an early wave of V1-directed antibodies selected for Env variants with shortened V1 loops, which in turn primed and expanded V3-glycan bNAb precursor B cells. This new outbred animal model allowed us to dissect the critical events involved in V3-glycan bNAb elicitation, beginning with the priming of multiple precursors and followed by affinity maturation to breadth and potency. Importantly, we identified shared routes of Env-bNAb coevolution across multiple animals, revealing a “molecular blueprint” for V3-glycan bNAb induction that can guide vaccine design efforts.

### Consistent induction of plasma breadth in SHIV.5MUT-infected macaques

Macaques were sequentially immunized with RC1 and 11MUTB as either protein nanoparticles (Group 1, n=8) or mRNA-LNPs (Group 2, n=8) and then infected with SHIV.5MUT eight weeks later. Two additional groups of unimmunized macaques were infected with either SHIV.5MUT (Group 3, n=12) or wildtype SHIV.BG505.N332 (Group 4, n=14) **(Fig. 1A, Supplementary Table 1)**. Details of protein and mRNA immunogen design are provided in the **Supplementary Text (fig. S1A-B)**. Vaccinated animals developed robust autologous neutralizing plasma responses within eight weeks of RC1 prime, with mean titers significantly higher in protein-versus mRNA-immunized animals **(fig. S1C)**. Titers increased following booster immunization with 11MUTB, with a dose escalation subgroup of protein-vaccinated macaques reaching significantly higher titers than the bolus subgroup **(fig. S1C)**. Following protein or mRNA vaccination, there was no cross-neutralization of viruses bearing 5MUT and BG505.N332 Envs, indicating successful immunofocusing of the initial B cell response to the glycan-depleted V1V3 region **(fig. S1C)**.

**Fig. 1.**
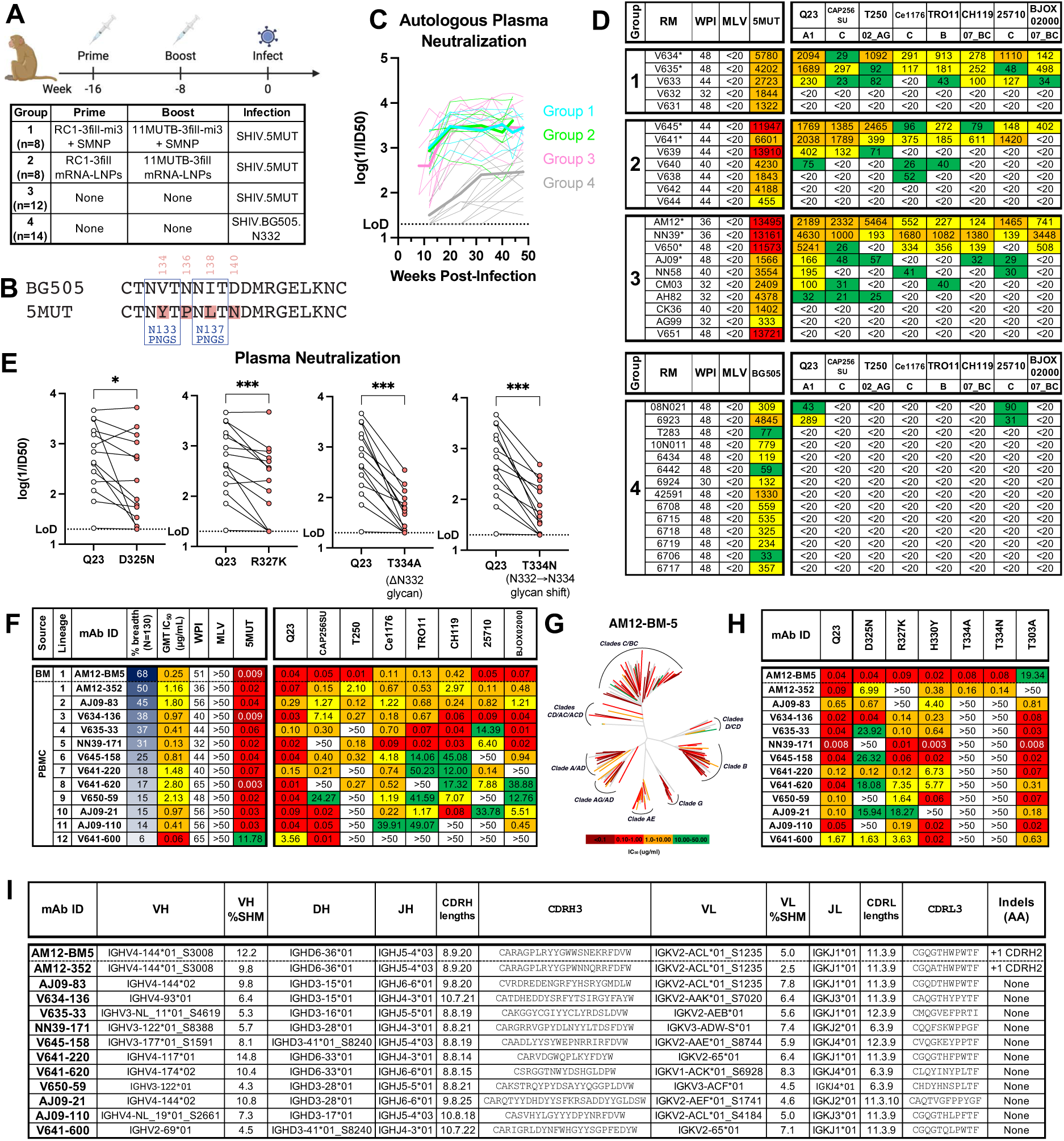
SHIV.5MUT rapidly and consistently induces V3-glycan bNAbs. **(A)** Study design. Macaques were immunized with 100 ug of protein nanoparticles adjuvanted with 750 U SMNP (delivered either as a single bolus or via an escalating dose regimen over the course of two weeks) or 100 ug of mRNA-LNPs (delivered as a single bolus) at eight-week intervals. All vaccinated as well as unvaccinated control animals were then infected with SHIV.5MUT or SHIV.BG505.N332. **(B)** Amino acid alignment of the BG505 and 5MUT Envs in the V1 region. Differences at positions 134, 136, 138, and 140 are highlighted in red, and PNGS sites are boxed. **(C)** Autologous plasma neutralization over time. Autologous plasma neutralization titers are shown for SHIV.5MUT-(Groups 1-3) and SHIV.BG505.N332- (Group 4) infected macaques, represented as log(1/ID_50_). Thin lines represent values from individual animals, and thick lines represent the group mean. **(D)** Development of plasma breadth. Plasma neutralization titers (represented as reciprocal ID_50_) are shown for each animal against eight heterologous viruses representing different M-group clades at the timepoint of greatest breadth within 48 weeks post-infection (WPI). Murine leukemia virus (MLV) is used as a negative control. Asterisks indicate animals from which bNAbs were isolated. **(E)** Mapping of the heterologous neutralizing response. Plasma samples from 14 SHIV.5MUT-infected macaques were tested against Q23 mutants lacking key V3-glycan epitope residues, with titers expressed as log(1/ID_50_). Wilcoxon matched-pairs signed-rank test was used. *P < 0.05; ***P < 0.001. **(F)** Neutralization profiles of monoclonal antibodies representing the twelve V3-glycan bNAb lineages isolated from eight SHIV.5MUT-infected macaques. Titers are expressed as IC_50_ in µg/mL. All antibodies were tested up to 50 µg/mL **(G)** Neutralization breadth and potency of the AM12-BM5 mAb against a 130-strain global virus panel, with a dendrogram illustrating the phylogenetic relatedness of tested Envs. Neutralization potency is indicated with a heatmap. **(H)** Epitope mapping of rhesus bNAbs using a panel of Q23 mutant viruses lacking key V3-glycan epitope residues. Titers are expressed as IC_50_ in µg/mL. **(I)** Immunogenetic features of rhesus V3-glycan bNAbs. V, D, and J gene segment usage is indicated. SMNP, saponin/MPLA nanoparticle; LNP, lipid nanoparticle; PNGS, potential N-linked glycosylation site; WPI, weeks post-infection; MLV, murine leukemia virus; RM, rhesus macaque; BM, bone marrow; PBMC, peripheral blood mononuclear cell; SHM, somatic hypermutation.

Next, we aimed to mature these vaccine-elicited responses by infecting with the “evolving immunogen” SHIV.5MUT **(Fig. 1A)**. The 5MUT Env differs from BG505 at only four positions in gp160: V1 loop residues V134Y, N136P, I138L, and D140N **(Fig. 1B)**. These substitutions were originally selected to enhance binding to a partially germline-reverted form of PGT121 (*24*). To verify that the 5MUT Env assembled well on virions, we determined the infectivity of SHIV.5MUT in TZM-bl cells and confirmed its tier-2, BG505.N332-like neutralization sensitivity. Antibodies targeting regions exposed on Envs with more open conformations such as CD4i, V2i, and linear V3 epitopes (*35–37*) generally failed to neutralize SHIV.5MUT. Conversely, V2-apex bNAbs that recognize quaternary epitopes on the Env trimer potently neutralized SHIV.5MUT **(fig. S1D)**. Together, these results confirm expression of intact, well-formed Env trimers on the virion surface. Additionally, SHIV.5MUT exhibited enhanced susceptibility to neutralization by multiple V3-glycan bNAbs, indicating that the mutations in 5MUT increase accessibility of this epitope as previously reported (*24*) **(fig. S1D)**.

Animals in Groups 1, 2, and 3 were intravenously inoculated with SHIV.5MUT and animals in Group 4 were inoculated with SHIV.BG505.N332. Productive infection ensued in all 42 macaques, with most animals reaching peak plasma viremia of 10^7^-10^8^ vRNA copies per mL at two weeks post-infection and a stable setpoint thereafter **(fig. S1E)**. Five macaques with sustained high viral loads rapidly progressed to AIDS and one macaque controlled its infection, exhibiting low-level viremia. These six animals were excluded from further analysis. The remaining 36 macaques were followed for one year and were serially screened for plasma neutralization activity against SHIV.5MUT, SHIV.BG505.N332, and a panel of eight heterologous, tier-2 viruses that contain a potential N-linked glycosylation site (PNGS) at position N332, which is required for neutralization by most V3-glycan bNAbs (*21*).

All three groups of SHIV.5MUT-infected animals developed potent autologous neutralizing responses within the first 12 weeks of infection that increased over time, plateaued by week 20, and persisted until at least week 48 **(Fig. 1C)**. SHIV.BG505.N332-infected animals exhibited similar responses but with lower autologous neutralization titers **(Fig. 1C)**. Surprisingly, we began observing heterologous neutralization activity in the plasma of SHIV.5MUT-infected macaques as early as week 20 post-infection, regardless of vaccination history **(fig. S2)**. Overall, 14 of 22 (64%) SHIV.5MUT-infected macaques developed bNAb responses, defined as neutralization of at least 3 out of 8 heterologous viruses with ID_50_ titers >1:20 within 48 weeks of infection. Eight animals neutralized at least 6 out of 8 viruses in the panel with ID_50_ titers frequently exceeding 1:1000 (average GMT ID_50_ = 1:507) **(Fig. 1D)**. All bNAb responses mapped to the V3-glycan epitope, as evidenced by substantial reduction or complete abrogation of neutralizing activity against mutant heterologous viruses lacking critical V3 residues including D325, R327, and N332 **(Fig. 1E)**. In contrast, 0 of 14 (0%) SHIV.BG505.N332-infected macaques developed plasma breadth within the same 48-week timeframe (p<0.0001, Fisher’s exact test) **(Fig. 1D)**. As the 5MUT Env differs from BG505 by only four amino acid substitutions in the V1 loop **(Fig. 1B)**, these mutations must endow SHIV.5MUT with its unique propensity to elicit V3-glycan bNAbs. Notably, the frequency of bNAb elicitation in vaccinated (Groups 1-2) versus unvaccinated (Group 3) SHIV.5MUT-infected animals was not significantly different (p=0.67, Fisher’s exact test), suggesting that SHIV.5MUT or a derivative induced these bNAb lineages rather than the RC1 or 11MUTB immunogens **(Fig. 1D)**. Together, these data demonstrate that SHIV.5MUT rapidly and consistently induces broad and potent V3-glycan-directed neutralizing responses in macaques.

### Isolation of immunogenetically diverse V3-glycan bNAbs

We next sought to isolate and characterize the monoclonal antibody (mAb) lineages responsible for the observed plasma breadth. We used fluorescence-activated cell sorting to isolate heterologous Env-binding IgG^+^ memory B cells from peripheral blood mononuclear cells (PBMCs) of the eight macaques with maximal plasma breadth **(fig. S3A)**. Single Env-specific B cells were sorted into individual wells of a 96-well plate and antibody genes were amplified and sequenced (*38, 39*). Representatives of expanded lineages were synthesized as recombinant mAbs and screened for neutralization activity against SHIV.5MUT and a panel of four heterologous viruses **(fig. S3B)**. Antibodies that potently neutralized at least two heterologous viruses were selected for epitope mapping and further testing against a larger panel of viruses. In total, we screened 238 mAbs representing 106 distinct lineages **(fig. S3B)**, yielding twelve V3-glycan bNAb lineages **(Fig. 1F)**. We also sorted bone marrow plasma cells from the macaque with the greatest plasma breadth **(fig. S3C)** and identified additional bNAb lineage members, one of which (AM12-BM5) potently neutralized all viruses in the screening panel **(Fig. 1F)** (*40*).

Neutralization profiles of bNAbs from each of the twelve lineages generally recapitulated the plasma breadth observed in the corresponding macaque, indicating successful identification of the predominant bNAb species (**Fig. 1D** and **F**). To characterize the breadth of these bNAbs more comprehensively, we assessed their neutralization against a diverse, 130-virus panel containing representatives of all major HIV-1 clades. The breadth of these SHIV.5MUT-elicited V3-glycan bNAbs ranged from 6-68% with GMT IC_50_ of 0.06-2.80 μg/mL **(Fig. 1F, fig. S4)**. When tested only against viruses containing an N332 glycan (n=89), breadth ranged from 9-75% **(fig. S4)**. Such breadth and potency are comparable to those of prototypical human V3-glycan bNAbs including DH270.6 (51%, 0.21 μg/mL) and PCDN76-33A (46%, 0.50 μg/mL). It is not surprising that the very best human V3-glycan bNAbs PGT121 (66%, 0.07 μg/mL), PGT128 (68%, 0.06 μg/mL), and BG18 (61%, 0.03 μg/mL) exhibit greater potencies, as they were isolated after many years of chronic HIV-1 infection and were identified retrospectively by screening large human cohorts for individuals with the greatest neutralization breadth and potency (*18, 20, 22, 41*). In contrast, the rhesus V3-glycan bNAbs reported here were identified prospectively within the first year of SHIV infection. Much like human V3-glycan bNAbs, almost all SHIV.5MUT-elicited bNAbs were restricted in their neutralization breadth to viruses containing an N332 glycan. The exceptions were AM12-352 and its bone marrow plasma cell-derived relative AM12-BM5, which were able to neutralize viruses lacking the N332 glycan **(fig. S4)**, thereby broadening their coverage to include many clade AE strains that circulate in Asia **(Fig. 1G, fig. S5A-B)** (*21, 42*).

To further map the SHIV.5MUT-elicited bNAbs, we tested their neutralization activity against a series of Q23.17 mutants that lacked known contact residues in the canonical V3-glycan epitope. Despite targeting the same overall V3 region of Env, the twelve bNAb lineages differed in their reliance on particular residues within the epitope. Consistent with the 130-virus neutralization profiles **(fig. S4, Fig. 1G)**, all bNAbs exhibited a strict dependence on the N332 glycan except AM12-352 and AM12-BM5, which instead required the N301 glycan **(Fig. 1H)**. This N301-dependence is desirable because the N301 glycan, unlike the N332 glycan, is highly conserved across all HIV-1 subtypes **(fig. S5B)**. Additionally, representative mAbs from all lineages required D325 or R327 (which are part of the ^324^GDIR_327_ peptide motif) or the downstream H330, although the pattern and extent of dependence varied. Interestingly, this diversity in the functional epitope was observed both between and within individual animals. For example, we isolated three different V3-glycan bNAb lineages from each of two macaques (AJ09 and V641), and representative antibodies from each of these lineages exhibited distinct patterns of dependence on D325, R327, and H330 **(Fig. 1H)**. We also observed intra-lineage heterogeneity in epitope recognition, with several members of the AM12-352 lineage depending completely on the N332 glycan instead of the N301 glycan **(fig. S6A)**. To be most protective, an immunization regimen should ideally accomplish something similar, as inducing multiple bNAb lineages or sub-lineages that target the same epitope in different ways may protect against infection by viral strains that have evolved to escape a given class of V3-glycan bNAbs.

To better understand the high frequency of V3-glycan bNAb elicitation in SHIV.5MUT-infected macaques, we analyzed the immunogenetic features of the antibodies. We found that SHIV.5MUT-induced V3-glycan bNAbs were immunogenetically diverse, utilizing various V_H_3- and V_H_4-family gene segments **(Fig. 1I)**, which comprise the most common V_H_ alleles in both humans and macaques (*43–45*). Thus, there does not seem to be a strict requirement for certain immunoglobulin alleles, unlike V2-apex and CD4-mimetic bNAbs (*46–49*). The range in CDRH3 length was also surprisingly wide (14-25 amino acids, with a median of 20) and well within the range of typical rhesus and human antibodies, as 7% and 13% of peripheral B cells, respectively, express a BCR with CDRH3 ≥20 amino acids **(Fig. S6B)** (*50–52*). Additionally, most V3-glycan bNAb lineages did not exhibit any indels except for the AM12-352 lineage, which acquired multiple independent insertions in CDRH2 **(Fig. 1I)**. The average frequency of V_H_ somatic mutation was 8.4% at the nucleotide level **(Fig. 1I)**, which is less than half that of previously described V3-glycan bNAbs (*18*), implying that once primed, these lineages may not require extensive maturation to achieve breadth and potency. Altogether, the immunogenetic diversity of the SHIV.5MUT-induced bNAbs and lack of apparent rare features likely contribute to the high prevalence of bNAb elicitation in this model system.

### Structural features of macaque V3-glycan bNAbs

To explore the structural implications of the remarkable immunogenetic diversity of rhesus V3-glycan bNAbs, we selected ten V3-glycan bNAbs representing seven lineages and determined single-particle cryo-EM structures of their Fabs in complex with soluble 5MUT Env trimer (*53*) **(Fig. 2A, fig. S7A,** and **Supplementary Table 2)**. Consistent with the neutralization-based mapping (**Fig. 1H** and **fig****. S6A)**, all ten rhesus bNAbs recognized the V3-glycan epitope, contacting residues in ^324^GDIR_327_ as well as the N332_gp120_ glycan **(Fig. 2B, fig. S7B)** in a manner similar to human V3-glycan bNAbs **(Fig. 2C)**. All SHIV.5MUT-elicited bNAbs primarily contacted the ^324^GDIR_327_ motif through tyrosine-mediated interactions **(Fig. 2B)**. Although tyrosine residues are also used by human V3-glycan bNAbs to contact ^324^GDIR_327_ **(Fig. 2C)**, the rhesus bNAbs used a more varied approach to interact with this motif (*54*). A critical interaction observed in human V3-glycan bNAb structures involves a negatively charged residue within CDHR3 (e.g., Glu100I_HC_ in 10-1074) contacting the conserved R327_gp120_ residue within the ^324^GDIR_327_ motif **(Fig. 2C)** (*54*). Interestingly, two SHIV.5MUT-elicited bNAbs, AJ09-83 and AJ09-110, recapitulated this interaction **(Fig. 2B)**.

**Fig. 2.**
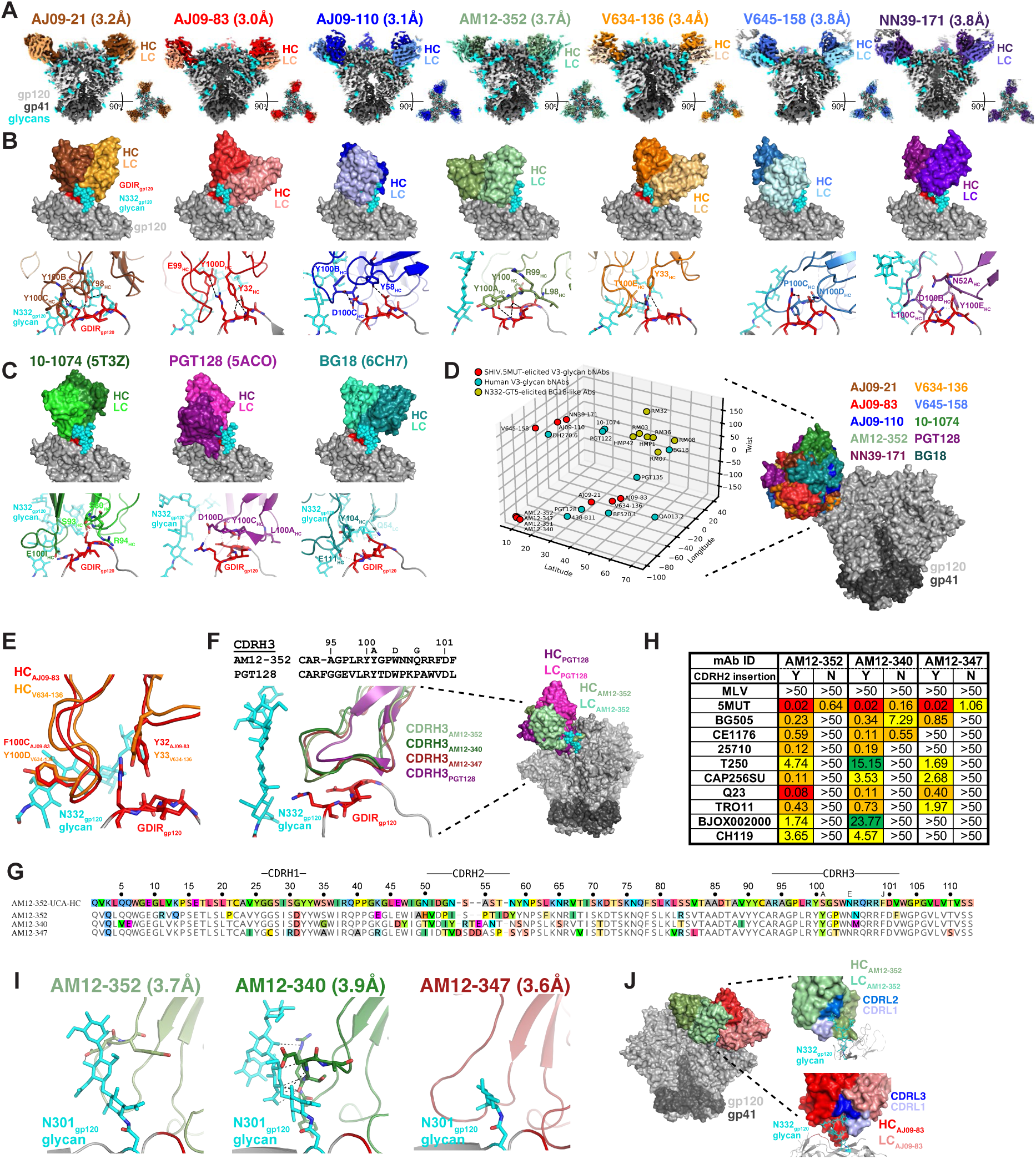
SHIV.5MUT-induced V3-glycan bNAbs are structurally diverse and resemble human bNAbs. **(A)** Cryo-EM maps of seven SHIV.5MUT-induced V3-glycan bNAbs in complex with 5MUT-3fill. The heavy chain of each bNAb is shown in a darker shade while the light chain is shown in a lighter shade. The N332 glycan is modeled as cyan spheres and the conserved GDIR peptide motif is red. **(B)** Top, surface representations of the V3-glycan bNAbs from **(A)** bound to gp120, with the N332 glycan and GDIR motif as depicted in **(A)**. Bottom, interfaces of the predicted interactions between the V3-glycan bNAbs and the GDIR motif. **(C)** Top: Surface representations of three well-characterized V3-directed bNAbs isolated from humans: 10-1074 (PDB ID: 5T3Z), PGT128 (PDB ID: 5ACO), and BG18 (PDB ID: 6CH7). Bottom: Potential interactions between the V3-glycan bNAbs and the GDIR motif (red) and N332 glycan (cyan) of gp120. **(D)** Comparative analysis of the angles of approach between V3-glycan bNAbs elicited by SHIV.5MUT infection (red), V3-glycan bNAbs isolated from humans (blue), and putative V3-glycan bNAb precursors isolated from N332-GT5-immunized macaques (*31*). Antibodies cluster based on similarity in approach angle. **(E)** Structural overlay of AJ09-83_HC_ and V634-136_HC_ demonstrating shared pairwise interactions with the GDIR motif and N332_gp120_ glycan within the V3 epitope. **(F)** Left: Amino acid sequence alignment (top) and structural overlay (bottom) of the CDRH3 loops from AM12-352 and PGT128. Right: Structural overlay of AM12-352 and PGT128 binding to the Env V3-glycan epitope. **(G)** Alignment of the AM12-352 lineage UCA heavy chain amino acid sequence with those of mature bNAbs representing the sub-lineages. Differences from the UCA, including indels in CDRH2, are highlighted. **(H)** Influence of CDRH2 insertions on neutralization breadth. The indicated antibodies were produced with (Y) and without (N) their respective CDRH2 insertions. Neutralization activity against a panel of viruses is shown, with titers expressed as IC_50_ in µg/mL. **(I)** Predicted interactions between the CDRH2 insertions of AM12-352, AM12-340, and AM12-347 with the N301_gp120_ glycan. **(J)** Left, structural overlay of AM12-352 and AJ09-83. Right, predicted interactions between the CDRL loops of AM12-352 (top) and AJ09-83 (bottom) with the N332_gp120_ glycan. HC, heavy chain; LC, light chain.

To evaluate the range of binding poses of the SHIV.5MUT-induced bNAbs, we calculated approach angles for the Fab-trimer complex structures and compared these to angles of human V3-glycan bNAbs as well as V3-directed antibodies elicited in macaques by N332-GT5, a germline-targeting immunogen designed to stimulate BG18-like precursors **(Fig. 2D, fig. S7C)** (*31*). We found that SHIV.5MUT-elicited bNAbs approached the epitope from a wide range of angles, much like human V3-glycan bNAbs. In contrast, antibodies elicited by N332-GT5 exhibited a narrow range of approach angles that resembled that of BG18, consistent with their restricted immunogenetics (*31*).

We also assessed whether any of the SHIV.5MUT-elicited bNAbs shared a common binding mode, as was found for VRC01-class bNAbs that recognize the CD4 binding site (*55*). Evidence for a common binding mode includes the occurrence of multiple, shared pairwise Env-antibody residue interactions. Antibodies with very different angles of approach are unlikely to share such pairwise interactions, except when these interactions involve similar CDRH3 loops that have different orientations with respect to the antibody V_H_ domain. Of the seven representative SHIV.5MUT-elicited bNAb lineages that we structurally characterized **(Fig. 2A-B)**, only AJ09-83 and V634-136 showed substantial shared pairwise interactions **(Fig. 2E)**. These include Phe100C_HC AJ09-83_ / Tyr100D_HC V634-136_, which interact with the N-acetylglucosamine of the N-glycan attached to N332_gp120_, and Tyr32_HC AJ09-83_ / Tyr33_HC V634-136_, which interact with the R327_gp120_ sidechain and the I326_gp120_ backbone. Interestingly, the Phe100C_HC AJ09-83_ / Tyr100D_HC V634-136_ residues are both contributed by the “YYY” motif encoded by the D3-15*01 gene segment in reading frame 2 **(Fig. 1I)**, although the common pairwise interactions occur at adjacent positions within this motif. Additionally, the shared tyrosine-mediated interactions at positions 32_HC AJ09-83_ and 33_HC V634-136_ are part of CDRH1 loops encoded by distinct V_H_ gene segments.

Despite the extensive diversity of the rhesus V3-glycan bNAbs, we found many examples of structural and immunogenetic similarities with human bNAbs. For example, although PGT128 (human) and AM12-352 (macaque) exhibited different angles of approach, their CDRH3 loops formed structurally similar architectures and interacted with the ^324^GDIR_327_ motif in an analogous manner **(Fig. 2F)**. While the overall CDRH3 amino acid sequence identity between PGT128 and AM12-352 was low (28.6%), both loops included an LRY motif that formed nearly identical architectures and made similar interactions with ^324^GDIR_327_ **(Fig. 2F)**. Additionally, the evolution of the AM12-352 lineage mirrored that of the PGT128 lineage – both split into sub-lineages defined by CDRH2 insertions that contributed to neutralization breadth (*56*). We isolated several members of the AM12-352 lineage, which segregated into distinct clades characterized by unique insertions in CDRH2 **(Fig. 2G, fig. S8)**. Longitudinal BCR repertoire sequencing of peripheral IgG^+^ B cells revealed four expanded sub-lineages that split between 24 and 32 weeks post-infection following independent insertion events **(Fig. 2G, fig. S8)**. Representatives of each sub-lineage (AM12-BM5, AM12-352, AM12-340, and AM12-347) demonstrated distinct neutralization profiles, covering 68%, 50%, 32%, and 9% of the 130-virus panel **(fig. S4)**. AM12-BM5 and AM12-352, which independently acquired an identical CDRH2 insertion **(fig. S8)**, were the only antibodies capable of neutralizing viruses lacking the N332_gp120_ glycan and exhibited a strong functional dependence on the N301_gp120_ glycan (**Fig. 1H** and **fig****. S4)**. Removing the CDRH2 insertion from AM12-352 and two other lineage members with different insertions (AM12-340 and AM12-347) severely diminished their neutralization breadth and potency **(Fig. 2H)**. Fab-Env structures of these same lineage members showed that the insertions formed completely different architectures and orientations, and thus each interacted with the N301_gp120_ glycan in a unique manner. The CDRH2 regions of AM12-352 and AM12-340 formed curved loops that stack against the face of the N301_gp120_ glycan, whereas that of AM12-347 formed a straighter loop that packs against the side of the glycan **(Fig. 2I)**. This is reminiscent of the PGT128 lineage, in which a CDRH2 insertion increased contact with the N332_gp120_ glycan and enhanced breadth against viruses containing this glycan, whereas lineage members without the insertion (such as PGT130) could better neutralize N334-glycan-containing viruses (*56*). Finally, while SHIV.5MUT-elicited V3-glycan bNAbs used diverse immunoglobulin gene segments, two bNAbs from different macaques, AM12-352 and AJ09-83, had nearly identical light chains, both using the IGKV2-ACL*01-S1235 and IGKJ1*01 alleles **(Fig. 1I)**. Structural overlay revealed that both light chains engaged the N332_gp120_ glycan, though the Fabs recognized opposing sides of the glycan **(Fig. 2J)**. While both AM12-352 and AJ09-83 interacted with the N332_gp120_ glycan using CDRL1, AM12-352 also used CDRL2 while AJ09-83 also used CDRL3.

### Ontogeny of rhesus V3-glycan bNAbs

Robust inferences of bNAb UCAs and intermediate ancestors define the critical stages in bNAb development and can serve as valuable guides for immunogen design (*19, 28*). We thus investigated the ontogeny of the SHIV.5MUT-induced V3-glycan bNAbs. An advantage of the SHIV model is that it enables the longitudinal BCR analyses necessary to infer high-confidence UCAs, which in turn is critical to identifying Env variants capable of priming and boosting these lineages. To generate an individualized immunoglobulin repertoire reference for each macaque, we deeply sequenced the BCRs of naïve IgM^+^IgD^+^ peripheral B cells and assigned animal-specific germline alleles using IgDiscover (*57*). In parallel, we sorted and sequenced IgG^+^ B cells from draining lymph nodes (post-immunization, Groups 1-2) and PBMCs (post-infection, Groups 1-3) at multiple timepoints, using the SONAR analysis pipeline (*58*) to trace the evolution of the bNAb lineages. In total, we analyzed 10,214,345 unique IgG and 2,172,207 unique IgM sequences from eight macaques to infer twelve high-confidence V3-glycan bNAb UCAs **(Fig. 3A)**.

**Fig. 3.**
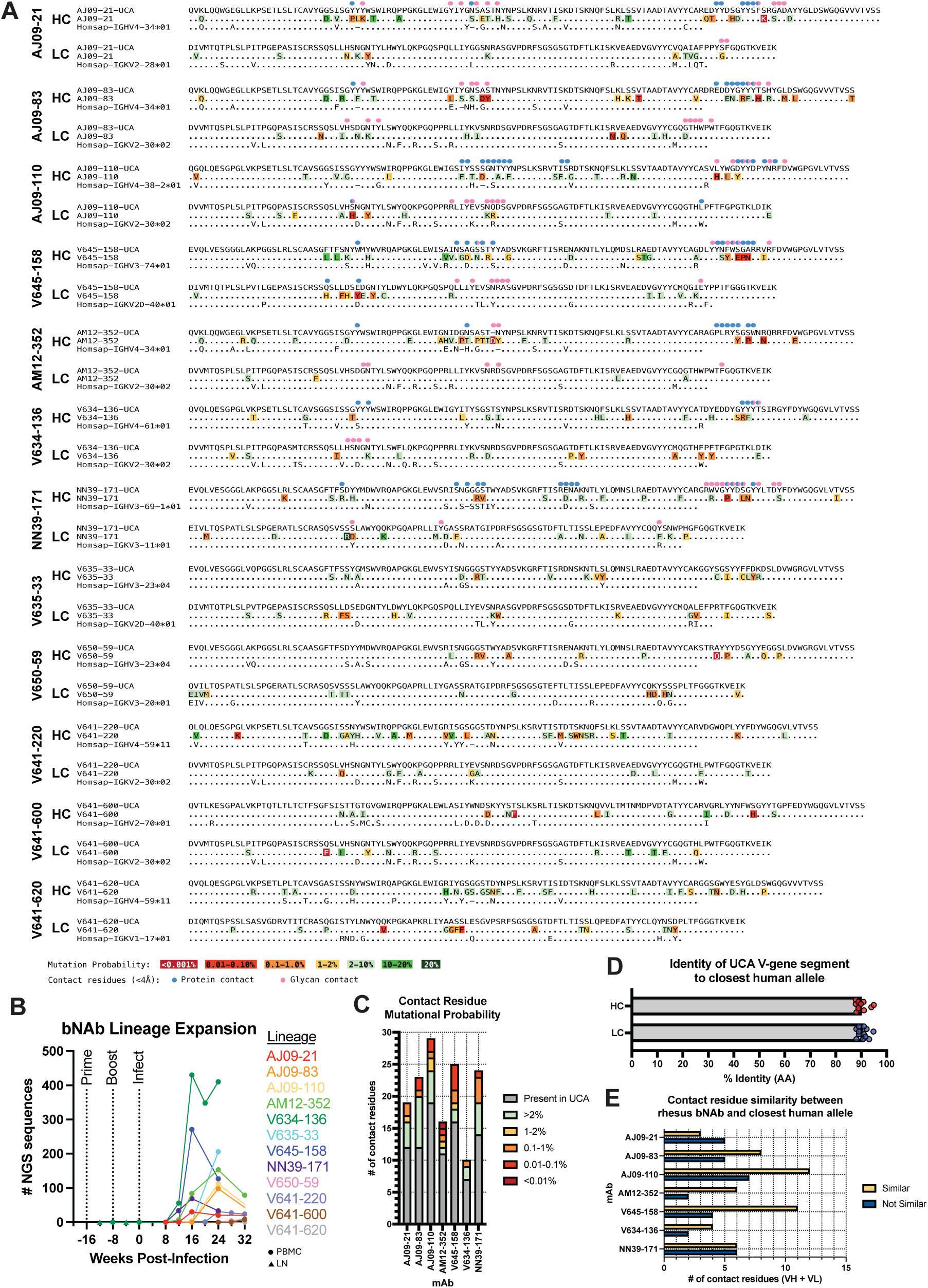
Characterization of inferred V3-glycan bNAb UCAs suggests similar antibodies may be elicitable in humans. **(A)** Alignments of heavy chain (HC) and light chain (LC) amino acid sequences of twelve inferred UCAs, their corresponding mature bNAbs, and closest human V_H_/V_K_ alleles are shown. Somatically mutated residues in the mature bNAb sequences are color-coded by estimated mutation probability, as determined by ARMADiLLO (*59*). Mature bNAb residues that contact amino acids within gp120 (defined as <4Å) are indicated by blue circles, whereas bNAb residues that contact glycans (defined as <4Å) are indicated by pink circles. **(B)** Kinetics of bNAb lineage stimulation and expansion, quantified as the number of sequences from bulk IgG^+^ B cells at each timepoint belonging to the indicated bNAb lineage. Circles indicate sequences were obtained from PBMCs, while triangles indicate sequences were obtained from lymph nodes. **(C)** Number of contact residues encoded by the UCA (grey) compared to those generated by somatic hypermutation for the indicated bNAbs. The somatically mutated contact residues are color-coded as in **(A)**. **(D)** Percent amino acid sequence identity between each UCA V_H_/V_K_ gene segment and the closest human allele. **(E)** Number of contact residues that are biochemically similar in the closest human V_H_ and V_K_ alleles. HC, heavy chain; LC, light chain; NGS, next-generation sequencing; PBMC, peripheral blood mononuclear cell; LN, lymph node; UCA, unmutated common ancestor; AA, amino acid.

To determine when these lineages were initiated, we queried the BCR sequencing dataset to identify the earliest timepoint at which lineage members could be detected in each animal. We did not find any lineage members in the draining lymph node or PBMC compartments of vaccinated animals (Groups 1 and 2) prior to infection with SHIV.5MUT, suggesting that the immunizations did not prime these bNAb lineages. In contrast, we observed early members within 12-24 weeks of infection in all groups **(Fig. 3B)**, indicating rapid and efficient priming by SHIV.5MUT or a derivative. This timing was consistent with the first detection of plasma neutralization breadth, typically around 20-32 weeks post-infection **(fig. S2)**.

Given the rapidity with which these lineages developed neutralization breadth **(fig. S2)** and their comparatively low rates of somatic mutation **(Fig. 1I)**, we hypothesized that their maturation pathways might be relatively simple. Indeed, we found that most structurally identified contact residues were UCA-encoded rather than the product of somatic hypermutation (**Fig. 3A** and **C**). We also analyzed the probability of occurrence of each somatic mutation using the ARMADiLLO computational pipeline, which takes into account both the number of nucleotide substitutions required to make a nonsynonymous change as well as the predilection of activation-induced cytidine deaminase (AID) to target certain sequence motifs (*59*). Of the non-UCA-encoded contact residues, half were predicted to be “probable” mutations, suggesting that they may be relatively easy to induce by vaccination (**Fig. 3A** and **C**).

To assess the translational relevance of our findings to humans, we searched the orgdb database (*60*) for human V_H_ and V_K_ alleles that had the highest amino acid identity to the rhesus bNAb UCAs. This analysis identified closely related human counterparts for all immunoglobulin gene segments used by the rhesus bNAb UCAs, with average amino acid identities of 90% (V_H_) and 91% (V_K_) (**Fig. 3A** and **D**). Interestingly, these included gene segments used by canonical human V3-glycan bNAbs: IGHV4-34 (used by PCDN-38A) and IGHV4-59 (used by PGT121) **(Fig. 3A)** (*15, 18, 41*). Additionally, of the 81 V-encoded bNAb contact residues, 50 were biochemically similar **(Fig. 3E)** and 41 were identical **(fig. S9)** to the closest human allele. Together, these data suggest that humans should be able to generate V3-glycan bNAbs akin to those described here, with similar developmental trajectories to breadth and potency.

### Env evolution in the V1 loop precedes bNAb induction

To elucidate the molecular events responsible for efficient V3-glycan bNAb priming and boosting in SHIV.5MUT-infected macaques, we conducted longitudinal single-genome sequencing (SGS) of circulating plasma viral RNA (*61, 62*). Plasma virus has a lifespan of ∼1 hour, and the cells producing it have a lifespan of ∼1 day, making the quasispecies composition a sensitive and dynamic indicator of viral fitness and selection pressures, including those exerted by neutralizing antibodies (*48, 63–66*). SGS provides a direct and proportional representation of target viral sequences in a plasma sample and has the advantage of maintaining sequence linkage across complete gp160 *env* genes, thereby enabling antibody pressure to be mapped across Env (*62*). We thus compared the patterns of Env sequence evolution in macaques infected with SHIV.5MUT versus the parental SHIV.BG505.N332 **(Fig. 1B)**, which failed to induce V3-glycan bNAbs.

To identify the SHIV.5MUT Env variants responsible for priming V3-glycan bNAb precursors, we serially analyzed the plasma virus quasispecies composition starting shortly after infection. Coincident with the rapid development of high-titer autologous neutralizing antibodies in the plasma **(Fig. 1C)**, we observed striking selection within the V1 loop of Env in all macaques infected with SHIV.5MUT, but in none of those infected with SHIV.BG505.N332 **(Fig. 4A, fig. S10)**. Mutations were detected as early as week 4 post-infection and led to complete replacement of the infecting SHIV.5MUT V1 loop sequences by week 16 in all animals **(Fig. 4A, fig. S10)**. Shared patterns of mutations were observed across SHIV.5MUT-infected animals, including a P136S variant that was present in 21/22 macaques and Δ135-138 / Δ136-139 (del4) deletion variants that encode identical V1 amino acid sequences and were present in 19/22 macaques **(Fig. 4B, fig. S10)**. Other mutations shared by multiple macaques included Δ137-139 (del3), Δ132-139 (del8), and R143G/K **(Fig. 4B, fig. S10)**.

**Fig. 4.**
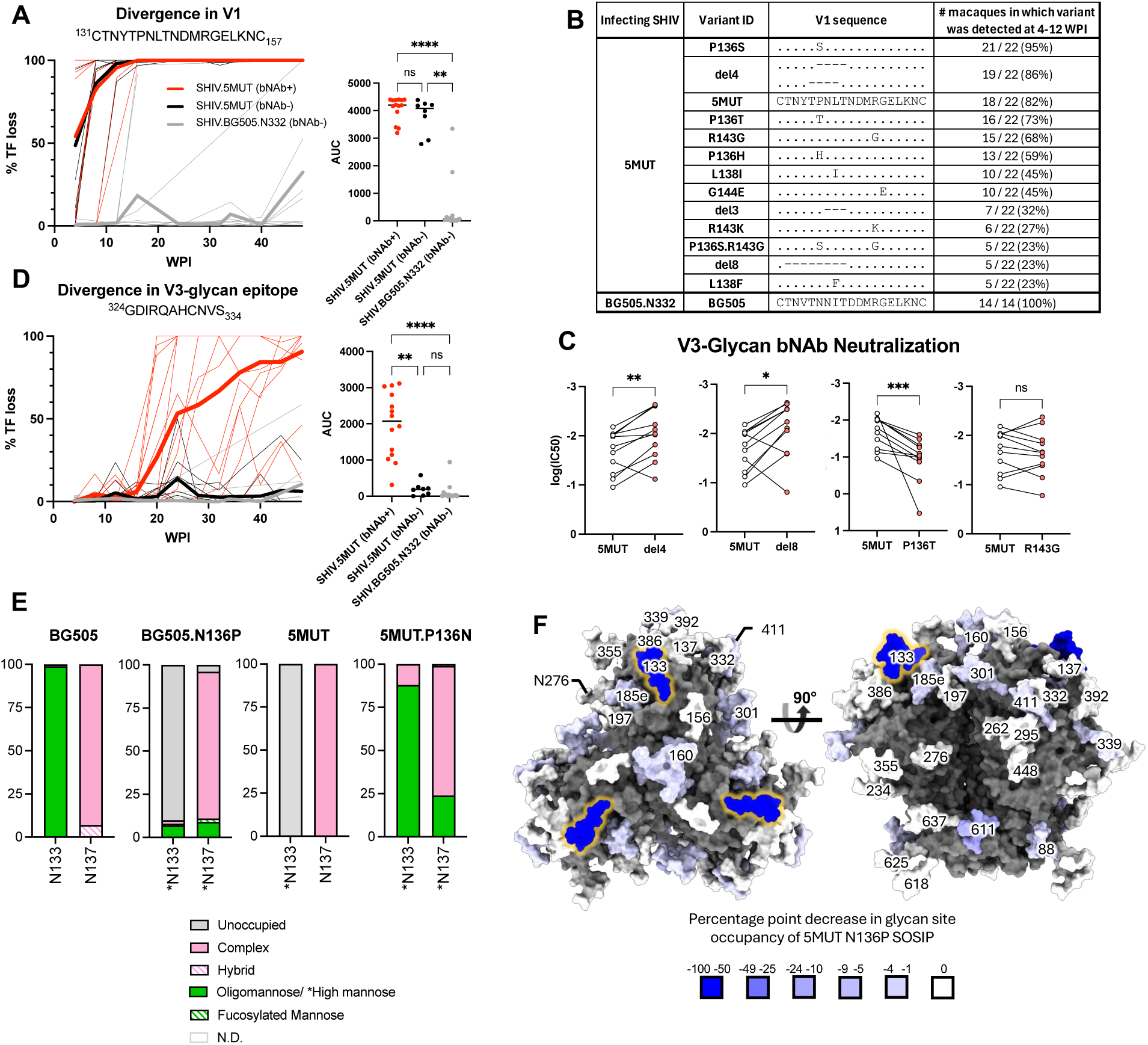
Selection in the V1 region precedes selection in the V3 region in SHIV.5MUT-infected macaques. (**A, D)** Left: Fraction of amino acid variants in the V1 **(A)** or V3 **(D)** regions of Env plotted over time for SHIV.5MUT-infected macaques that did (red) versus did not (black) develop bNAbs, as well as for SHIV.BG505.N332-infected macaques (grey). Thin lines indicate data from individual animals, and thick lines represent group averages. Right: Area under the curve (AUC) for each group. Kruskal-Wallis test with Dunn’s correction for multiple comparisons was used. Not significant (ns) P > 0.05; *P < 0.05; **P < 0.01; ***P < 0.001; ****P < 0.0001. **(B)** V1 sequence variants appearing in at least 20% of macaques between 4-12 weeks post-infection are shown. **(C)** Neutralization potency of representative V3-glycan bNAbs against common V1 variants, shown as log(IC_50_). Wilcoxon matched-pairs signed-rank test was used. Not significant (ns) P > 0.05; *P < 0.05; **P < 0.01; ***P < 0.001. **(E)** Mass spectroscopy-based site-specific glycan analysis of MD39-stabilized 5MUT and BG505.N332 SOSIP trimers. Glycan compositions are grouped into their corresponding categories, with complex-type glycans displayed in pink, oligomannose in green, and unoccupied in gray. For sites in which the site-specific data was deduced by employing glycosidase digestion by Endo H followed by PNGase F in O18 water (indicated by asterisks), the peptide modifications that correspond to distinct glycan types are grouped into high mannose in green, fucosylated mannose in hatched green, complex in pink, and unoccupied in gray. Glycan sites that could not be determined are denoted as “N.D.” **(F)** A glycosylated model of 5MUT SOSIP was generated using the 5MUT-3fill structure, and a heat map of the percentage point change in glycan site occupancy was generated and plotted on the Env model. The N185h glycan is not modelled due to a lack of structural determination of the underlying protein sequence covering this region. Man_5_GlcNAc_2_ glycans were modelled at all other PNGS within the structure using GlycoShape Re-Glyco (*101*) and ChimeraX (*102*). AUC, area under the curve.

Hypothesizing that a subset of these V1 variants primed the V3-glycan bNAb precursors *in vivo*, we generated representatives as infectious Envs and assessed neutralization sensitivity to mature V3-glycan bNAbs, reasoning that enhanced sensitivity may indicate greater epitope accessibility and thus greater priming potential. Indeed, variants with shortened V1 loops such as del4 and del8 were neutralized with significantly greater potency than the parental SHIV.5MUT. In contrast, the P136T variant was more resistant to neutralization and the R143G variant showed no difference **(Fig. 4C)**. Consistent with the hypothesis that early, shortened-V1 variants like del4 and del8 primed the bNAb lineages, we observed sequence signatures of bNAb escape at the V3-glycan epitope and accrual of plasma breadth shortly after their emergence. Specifically, we observed mutations in the ^324^GDIR_327_ motif and at H330, as well as deletion or shifting of the N332 glycan, starting as early as week 16 post-infection in SHIV.5MUT-infected animals that went on to develop plasma breadth **(Fig. 4D)**. In contrast, SHIV.BG505.N332-infected macaques showed almost no V1 and V3 selection throughout the first 48 weeks of infection (**Fig. 4A, B,** and **D, fig. S10)**.

The vastly different V1 selection patterns observed in SHIV.5MUT- versus SHIV.BG505.N332-infected macaques suggested that the four distinguishing residues in the 5MUT Env enhance its immunogenicity **(Fig. 1B)**. BG505 contains PNGS sequons within the V1 loop at positions N133 and N137 (*67, 68*), comprising part of the glycan shield thought to dampen the immunogenicity of HIV-1 Env (*69, 70*). While 5MUT retains both of these PNGS sequons (*24*), we hypothesized that the proline introduced at position 136 could compromise glycosylation at the adjacent N133 and/or N137 sites, as proline residues immediately before or after a PNGS sequon have been shown to reduce N-glycan occupancy (*31, 71, 72*). Glycan analysis by mass spectrometry confirmed that the N133 PNGS in 5MUT SOSIP was almost completely glycan-devoid, while the N137 PNGS remained fully occupied **(Fig. 4E-F, fig. S11-12)**. Similarly, introducing an N136P substitution in BG505 SOSIP substantially reduced glycan occupancy at position N133, but did not appreciably affect occupancy at position N137 **(Fig. 4E, fig. S11-12)**. Reduced V1 glycosylation would be expected to enhance 5MUT immunogenicity compared to the parental BG505 Env, inducing a targeted humoral response that accounts for the striking V1 selection that was observed. Indeed, most of the V1 variants shared among macaques eliminated the P136 residue by point mutation or deletion **(Fig. 4B, fig. S10)**, thereby restoring V1 glycosylation at N133 and escaping the early plasma neutralizing response.

### SHIV.5MUT induces V3-glycan bNAbs via a two-step mechanism

Given that V1 selection preceded bNAb lineage initiation and subsequent V3 selection in SHIV.5MUT-infected animals, we hypothesized that bNAb elicitation occurs by a two-step process. Specifically, we propose a model whereby an initial wave of V1-directed antibodies exerts selective pressure on the V1-glycan-deficient 5MUT Env, prompting escape mutants that shorten the V1 loop and expose the underlying V3-glycan epitope, which in turn prime V3-glycan bNAb precursor B cells **(Fig. 5A)**. To test this model, we mapped the earliest plasma response against viruses expressing 5MUT-derived V1 mutant Envs including del4, del8, and P136T. Consistently, we observed a significant reduction in neutralization activity against these variants compared to the parental SHIV.5MUT, confirming that much of the early neutralizing response is V1-directed in all SHIV.5MUT-infected macaques **(Fig. 5B)**. To corroborate this conclusion, we isolated antibodies responsible for driving the selection of these V1 escape mutants *in vivo* by sorting single 5MUT^++^del4^-^ or 5MUT^++^BG505^-^ IgG^+^ B cells from PBMCs, amplifying their antibody genes, and synthesizing representative mAbs. This approach yielded V1-specific antibody lineages from two different macaques that recapitulated the early plasma activity, potently neutralizing SHIV.5MUT but not the V1 escape variants **(Fig. 5B)**. We determined a single-particle cryo-EM structure of one V1-directed Fab, NN39-25, in complex with the 5MUT SOSIP **(Fig. 5C)**. As expected, NN39-25 interacted with multiple V1 residues including ^134^YTPNLTN_140_, which contains the four 5MUT substitutions and would therefore be exposed by the reduced N-glycan occupancy at position N133_gp120_ **(Fig. 5D)**. Structural overlay with NN39-171, a bNAb isolated from the same animal, revealed that while NN39-25 primarily interacts with the V1 loop, NN39-171 exhibits a shifted binding pose that favors interactions with the ^324^GDIR_327_ motif and the N332_gp120_ glycan **(fig. S13)**.

**Fig. 5.**
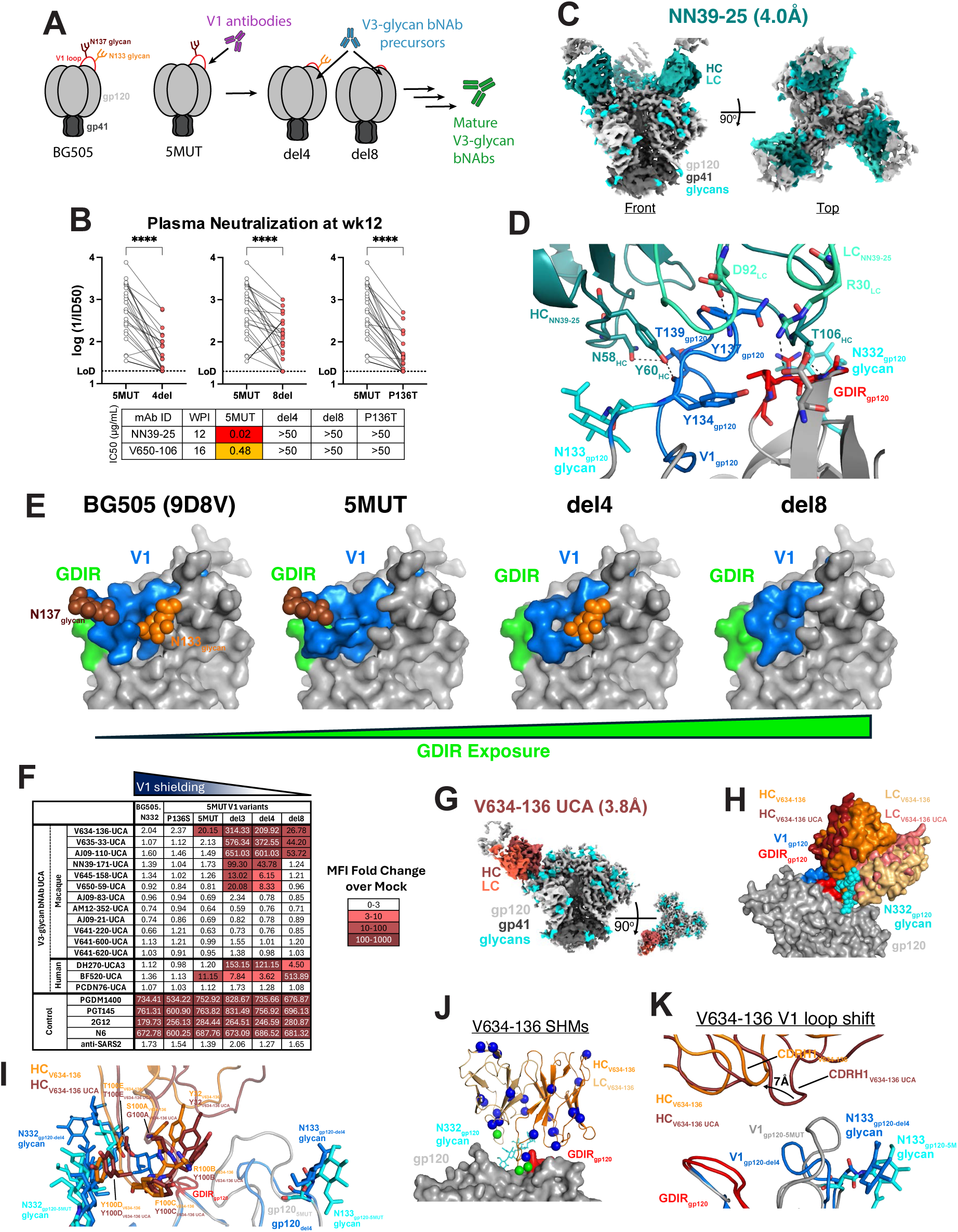
SHIV.5MUT elicits V3-glycan bNAbs via a two-step process. **(A)** Schematic model of SHIV.5MUT-mediated V3-glycan bNAb elicitation. **(B)** Top, neutralization of the indicated V1 variants by plasma from SHIV.5MUT-infected macaques at week 12 post-infection, with titers represented as log(1/ID_50_). Wilcoxon matched-pairs signed-rank test was used. ****P < 0.0001. Bottom, neutralization activity of two V1-directed monoclonal antibodies, represented as IC_50_ in µg/mL. **(C)** Cryo-EM map of V1-directed antibody NN39-25 in complex with 5MUT-3fill SOSIP. **(D)** Potential interactions between NN39-25 and the GDIR motif (red) and V1 loop (blue) of the 5MUT gp120. **(E)** Cryo-EM structures of BG505 (PDB: 9D8V), 5MUT-3fill, del4-3fill, and del8-3fill, highlighting structural differences in GDIR motif exposure as influenced by V1 loop length and glycosylation. **(F)** UCA binding to cell-surface expressed Envs. mRNA encoding the indicated Envs was transfected into 293F cells and binding was assessed by flow cytometry. Binding is quantified as fold-change in mean fluorescence intensity (MFI) over mock-transfected cells, and data points with a >3-fold increase are colored as indicated. The results shown are the average of two independent experiments. (**G)** Cryo-EM map of V634-136-UCA in complex with del4-3fill SOSIP. **(H-I)** Structural overlays of V634-136 UCA and mature bNAb revealing nearly identical approach angles (**H**) and CDRH3 architectures **(I)**. **(J)** Somatic hypermutations (SHMs) acquired by V634-136-UCA are mapped onto the coordinates. Green spheres represent residues on V634-136-UCA that form contacts with gp120, while blue spheres represent residues outside of the gp120 binding interface. **(K)** Structural overlay of V634-136-UCA and the mature V634-136 bNAb revealing a shift in CDRH1 to accommodate the longer V1 loop in 5MUT. UCA, unmutated common ancestor; MFI, mean fluorescence intensity; PNGS, potential N-linked glycosylation site; SHMs, somatic hypermutations.

Having shown that induction of V1-directed antibodies like NN39-25 led to the selection of V1 variants *in vivo*, we next characterized the variant Envs themselves. We focused on the del4 and del8 variants because viruses bearing these Envs exhibited enhanced neutralization sensitivity to V3-glycan bNAbs **(Fig. 4C)**, suggesting they may provide increased epitope accessibility and thus enable UCA binding. Additionally, the V1 loop directly overlies the V3-glycan epitope in the pre-fusion Env conformation (*73, 74*), lending credence to the hypothesis that shortening V1 increases epitope exposure. To test this hypothesis, we determined single-particle cryo-EM structures of unliganded SOSIP Env versions of 5MUT, del4, and del8 **(Fig. 5E, Supplementary Table 2)**. As expected, the smaller footprint of the del4 and del8 V1 loops resulted in greater exposure of the V3-glycan epitope, including the conserved ^324^GDIR_327_ motif, compared to BG505 and 5MUT. Reduced V1 glycosylation in 5MUT also likely increased accessibility of the V3-glycan epitope compared to BG505, as evidenced by the site-specific mass spectrometry analysis **(Fig. 4E-F, fig. S11-12)**, although this could not be demonstrated in our cryo-EM structures as the N133_gp120_ and N137_gp120_ glycans were not well-ordered and coordinates could not be assigned past the initial GlcNAc residue in each case **(fig. S14)**. Together, these data support a model in which hypoglycosylated SHIV.5MUT induces an early wave of V1-directed antibodies that select for Env V1 escape mutations, including deletions. Such variants, typified by del4 and del8, expose the underlying V3-glycan epitope and likely prime bNAb precursors.

### Identification of early Env variants that primed V3-glycan bNAb lineages

To identify Env mutants capable of engaging diverse V3-glycan bNAb UCAs, we interrogated the Env-antibody coevolution datasets. Based on sequencing **(Fig. 4B)**, structural **(Fig. 5E),** and functional **(Fig. 4C)** data, we identified del4 and del8 as the most promising candidates and generated an mRNA-encoded version of each, employing the same stabilization strategy used for RC1 and 11MUTB **(fig. S1A)**. For comparison, we generated mRNA constructs encoding other V1 variants (del3 and 5MUT.P136S) as well as controls (5MUT and BG505.N332). These Env variants were expressed on the surface of 293F cells and assessed for their ability to bind human **(Supplementary Table 3)** and macaque V3-glycan bNAb UCAs **(Fig. 5F)**. All constructs expressed high levels of well-formed Env trimers on the cell surface, as evidenced by robust binding of bNAbs PGT145 and PGDM1400, which recognize quaternary epitopes (*75, 76*) **(Fig. 5F)**. Consistent with their full-length, glycosylated V1 loops, neither BG505.N332 nor 5MUT.P136S exhibited detectable binding (>3-fold over background) to any UCA tested, and V1-glycan-deficient 5MUT showed only low-level binding to 2 of 15 UCAs. In contrast, Env variants with short V1 loops and lacking glycans (del3, del4, and del8) bound up to 8 of 15 UCAs and did so much more robustly than did 5MUT **(Fig. 5F)**. Thus, these V1 mutant Envs could have primed V3-glycan bNAb precursors *in vivo*.

To visualize the molecular interactions of an authentic UCA with one of these novel priming immunogens, we determined a single-particle cryo-EM structure of V634-136-UCA Fab in complex with the del4 SOSIP **(Fig. 5G, Supplementary Table 2)**. Interestingly, V634-136-UCA targets the V3-glycan epitope with a near-identical angle of approach **(Fig. 5H)** and nearly all the same interactions **(Fig. 5I)** as the mature V634-136 bNAb, which is consistent with the fact that it acquired only three somatic mutations in its gp120 binding interface over the course of its development **(Fig. 3A**, **Fig. 5J)**. The main difference between the UCA and mature bNAb structures is that the CDRH1 loop of V634-136 was shifted away from gp120 by ∼7Å compared to that of V634-136-UCA, which likely serves to accommodate the longer V1 loop in 5MUT compared to del4 **(Fig. 5K)**.

Six UCAs, including AM12-352-UCA and AJ09-83-UCA, did not exhibit detectable binding to any of the SHIV.5MUT V1 escape variants. To identify Env variants that could have primed these lineages *in vivo*, we produced intermediate mAbs from various stages of lineage maturation by pairing heavy and light chains inferred from the longitudinal BCR sequence dataset **(fig. S8, fig. S15A)**. Interestingly, we detected robust Env binding to early inferred ancestor (iA) mAbs from both lineages: AJ09-83-iA2 bound del3 and del4, and AM12-352-iA3 bound 5MUT, del3, and del4, and the binding magnitudes increased as more-mature lineage members were tested **(fig. S15B)**. Although no binding of del3, del4, and 5MUT to AM12-352-UCA or AJ09-83-UCA was detected in the flow-based assay, this does not exclude the possibility that these Envs stimulated these lineages *in vivo*. The binding threshold for BCR activation in a germinal center remains poorly understood, and given the avidity effects afforded by the follicular dendritic cell network (*77*), it is possible that the detection limit of the flow-based assay is too low to capture physiologically relevant low-level binding. Additionally, given the minimal Env diversity in the circulating quasispecies around the time these lineages were initiated (week 4-12) **(fig. S10)**, it is unlikely that we overlooked other plausible priming Envs variants.

### Env-antibody coevolution guides immunogen design

In humans, Env-bNAb coevolution has been shown to be an arms race in which bNAb lineage members exert pressure on the contemporaneous viral quasispecies, leading to the selection of viral escape variants that in turn select for maturing bNAb lineage members capable of binding to them (*19, 78, 79*). Over multiple rounds of antibody recognition and virus escape, the viral quasispecies samples multiple variant forms that impair antibody recognition, often including amino acids that are common in globally circulating HIV-1 strains. As bNAb lineages mature to accommodate these quasispecies mutations, they incrementally acquire heterologous neutralization breadth (*19, 78, 79*). Having observed such developmental trajectories in several longitudinal human studies, we explored here whether this concept could be applied to inform the design of boost immunogens for SHIV.5MUT-based vaccines.

We first analyzed Env sites under selection in SHIV.5MUT- and SHIV.BG505.N332-infected macaques using LASSIE (*80*), defined as transmitted/founder (TF) loss in ≥80% of quasispecies sequences at any single timepoint within the first year of infection **(Fig. 6A)**. This analysis identified a total of 78 unique sites under selection, with a median of seven sites (range 0-26) per macaque **(fig. S16A)**. SHIV.5MUT-infected animals that developed bNAbs had a significantly greater number of selected Env sites than those that did not develop bNAbs (p=0.0006, Wilcoxon rank-sum test) and SHIV.BG505.N332-infected macaques (p=0.00002, Wilcoxon rank-sum test), consistent with their higher viral loads and stronger antigen drive (**fig. S17A-B**). Fifty-one of the 78 LASSIE-identified sites were found to be under selection in more than one animal, with almost all macaques showing selection signatures in known strain-specific, immunodominant epitopes on the BG505 Env including the N241/N289 glycan hole and the C3-V4-V5 region **(fig. S16B)**.

**Fig. 6.**
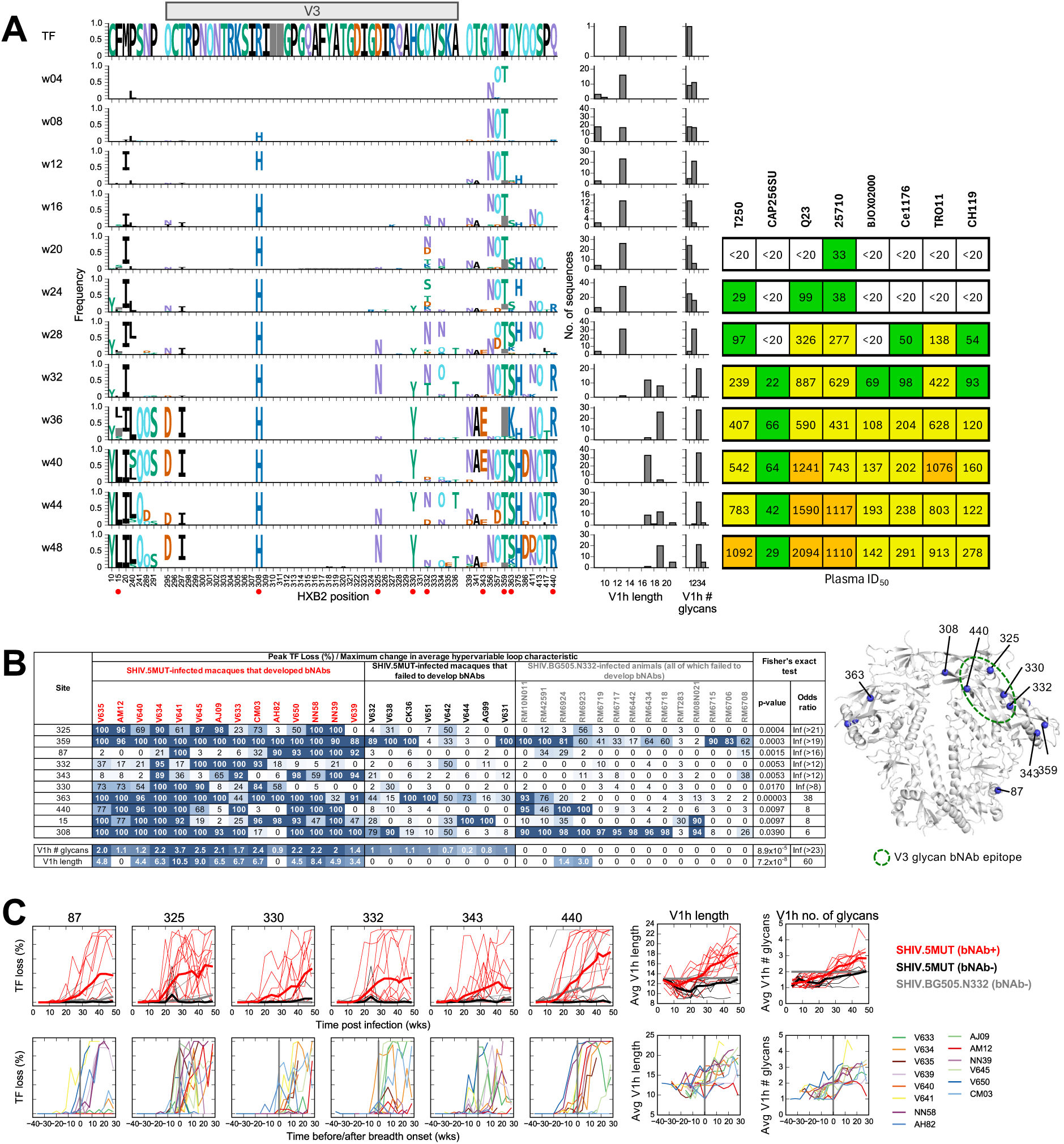
Env-antibody coevolution analysis reveals key Env mutations enriched in SHIV.5MUT macaques that develop bNAbs. **(A)** Longitudinal patterns of Env mutations and development of plasma neutralization breadth. Left: Mutations in the quasispecies of animal V634 are depicted as logos, where the height of the letter is proportional to its frequency in the Env sequences from the corresponding timepoint. Cyan colored “O” indicates asparagine (N) embedded in an N-linked glycosylation motif (NXS or NXT, where X can be any amino acid except proline). Numbers at the bottom indicate residue positions (HXB2 numbering). Amino acids that match the infecting SHIV.5MUT strain (designated transmitted founder or TF) are hidden. Env positions shown include residues 295-336, which comprise the entire V3 loop and adjacent PNGS, as well as all sites that were deemed to be under selection in the evolving quasispecies by meeting a pre-determined ≥80% TF loss criteria at any timepoint within the first year of infection, as defined by LASSIE (*80*). Red dots indicate sites under selection that are enriched in animals that developed bNAbs. Middle: Bar plots illustrating the fraction of sequences at each timepoint with a given hypervariable V1 (V1h) loop length or number of glycans. The SHIV.5MUT TF has a V1h length of 13 AA and one V1h glycan, and an increase in either is associated with neutralization resistance. Right: Plasma neutralization over time against eight heterologous viruses, expressed as reciprocal ID_50_. **(B)** Left: LASSIE-selected sites that are significantly enriched in SHIV.5MUT-infected macaques that developed bNAbs as compared to SHIV.5MUT- and SHIV.BG505.N332-infected macaques that did not develop bNAbs, as determined by Fisher’s exact test (p < 0.05, uncorrected). Numbers for each of the ten sites indicate percentage of peak TF loss, and only those sites that met a ≥80% peak TF loss cutoff within a given animal were counted. The maximum increase in average number of V1h glycans (V1h # glycans) and V1h length are also shown. Right: Structural mapping of selected sites that were enriched in SHIV.5MUT-infected macaques that developed bNAbs onto the BG505 Env (PDB ID: 9EHL). The green ellipse demarcates the approximate location of the V3-glycan bNAb epitope. **(C)** Top: Mutation frequency (% TF loss) at six sites enriched in SHIV.5MUT-infected macaques that developed bNAbs (indicated at top) and average V1h loop length and number of PNGS over 48 weeks of infection. Red and black curves denote SHIV.5MUT-infected macaques that did versus did not develop bNAbs, respectively, while grey curves indicate SHIV.BG505.N332-infected animals (all of which failed to develop bNAbs). Thin lines indicate data from individual animals and thick lines represent group averages. Bottom: Mutation frequency (% TF loss) in SHIV.5MUT-infected macaques that developed bNAbs over time, with the timepoint at which plasma breadth is first detected (defined as at least 1 heterologous virus with reciprocal ID_50_ > 20) set to zero weeks (wks). Different colors indicate different macaques. Plots for the remaining four selected sites (15, 308, 359, 363) are shown in **fig. S16C**. When two V1 glycan PNGS sequons were separated by a single proline residue (as in 5MUT), only one was counted as glycosylated based on glycan occupancy determined for 5MUT SOSIP.

To elucidate which mutations likely contributed to V3-glycan bNAb maturation, we next identified ten LASSIE-selected sites that were significantly enriched (p<0.05, Fisher’s exact test) in macaques that developed bNAbs compared to those that did not (combining SHIV.5MUT-infected macaques that did not develop bNAbs and all SHIV.BG505.N332-infected animals into a single dataset) (**Fig. 6B**). These ten sites were under selection in a median of 8 of the 14 animals that developed bNAbs (range: 5-14), as opposed to 0 of the 22 animals that failed to develop bNAbs (range: 0-11). Remarkably, these sites included residues 325, 330, and 332 within the V3-glycan epitope as well as residue 440, which is in close physical proximity to the ^324^GDIR_327_ motif **(Fig. 6B)** and has previously been identified as a recurrent and robust signature site for human V3-glycan bNAbs (*21*). Furthermore, we found that insertions and additions of PNGS sites in the hypervariable portion of the V1 region (V1h, comprising residues ^132^TNYTPNLTNDMRG_152_) were significantly more frequent in macaques that developed bNAbs compared to those that failed to develop bNAbs (**Fig. 6B)**.

Because neutralization breadth is acquired incrementally over the course of bNAb lineage development, Env mutations that are temporally correlated with its onset and enhancement may be important for bNAb maturation. Of the ten selected sites enriched in macaques that developed bNAbs, mutations at residues 87, 325, 330, 332, 343, and 440 arose either immediately prior to or shortly after the onset of plasma neutralization breadth. Similarly, increases in V1 length and number of PNGS sequons were observed around the time of breadth acquisition (**Fig. 6C** and **fig****. S18-31**). In contrast, mutations at the four remaining sites (15, 308, 359, and 363) were not temporally associated with plasma breadth **(fig. S16C)**, though they may still be relevant to bNAb development as they were present in the host quasispecies as the bNAb lineages matured (**fig. S18-31**). For example, site 15 is within the signal peptide and could be relevant to V3-glycan bNAb development since signal peptide modifications have been shown to impact Env glycan occupancy, which could affect antibody recognition (*81–83*).

We hypothesized that among the ten selected sites and V1h modifications, there may be a set of key sites that restrict heterologous neutralization until relevant variants are sampled by the viral quasispecies. To test this hypothesis, we first determined which of the ten selected sites differed between 5MUT and each of the eight heterologous viruses in our screening panel **(Fig. 7A)**. Key sites were defined as the subset of positions that differed in the heterologous virus and were under selection in the corresponding animal. We reasoned that amino acid variants at these sites would need to be recognized by the maturing antibody lineage in order to gain neutralization activity against the corresponding heterologous virus. Two of the ten selected sites, the PNGS at position 332 (referred to as O332) and the isoleucine at position 359 (I359), were invariant in our heterologous panel and thus could not be evaluated, though they may still be relevant to bNAb recognition.

**Fig. 7.**
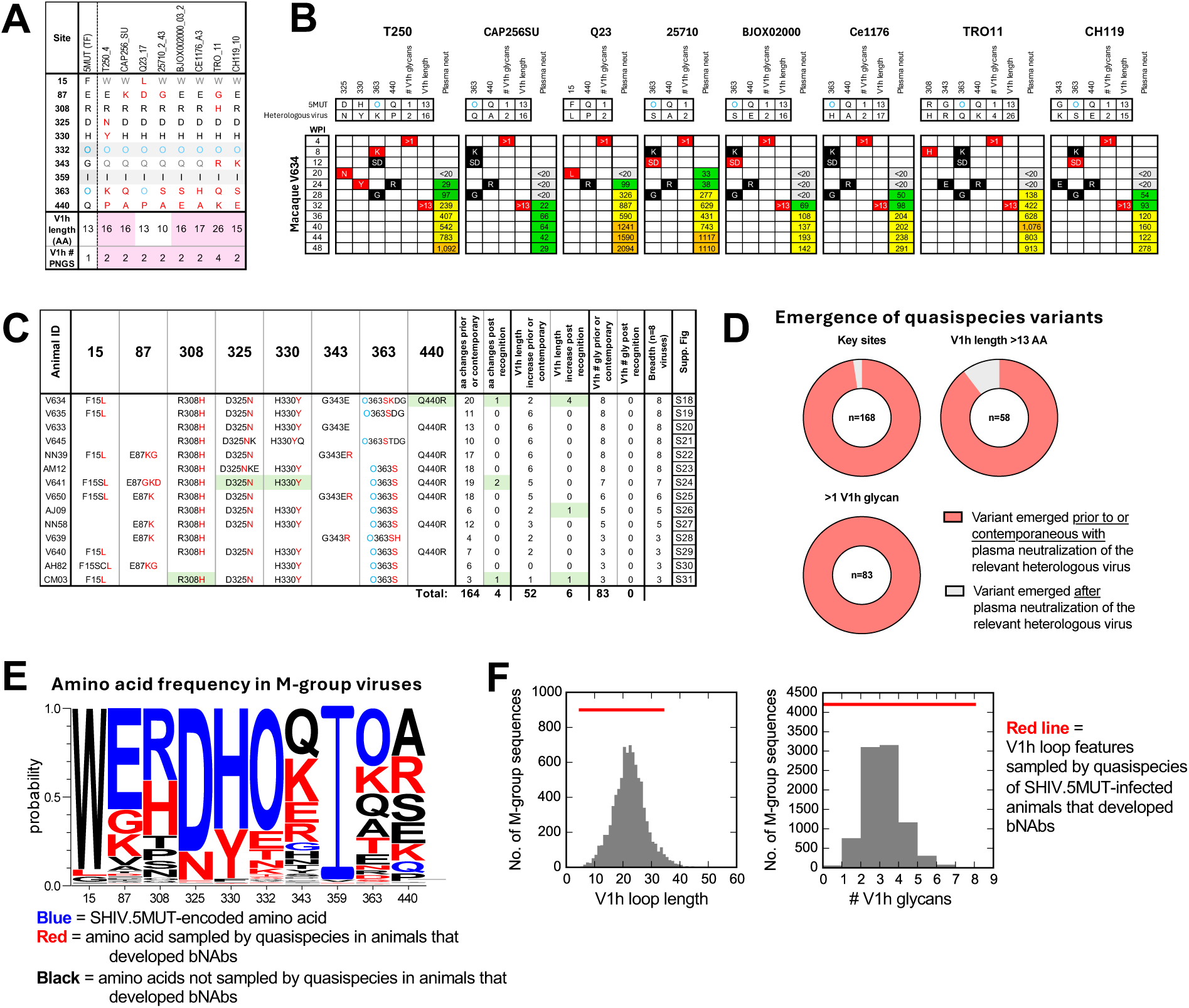
Quasispecies sampling of M-group diversity at key sites precedes or coincides with the acquisition of plasma neutralization breadth. **(A)** Amino acids at the ten Env sites that were preferentially under selection in SHIV.5MUT-infected macaques that developed bNAbs as well as V1h length and number of PNGS compared to the eight heterologous viruses in the neutralization test panel. Sites 332 and 359 (highlighted in grey) were identical in all heterologous viruses tested and matched SHIV.5MUT, and thus could not be evaluated using this panel. Residues in heterologous Envs that match versus do not match those of SHIV.5MUT are shown in black and red, respectively, and potential N-linked glycosylation sites (PNGS) are shown as a blue “O”. W15 and Q343 (shown in grey) match the global HIV-1 consensus but were never sampled in the quasispecies of any SHIV.5MUT-infected macaque that developed bNAbs and thus were not included in subsequent analyses. V1h loops that are longer or have more PNGS than that of SHIV.5MUT are highlighted in pink. **(B)** Consistent emergence of key mutations in the quasispecies of macaques that developed bNAbs prior to their ability to neutralize heterologous viruses containing these residues. The subset of the ten bNAb-associated sites that are under selection in macaque V634 and differ between each heterologous virus (indicated at top) and SHIV.5MUT is shown. For each site, both the time point at which mutations are first detected in the evolving quasispecies and the sampled residues are indicated. Red boxes indicate the sampled residue matches that of the heterologous virus, black boxes indicate the sampled residue does not match the heterologous virus, and blue font indicates a potential N-linked glycosylation site (PNGS). For heterologous viruses that have longer V1h loops or more PNGS than SHIV.5MUT, the timepoint at which the evolving quasispecies first showed increased V1h length or number of glycans is noted. Serial plasma neutralization activity against each heterologous virus is listed as reciprocal ID_50_. In general, mutations at all relevant key sites for a given heterologous virus as well as increases in V1h length and number of glycans are detected prior to or concurrent with acquisition of plasma neutralization activity against that virus. WPI, weeks post-infection. **(C)** Summary of the analysis in **(B)** but applied to all 14 macaques that developed bNAbs. Mutations are colored as in **(B)** and are listed in the following format: SHIV.5MUT TF AA / position / AA(s) sampled by the quasispecies in the indicated macaque. For each animal, the number of times that mutations emerged at the relevant sites prior to or concurrent with the acquisition of plasma neutralization activity against the corresponding virus were counted. Similarly, instances in which the quasispecies sampled increased V1h length or number of PNGS prior to or concurrent with acquisition of neutralization activity against the relevant virus were counted. The few instances in which mutations or V1 features were detected after acquisition of neutralization activity are highlighted in green. Refer to **figs. S18-31** for an in-depth analysis of each animal. **(D)** Summary of results in **(C)**. Pie charts depict the frequency with which key quasispecies variants (containing mutations at key sites, elongated V1h loops, or additional V1h glycans) emerged prior to or contemporaneous with acquisition of plasma neutralization against the relevant heterologous viruses. Numbers in the center represent the total instances of the relevant variants being detected across all 14 SHIV.5MUT-infected macaques that developed bNAbs and all eight heterologous viruses. **(E)** Amino acid frequency in global M-group viruses at the ten key sites enriched in SHIV.5MUT-infected macaques that developed bNAbs. Blue font indicates AAs encoded by SHIV.5MUT, red font indicates AAs that were sampled by the quasispecies of macaques that developed bNAbs, and black font indicates AAs that were not sampled. **(F)** Distribution of global M-group V1h loop lengths (left) and number of glycans (right). Horizontal red lines show the range of V1h length or number of glycans sampled across all SHIV.5MUT-infected macaques that developed bNAbs cumulatively. When two V1 glycan PNGS sequons were separated by a single proline residue (as in 5MUT), only one was counted as glycosylated based on glycan occupancy determined for 5MUT SOSIP.

We then assessed the time at which mutations emerged at key sites in each animal and compared it to when neutralization breadth was first detected against the corresponding heterologous virus **(Fig. 7B-C, fig. S18-31)**. Remarkably, in 164 of 168 (98%) cases across all animals and pseudoviruses, mutations at the relevant key sites were observed prior to or concurrent with the acquisition of plasma neutralization against the respective heterologous virus **(Fig. 7C-D)**. There were only four exceptions, and in each of these cases the neutralization potency prior to the emergence of the relevant mutations was very weak. Strikingly, the mutations away from the neutralization-sensitive 5MUT-encoded amino acid in the quasispecies often directly yielded the more resistant amino acid encoded by the heterologous virus, lending further credence to the notion that accommodation of these specific residues by the maturing bNAb lineages contributes to the acquisition of breadth **(Fig. 7A-C, fig. S18-31)**. Indeed, much of the heterologous M-group diversity at the ten key sites was captured by the quasispecies of SHIV.5MUT-infected animals that developed bNAbs, particularly at residues within the V3-glycan epitope (325, 330, and 332) **(Fig. 7E)**. Additionally, in 52 of 58 (90%) instances, the host quasispecies sampled V1h insertions before heterologous plasma neutralization activity was observed **(Fig. 7C-D)**. Finally, while the 5MUT Env has only one V1h glycan **(Fig. 4E-F)**, all heterologous viruses in our screening panel have two or more **(Fig. 7A)**. Consistently, in 83 of 83 (100%) cases, V1h PNGS additions were sampled in the quasispecies prior to the acquisition of neutralization breadth **(Fig. 7C-D)**. In fact, V1 loop length and number of PNGS continued to increase over time in macaques that developed bNAbs but not in those that failed to develop bNAbs **(Fig. 6C)**, thereby sampling V1 features that are represented in an even greater fraction of globally circulating viruses **(Fig. 7F)** and possibly promoting even greater breadth.

Overall, emergence of mutations at the key sites was gradual and plasma-level recognition of heterologous viruses was almost never detectable until mutations at all relevant key sites were observed **(Fig 7C-D**, **fig. S18-31)**. These results provide evidence that viral acquisition of key resistance mutations in response to a developing bNAb lineage *in vivo* provides critical exposure to viral variants that positively select for antibodies within the lineage, which in turn can tolerate diversity at these sites. However, although required, viral acquisition of key resistance mutations was not always sufficient for breadth development. Indeed, there were several instances in which a macaque did not gain plasma neutralization activity against a certain heterologous virus despite the quasispecies having sampled mutations at all key sites (e.g. animal V640, **fig. S29)**. Nonetheless, our analysis of Env-bNAb coevolution in SHIV.5MUT-infected macaques revealed common mechanisms of priming and maturing a diverse set of V3-glycan bNAb lineages.

## Discussion

The development of effective HIV-1 prevention and cure strategies will likely require the induction of protective immune responses, including the consistent elicitation of high-titer bNAbs. A major roadblock to vaccine design efforts has been the lack of a tractable model of bNAb induction in outbred animals. To date, the only example of consistent bNAb elicitation has been fusion peptide-directed bNAbs in rodents (*84*) and macaques (*65*), although these were of low potency and, in the latter case, took over two years to develop. Here, we address this gap with SHIV.5MUT, which elicited V3-glycan bNAbs in 14 of 22 (64%) macaques within the first year of infection and thus represents the first reproducible model of rapid and potent bNAb induction in outbred animals.

Plasma neutralization breadth and potency were generally high in SHIV.5MUT-infected animals, with eight macaques achieving 75% breadth on an eight-virus screening panel and ID_50_ titers exceeding 1:1000 in seven of them **(Fig. 1D)**. Plasma breadth emerged rapidly and was first detected an average of 25 weeks post-infection (range: 20-40), which represents a much-accelerated timeframe compared to natural infection in humans, where appreciable plasma breadth often takes years to develop (*12, 85, 86*). Heterologous neutralization activity was directed to the V3-glycan epitope in all instances, exhibiting strong dependence on the N332 glycan and residues in the ^324^GDIR_327_ motif **(Fig. 1E)**. Importantly, bNAb elicitation was unrelated to prior vaccination with RC1 or 11MUTB but was instead associated with SHIV viral load **(fig. S17A)**, corroborating previous findings that high antigen load promotes bNAb development (*86–88*). Overall, SHIV.5MUT infection elicited plasma neutralization breadth with potency and speed unparalleled by any outbred animal model of HIV-1 bNAb induction.

bNAbs isolated from macaques with the greatest plasma activity exhibited neutralization breadth of up to 68% on a panel of 130 global HIV-1 strains representing all major M-group subtypes **(Fig. 1F, fig. S4)**. This breadth increased to 75% when tested against just those viruses that carry an N332 glycan **(fig. S4)**. Rhesus bNAbs demonstrated functional, structural, and immunogenetic diversity, much like their human counterparts. Indeed, SHIV.5MUT infection induced at least three distinct V3-glycan bNAb lineages in each of two macaques (AJ09 and V641). Importantly, these concurrent bNAb lineages targeted the epitope in different ways **(Fig. 1H**, **Fig. 2A-B)** and neutralized complementary sets of viruses **(fig. S4)**. While most rhesus bNAbs were strictly N332 glycan-dependent, representatives of the broadest lineage (AM12-BM5 and AM12-352) depended instead on the N301 glycan, thereby broadening their coverage to include many clade AE viruses that lack an N332 glycan **(fig. S5A-B)**. Notably, the recently described N332-independent bNAb 007 is similarly among the broadest and most potent human V3-glycan bNAbs (*89*). Such diversity in epitope recognition is desirable because it can afford protection against viral strains that are resistant to other classes of V3-glycan bNAbs. Additionally, rhesus V3-glycan bNAbs matured rapidly and did not require extensive somatic hypermutation or indels, making them favorable targets for vaccine elicitation. Together, the breadth, potency, and diversity of SHIV.5MUT-induced bNAbs rival the best human V3-glycan bNAbs and can thus serve as a benchmark for future vaccine trials.

The fact that SHIV.5MUT rapidly induced diverse bNAbs in 64% of macaques indicates that authentic V3-glycan bNAb precursors are more abundant than previously appreciated. This is in part because, unlike the V2-apex and CD4bs bNAb classes (*46–49*), we found no readily recognizable immunogenetic features that defined or limited V3-glycan bNAb precursors. Other investigators have developed rules to define putative precursors that are based on immunogenetic similarities with one particular UCA, including gene segment usage and CDRH3 length, and these rules often yield vanishingly small estimates of precursor frequency in the naïve rhesus and human repertoires (*31, 90*). In contrast, SHIV.5MUT infection consistently elicited highly diverse bNAbs with widely varied immunoglobulin gene usage and CDRH3 lengths ranging from 14-25 amino acids, suggesting that current precursor definitions for V3-glycan bNAbs are overly restrictive **(Fig. 1I)**. In our study, we traced all twelve SHIV.5MUT-elicited lineages to their respective UCAs, thereby quadrupling the number of authentic V3-glycan bNAbs UCAs available to screen candidate priming immunogens. Priming such a diverse array of bNAb precursors can be viewed as another metric of success for future V3-glycan-targeted vaccines.

Cryo-EM analysis revealed that the SHIV.5MUT-induced bNAbs bound to Env with a wide range of approach angles, recapitulating the structural diversity seen in V3-glycan bNAbs isolated from humans (*13*). Indeed, several important structural parallels emerged between rhesus and human V3-glycan bNAbs. For example, while AM12-352 (rhesus) and PGT128 (human) use distinct binding poses, their CDRH3 loops adopted nearly identical architectures to interact with the ^324^GDIR_327_ motif **(Fig. 2F)** and both lineages acquired insertions in CDRH2 that enable glycan accommodation **(Fig. 2I)** (*56*). Similarly, rhesus bNAbs AJ09-83 and AJ09-110 use negatively charged residues in their CDRH3 loops to contact R327 within the ^324^GDIR_327_ motif **(Fig. 2B),** mimicking an interaction observed in structures of human V3-glycan bNAbs including 10-1074 **(Fig. 2C)** (*54*). Additionally, our data demonstrate that antibodies can converge on similar binding modes despite using distinct germline building blocks, as exemplified by rhesus bNAbs AJ09-83 and V634-136, which bind Env with similar poses but are derived from different V_H_ gene segments **(Fig. 2E)**. Overall, the comprehensive structural analyses of ten mature SHIV.5MUT-elicited V3-glycan bNAbs and one UCA in complex with Env trimers support the relevance of the rhesus model for immunogen design.

One of the most interesting findings of the current study is the mechanism by which SHIV.5MUT elicited V3-glycan bNAbs with such consistency, despite differing from wildtype SHIV.BG505.N332 by only four residues in V1 **(Fig. 1B)**. While none of these mutations alter PNGS sequons, we found that the asparagine-to-proline substitution at position 136 in 5MUT abrogated glycosylation at the adjacent N133 site **(Fig. 4E-F)**, consistent with previous reports that proline residues adversely affect N-glycan occupancy at neighboring PNGS (*31, 71, 72*). This hypoglycosylation rendered the V1 region of the 5MUT Env highly immunogenic, inducing an early wave of neutralizing antibodies that in turn selected for viral escape variants with shortened V1 loops **(Fig. 4A-B)**. These variants selectively exposed the underlying ^324^GDIR_327_ epitope **(Fig. 5E),** allowing them to engage V3-glycan bNAb precursors **(Fig. 5F).** Subsequent diversification in the V1 and V3 regions of Env then guided bNAb maturation to breadth and potency by recapitulating the natural sequence diversity found in circulating HIV-1 strains **(Fig. 7E-F)**. Altogether, these findings coalesce into a model of V3-glycan bNAb induction in SHIV.5MUT-infected macaques that is illustrated in **Fig. 5A**.

A similar mechanism of V1-directed antibodies selecting for viral variants with shortened V1 loops was previously proposed to be responsible for the delayed induction of V3-glycan bNAbs observed in human subject CH848 (*19*) and in two macaques infected with the Env-matched SHIV.CH848 (*87*). Until now, however, direct evidence was lacking. Nonetheless, these findings spurred the development of a germline-targeting immunogen with a short, glycan-deficient V1 loop designed to elicit DH270-like antibodies that has recently entered human clinical trials (HVTN307 / NCT05903339) (*19, 28*). Other immunogens designed to elicit BG18-like antibodies also have heavily engineered V1 loops that lack glycans, and these too have entered human clinical trials (HVTN144 / NCT06033209) (*30, 31*). Altogether, the data indicate that V1 modification is a promising strategy for enhancing Env binding to V3-glycan bNAb precursors. A key aspect of the current study is that SHIV.5MUT infection has revealed a set of novel priming immunogens that can stimulate an even larger and more diverse group of V3-glycan bNAb precursors, which can serve as a standard for future vaccine trials. More broadly, sterically increasing epitope exposure by glycan elimination or shortening of hypervariable loops may serve as a generalizable strategy for inducing bNAbs against HIV-1 and other difficult-to-neutralize viruses. Indeed, we recently showed that structure-guided deletion of glycans adjacent to the CD4 binding site markedly enhanced elicitation of bNAbs to this epitope in a SHIV model (*91*). This approach could potentially be extended to other viral systems such as the hepatitis C virus E1E2 glycoprotein, which contains a flexible hypervariable region (HVR1) that partially occludes bNAb epitopes adjacent to the host receptor binding site (*92–94*). E1E2 variants with a shortened HVR1 may be analogous to our del4 and del8 Envs, increasing epitope accessibility and potentially enhancing bNAb precursor binding (*95, 96*).

SHIV.5MUT infection not only primed V3-glycan bNAb lineages by *in vivo* generation of V1-deleted Env variants but also guided their maturation to breadth and potency via Env-bNAb coevolution. To understand which specific SHIV.5MUT Env variants promoted bNAb maturation, we compared the kinetics of Env sequence diversification and acquisition of heterologous plasma neutralization breadth in each animal. This analysis identified shared routes of Env-bNAb coevolution across multiple SHIV.5MUT-infected macaques, suggesting a general framework for vaccine design. Following bNAb precursor priming by early V1-deleted Env variants including del4 and del8, the viral quasispecies continued to diversify in response to antibody pressure. Therefore, bNAb lineage affinity maturation occurred in the presence of Env escape variants that positively selected antibody lineage members capable of tolerating these very resistance mutations. If these Env resistance mutations are well-represented in global M-group viruses, they can drive breadth acquisition by the evolving bNAb lineage. Here, we identified ten selected sites and two V1 hypervariable region modifications (increased length and glycosylation) that were significantly enriched in SHIV.5MUT-infected macaques that developed bNAbs **(Fig. 6B)**. Emergence of mutations at these sites *in vivo* preceded or coincided with the ability of the plasma to neutralize heterologous viruses that carried the mutant residues, suggesting that exposure to these Env variants promoted bNAb maturation toward breadth. Overall, the shared Env-bNAb coevolutionary pathways described here can serve as a roadmap for a sequential vaccination regimen. Specifically, we propose using del8 and del4 immunogens to prime *bona fide* V3-glycan bNAb precursors and using immunogens bearing longer and more glycosylated V1 loops and common resistance mutations at the ten key sites identified here to boost these responses toward breadth and potency. Moreover, initial boost immunogens should contain variation at the subset of sites that are in close physical proximity to the V3-glycan epitope and that exhibit mutation kinetics coinciding with breadth acquisition **(Fig. 6C)**. We have recently initiated a preclinical trial in macaques testing these concepts.

The Envs identified here are uniquely positioned to be rapidly translated into immunogens for the next generation of human vaccine trials. Our data demonstrate that specific Env variants sampled during SHIV.5MUT infection successfully elicit diverse V3-glycan bNAbs with multiple, distinct immunogenetic and structural solutions to epitope engagement. Given that the human and macaque antibody repertoires are comparable in size and diversity (*97*), several such solutions for analogous bNAbs should be available in humans. The ability to induce such diverse responses would be a desirable property in a vaccine, as it would maximize effectiveness in subjects with disparate immunoglobulin repertoires. Conversely, if SHIV.5MUT infection had led to a single, stereotypical bNAb solution, then elicitation of similar antibodies in humans would require certain critical immunogenetic features to be widespread in the human repertoire. For example, the CD4bs immunogen eOD-GT8 requires permissive VH1-2*02 or *04 alleles, and clinical trial participants lacking these were non-responders (*98*).

A limitation of this study is that not all SHIV.5MUT-infected animals developed V3-glycan bNAbs even though shortened-V1 Env variants with priming potential were present in the early viral quasispecies of most animals. We suspect this is due in large part to insufficient antigen drive for bNAb priming and maturation, as macaques that did not develop bNAbs had significantly lower plasma viral loads **(fig. S17A)** and autologous neutralization titers **(fig. S17B)** than those that did develop bNAbs, consistent with previous studies (*86–88*). If sufficiently high levels of antigen can be delivered by vaccination or if avidity can be enhanced via multimerization on nanoparticles or virus-like particles (*99, 100*), it may be possible that a larger proportion of animals would be expected to develop V3-glycan bNAbs that have even greater breadth and potency. Regardless, the outbred animal model of V3-glycan bNAb elicitation described here can be leveraged to rigorously and iteratively dissect the challenges of inducing clinically protective bNAb responses and to inform future vaccine trials.

## Acknowledgments

We are grateful to the staff at Bioqual, Inc. for the diligent care of nonhuman primates and technical support, to the staff of the Nonhuman Primate Core Virological Laboratory for AIDS Vaccine Research and Development at Duke University for determination of plasma viral RNA levels, to Casey Lee for technical support, to the NIAID/DAIDS Simian Vaccine Evaluation Unit, and to Nancy Miller for advice and support in the development of this project.

## Author contributions

Conceptualization: ANS, HBG, HL, EG, PJB, BHH, GMS.

Investigation: ANS, HBG, HL, EG, LM, MLN, WL, MSC, KW (Winters), CGG, RAO, MJA, KA, YL, AS, KC, YP, CZ, XL, JWC, AA, JRK, RH, DJM, FBR, NSK, SJP, CLM, JL, EJDW, MGL, AAW, GA, NM, KA.

Formal analysis: ANS, HBG, HL, AJC, MPH, MLN, APW, EVI, ATD, KW (Wagh), BK. Supervision: ANS, EFK, JDA, FBR, KJW, BTK, BFH, MAM, MCC, MSS, DJI, RA, MC, DW, PJB, BHH, GMS.

Funding acquisition: ANS, BFH, PJB, BHH, GMS. Writing – original draft: ANS, HBG, PJB, BHH, GMS. Writing – review & editing: All authors.

## Data and materials availability

All monoclonal antibody isolate sequences are deposited at GenBank under accession numbers PX059660-PX060135 and PX390224-PX391126. BCR repertoire next-generation sequencing data are available under the NCBI BioProject PRJNA1305399. Longitudinal Env SGS gp140 sequences are deposited at GenBank under accession numbers PX297533-PX306283 and MN467402-MN472740. Models and cryo-EM density maps were deposited in the PDB (9YHO, 9YHQ, 9YHR, 9YHS, 9YHT, 9YIB, 9YID, 9YIE, 9YIF, 9YIG, 9YIE, 9YII, 9YIJ, 9YIK, 9YIL) and EMDB (EMD-72969 through EMD-72973 and EMD-72985 through EMD-72994). All other data are available in the main text or supplementary materials.

## Supplementary Materials

Materials and Methods

Supplementary Text

Figs. S1 to S31

Tables S1 to S3

## References

1. A. Pegu et al., A meta-analysis of passive immunization studies shows that serum-neutralizing antibody titer associates with protection against SHIV challenge. Cell Host Microbe 26, 336–346.e333 (2019). <10.1016/j.chom.2019.08.014>

2. M. Shingai et al., Passive transfer of modest titers of potent and broadly neutralizing anti-HIV monoclonal antibodies block SHIV infection in macaques. J Exp Med 211, 2061– 2074 (2014). <10.1084/jem.20132494>

3. A. Pegu et al., Neutralizing antibodies to HIV-1 envelope protect more effectively in vivo than those to the CD4 receptor. Sci Transl Med 6, 243ra288 (2014). <10.1126/scitranslmed.3008992>

4. B. Moldt et al., Highly potent HIV-specific antibody neutralization in vitro translates into effective protection against mucosal SHIV challenge in vivo. Proc Natl Acad Sci USA 109, 18921–18925 (2012). <10.1073/pnas.1214785109>

5. L. Corey et al., Two randomized trials of neutralizing antibodies to prevent HIV-1 acquisition. N Engl J Med 384, 1003–1014 (2021). <10.1056/NEJMoa2031738>

6. P. B. Gilbert et al., Neutralization titer biomarker for antibody-mediated prevention of HIV-1 acquisition. Nat Med 28, 1924–1932 (2022). <10.1038/s41591-022-01953-6>

7. Y. Nishimura et al., Early antibody therapy can induce long-lasting immunity to SHIV. Nature 543, 559–563 (2017). <10.1038/nature21435>

8. E. N. Borducchi et al., Antibody and TLR7 agonist delay viral rebound in SHIV-infected monkeys. Nature 563, 360–364 (2018). <10.1038/s41586-018-0600-6>

9. C. Gaebler et al., Prolonged viral suppression with anti-HIV-1 antibody therapy. Nature 606, 368–374 (2022). <10.1038/s41586-022-04597-1>

10. M. C. Sneller et al., Combination anti-HIV antibodies provide sustained virological suppression. Nature 606, 375–381 (2022). <10.1038/s41586-022-04797-9>

11. J. F. Scheid et al., HIV-1 antibody 3BNC117 suppresses viral rebound in humans during treatment interruption. Nature 535, 556–560 (2016). <10.1038/nature18929>

12. E. Landais et al., Broadly neutralizing antibody responses in a large longitudinal sub-saharan HIV primary infection cohort. PLoS Pathog 12, e1005369 (2016). <10.1371/journal.ppat.1005369>

13. L. Kong et al., Supersite of immune vulnerability on the glycosylated face of HIV-1 envelope glycoprotein gp120. Nat Struct Mol Biol 20, 796–803 (2013). <10.1038/nsmb.2594>

14. R. Pejchal et al., A potent and broad neutralizing antibody recognizes and penetrates the HIV glycan shield. Science 334, 1097–1103 (2011). <10.1126/science.1213256>

15. D. T. MacLeod et al., Early antibody lineage diversification and independent limb maturation lead to broad HIV-1 neutralization targeting the Env high-mannose patch. Immunity 44, 1215–1226 (2016). <10.1016/j.immuni.2016.04.016>

16. D. Sok et al., A prominent site of antibody vulnerability on HIV envelope incorporates a motif associated with CCR5 binding and its camouflaging glycans. Immunity 45, 31–45 (2016). <10.1016/j.immuni.2016.06.026>

17. D. R. Burton, L. Hangartner, Broadly neutralizing antibodies to HIV and their role in vaccine design. Annu Rev Immunol 34, 635–659 (2016). <10.1146/annurev-immunol-041015-055515>

18. B. F. Haynes et al., Strategies for HIV-1 vaccines that induce broadly neutralizing antibodies. Nat Rev Immunol 23, 142–158 (2023). <10.1038/s41577-022-00753-w>

19. M. Bonsignori et al., Staged induction of HIV-1 glycan-dependent broadly neutralizing antibodies. Sci Transl Med 9, (2017). <10.1126/scitranslmed.aai7514>

20. N. T. Freund et al., Coexistence of potent HIV-1 broadly neutralizing antibodies and antibody-sensitive viruses in a viremic controller. Sci Transl Med 9, (2017). <10.1126/scitranslmed.aal2144>

21. C. A. Bricault et al., HIV-1 neutralizing antibody signatures and application to epitope-targeted vaccine design. Cell Host Microbe 25, 59–72.e58 (2019). <10.1016/j.chom.2018.12.001>

22. L. M. Walker et al., Broad neutralization coverage of HIV by multiple highly potent antibodies. Nature 477, 466–470 (2011). <10.1038/nature10373>

23. D. R. Burton, What are the most powerful immunogen design vaccine strategies? Reverse vaccinology 2.0 shows great promise. Cold Spring Harb Perspect Biol 9, (2017). <10.1101/cshperspect.a030262>

24. J. M. Steichen et al., HIV vaccine design to target germline precursors of glycan-dependent broadly neutralizing antibodies. Immunity 45, 483–496 (2016). <10.1016/j.immuni.2016.08.016>

25. A. Escolano et al., Sequential immunization elicits broadly neutralizing anti-HIV-1 antibodies in Ig knockin mice. Cell 166, 1445–1458.e1412 (2016). <10.1016/j.cell.2016.07.030>

26. A. Escolano et al., Immunization expands B cells specific to HIV-1 V3 glycan in mice and macaques. Nature 570, 468–473 (2019). <10.1038/s41586-019-1250-z>

27. A. Escolano et al., Sequential immunization of macaques elicits heterologous neutralizing antibodies targeting the V3-glycan patch of HIV-1 Env. Sci Transl Med 13, eabk1533 (2021). <10.1126/scitranslmed.abk1533>

28. K. O. Saunders et al., Targeted selection of HIV-specific antibody mutations by engineering B cell maturation. Science 366, (2019). <10.1126/science.aay7199>

29. O. M. Swanson et al., An engineered immunogen activates diverse HIV broadly neutralizing antibody precursors and promotes acquisition of improbable mutations. Sci Transl Med 17, eadr2218 (2025). <10.1126/scitranslmed.adr2218>

30. J. M. Steichen et al., A generalized HIV vaccine design strategy for priming of broadly neutralizing antibody responses. Science 366, (2019). <10.1126/science.aax4380>

31. J. M. Steichen et al., Vaccine priming of rare HIV broadly neutralizing antibody precursors in nonhuman primates. Science 384, eadj8321 (2024). <10.1126/science.adj8321>

32. Z. Xie et al., mRNA-LNP HIV-1 trimer boosters elicit precursors to broad neutralizing antibodies. Science 384, eadk0582 (2024). <10.1126/science.adk0582>

33. A. T. DeLaitsch et al., Neutralizing antibodies elicited in macaques recognize V3 residues on altered conformations of HIV-1 Env trimer. NPJ Vaccines 9, 240 (2024). <10.1038/s41541-024-01038-0>

34. Z. Yang et al., Neutralizing antibodies induced in immunized macaques recognize the CD4-binding site on an occluded-open HIV-1 envelope trimer. Nat Commun 13, 732 (2022). <10.1038/s41467-022-28424-3>

35. Z. Yang, K. A. Dam, J. M. Gershoni, S. Zolla-Pazner, P. J. Bjorkman, Antibody recognition of CD4-induced open HIV-1 Env trimers. J Virol 96, e0108222 (2022). <10.1128/jvi.01082-22>

36. M. K. Gorny et al., Cross-clade neutralizing activity of human anti-V3 monoclonal antibodies derived from the cells of individuals infected with non-B clades of human immunodeficiency virus type 1. J Virol 80, 6865–6872 (2006). <10.1128/jvi.02202-05>

37. X. Jiang et al., Conserved structural elements in the V3 crown of HIV-1 gp120. Nat Struct Mol Biol 17, 955–961 (2010). <10.1038/nsmb.1861>

38. T. Tiller et al., Efficient generation of monoclonal antibodies from single human B cells by single cell RT-PCR and expression vector cloning. J Immunol Methods 329, 112–124 (2008). <10.1016/j.jim.2007.09.017>

39. R. D. Mason et al., Targeted isolation of antibodies directed against major sites of SIV Env vulnerability. PLoS Pathog 12, e1005537 (2016). <10.1371/journal.ppat.1005537>

40. R. P. Staupe et al., Single cell multi-omic reference atlases of non-human primate immune tissues reveals CD102 as a biomarker for long-lived plasma cells. Commun Biol 5, 1399 (2022). <10.1038/s42003-022-04216-9>

41. H. Mouquet et al., Complex-type N-glycan recognition by potent broadly neutralizing HIV antibodies. Proc Natl Acad Sci USA 109, E3268–3277 (2012). <10.1073/pnas.1217207109>

42. K. E. Stephenson, K. Wagh, B. Korber, D. H. Barouch, Vaccines and broadly neutralizing antibodies for HIV-1 prevention. Annu Rev Immunol 38, 673–703 (2020). <10.1146/annurev-immunol-080219-023629>

43. N. Vázquez Bernat et al., Rhesus and cynomolgus macaque immunoglobulin heavy-chain genotyping yields comprehensive databases of germline vdj alleles. Immunity 54, 355–366.e354 (2021). <10.1016/j.immuni.2020.12.018>

44. A. Ramesh et al., Structure and diversity of the rhesus macaque immunoglobulin loci through multiple de novo genome assemblies. Front Immunol 8, 1407 (2017). <10.3389/fimmu.2017.01407>

45. F. Matsuda et al., The complete nucleotide sequence of the human immunoglobulin heavy chain variable region locus. J Exp Med 188, 2151–2162 (1998). <10.1084/jem.188.11.2151>

46. J. Jardine et al., Rational HIV immunogen design to target specific germline B cell receptors. Science 340, 711–716 (2013). <10.1126/science.1234150>

47. J. G. Jardine et al., HIV-1 broadly neutralizing antibody precursor B cells revealed by germline-targeting immunogen. Science 351, 1458–1463 (2016). <10.1126/science.aad9195>

48. R. Habib et al., Env-antibody coevolution identifies B cell priming as the principal bottleneck to HIV-1 V2 apex broadly neutralizing antibody development. bioRxiv, (2025). <10.1101/2025.05.03.652068>

49. R. S. Roark et al., Structural and genetic basis of HIV-1 envelope V2 apex recognition by rhesus broadly neutralizing antibodies. J Exp Med 222, (2025). <10.1084/jem.20250638>

50. B. Briney, A. Inderbitzin, C. Joyce, D. R. Burton, Commonality despite exceptional diversity in the baseline human antibody repertoire. Nature 566, 393–397 (2019). <10.1038/s41586-019-0879-y>

51. C. Joyce, D. R. Burton, B. Briney, Comparisons of the antibody repertoires of a humanized rodent and humans by high throughput sequencing. Sci Rep 10, 1120 (2020). <10.1038/s41598-020-57764-7>

52. J. D. Galson et al., Analysis of B cell repertoire dynamics following hepatitis B vaccination in humans, and enrichment of vaccine-specific antibody sequences. EBioMedicine 2, 2070–2079 (2015). <10.1016/j.ebiom.2015.11.034>

53. R. W. Sanders et al., A next-generation cleaved, soluble HIV-1 Env trimer, BG505 SOSIP.664 gp140, expresses multiple epitopes for broadly neutralizing but not non-neutralizing antibodies. PLoS Pathog 9, e1003618 (2013). <10.1371/journal.ppat.1003618>

54. H. B. Gristick et al., Natively glycosylated HIV-1 Env structure reveals new mode for antibody recognition of the CD4-binding site. Nat Struct Mol Biol 23, 906–915 (2016). <10.1038/nsmb.3291>

55. A. P. West, Jr., R. Diskin, M. C. Nussenzweig, P. J. Bjorkman, Structural basis for germ-line gene usage of a potent class of antibodies targeting the CD4-binding site of HIV-1 gp120. Proc Natl Acad Sci USA 109, E2083–2090 (2012). <10.1073/pnas.1208984109>

56. K. J. Doores et al., Two classes of broadly neutralizing antibodies within a single lineage directed to the high-mannose patch of HIV envelope. J Virol 89, 1105–1118 (2015). <10.1128/jvi.02905-14>

57. M. M. Corcoran et al., Production of individualized V gene databases reveals high levels of immunoglobulin genetic diversity. Nat Commun 7, 13642 (2016). <10.1038/ncomms13642>

58. C. A. Schramm et al., Sonar: A high-throughput pipeline for inferring antibody ontogenies from longitudinal sequencing of B cell transcripts. Front Immunol 7, 372 (2016). <10.3389/fimmu.2016.00372>

59. J. S. Martin Beem, S. Venkatayogi, B. F. Haynes, K. Wiehe, Armadillo: A web server for analyzing antibody mutation probabilities. Nucleic Acids Res 51, W51–w56 (2023). <10.1093/nar/gkad398>

60. A. M. Collins et al., AIRR-C ig reference sets: Curated sets of immunoglobulin heavy and light chain germline genes. Front Immunol 14, 1330153 (2023). <10.3389/fimmu.2023.1330153>

61. J. F. Salazar-Gonzalez et al., Deciphering human immunodeficiency virus type 1 transmission and early envelope diversification by single-genome amplification and sequencing. J Virol 82, 3952–3970 (2008). <10.1128/jvi.02660-07>

62. B. F. Keele et al., Identification and characterization of transmitted and early founder virus envelopes in primary HIV-1 infection. Proc Natl Acad Sci USA 105, 7552–7557 (2008). <10.1073/pnas.0802203105>

63. X. Wei et al., Viral dynamics in human immunodeficiency virus type 1 infection. Nature 373, 117–122 (1995). <10.1038/373117a0>

64. A. S. Perelson, A. U. Neumann, M. Markowitz, J. M. Leonard, D. D. Ho, HIV-1 dynamics in vivo: Virion clearance rate, infected cell life-span, and viral generation time. Science 271, 1582–1586 (1996). <10.1126/science.271.5255.1582>

65. H. Wang et al., Potent and broad HIV-1 neutralization in fusion peptide-primed SHIV-infected macaques. Cell 187, 7214–7231.e7223 (2024). <10.1016/j.cell.2024.10.003>

66. K. J. Bar et al., Early low-titer neutralizing antibodies impede HIV-1 replication and select for virus escape. PLoS Pathog 8, e1002721 (2012). <10.1371/journal.ppat.1002721>

67. W. B. Struwe et al., Site-specific glycosylation of virion-derived HIV-1 Env is mimicked by a soluble trimeric immunogen. Cell Rep 24, 1958–1966.e1955 (2018). <10.1016/j.celrep.2018.07.080>

68. R. Derking et al., Enhancing glycan occupancy of soluble HIV-1 envelope trimers to mimic the native viral spike. Cell Rep 35, 108933 (2021). <10.1016/j.celrep.2021.108933>

69. L. E. McCoy et al., Holes in the glycan shield of the native HIV envelope are a target of trimer-elicited neutralizing antibodies. Cell Rep 16, 2327–2338 (2016). <10.1016/j.celrep.2016.07.074>

70. P. J. Klasse et al., Epitopes for neutralizing antibodies induced by HIV-1 envelope glycoprotein BG505 SOSIP trimers in rabbits and macaques. PLoS Pathog 14, e1006913 (2018). <10.1371/journal.ppat.1006913>

71. J. L. Mellquist, L. Kasturi, S. L. Spitalnik, S. H. Shakin-Eshleman, The amino acid following an asn-X-Ser/Thr sequon is an important determinant of N-linked core glycosylation efficiency. Biochemistry 37, 6833–6837 (1998). <10.1021/bi972217k>

72. M. Bañó-Polo, F. Baldin, S. Tamborero, M. A. Marti-Renom, I. Mingarro, N-glycosylation efficiency is determined by the distance to the C-terminus and the amino acid preceding an Asn-Ser-Thr sequon. Protein Sci 20, 179–186 (2011). <10.1002/pro.551>

73. J. P. Julien et al., Crystal structure of a soluble cleaved HIV-1 envelope trimer. Science 342, 1477–1483 (2013). <10.1126/science.1245625>

74. R. Henderson et al., Structural basis for breadth development in the HIV-1 V3-glycan targeting dh270 antibody clonal lineage. Nat Commun 14, 2782 (2023). <10.1038/s41467-023-38108-1>

75. J. H. Lee et al., A broadly neutralizing antibody targets the dynamic HIV envelope trimer apex via a long, rigidified, and anionic β-hairpin structure. Immunity 46, 690–702 (2017). <10.1016/j.immuni.2017.03.017>

76. J. Gorman et al., Structures of HIV-1 Env V1V2 with broadly neutralizing antibodies reveal commonalities that enable vaccine design. Nat Struct Mol Biol 23, 81–90 (2016). <10.1038/nsmb.3144>

77. G. D. Victora, M. C. Nussenzweig, Germinal centers. Annu Rev Immunol 40, 413–442 (2022). <10.1146/annurev-immunol-120419-022408>

78. M. Bonsignori et al., Maturation pathway from germline to broad HIV-1 neutralizer of a CD4-mimic antibody. Cell 165, 449–463 (2016). <10.1016/j.cell.2016.02.022>

79. B. Korber, P. Hraber, K. Wagh, B. H. Hahn, Polyvalent vaccine approaches to combat HIV-1 diversity. Immunol Rev 275, 230–244 (2017). <10.1111/imr.12516>

80. P. Hraber et al., Longitudinal antigenic sequences and sites from intra-host evolution (LASSIE) identifies immune-selected HIV variants. Viruses 7, 5443–5475 (2015). <10.3390/v7102881>

81. G. S. Lambert, C. Upadhyay, HIV-1 envelope glycosylation and the signal peptide. Vaccines (Basel) 9, (2021). <10.3390/vaccines9020176>

82. C. Upadhyay et al., Signal peptide of HIV-1 envelope modulates glycosylation impacting exposure of V1V2 and other epitopes. PLoS Pathog 16, e1009185 (2020). <10.1371/journal.ppat.1009185>

83. C. Upadhyay et al., Signal peptide exchange alters HIV-1 envelope antigenicity and immunogenicity. Front Immunol 15, 1476924 (2024). <10.3389/fimmu.2024.1476924>

84. K. Xu et al., Epitope-based vaccine design yields fusion peptide-directed antibodies that neutralize diverse strains of HIV-1. Nat Med 24, 857–867 (2018). <10.1038/s41591-018-0042-6>

85. I. Mikell et al., Characteristics of the earliest cross-neutralizing antibody response to HIV-1. PLoS Pathog 7, e1001251 (2011). <10.1371/journal.ppat.1001251>

86. E. S. Gray et al., The neutralization breadth of HIV-1 develops incrementally over four years and is associated with CD4+ T cell decline and high viral load during acute infection. J Virol 85, 4828–4840 (2011). <10.1128/jvi.00198-11>

87. R. S. Roark et al., Recapitulation of HIV-1 Env-antibody coevolution in macaques leading to neutralization breadth. Science 371, (2021). <10.1126/science.abd2638>

88. P. Rusert et al., Determinants of HIV-1 broadly neutralizing antibody induction. Nat Med 22, 1260–1267 (2016). <10.1038/nm.4187>

89. L. Gieselmann et al., Identification of a broad and potent V3 glycan site bNAb targeting an n332gp120 glycan-independent epitope. bioRxiv, (2025). <10.1101/2025.09.05.674437>

90. O. Swanson et al., Identification of CDRH3 loops in the B cell receptor repertoire that can be engaged by candidate immunogens. PLoS Pathog 19, e1011401 (2023). <10.1371/journal.ppat.1011401>

91. D. J. Morris et al., Transient glycan shield reduction induces CD4-binding site broadly neutralizing antibodies in SHIV-infected macaques. Cell Rep 44, 115848 (2025). <10.1016/j.celrep.2025.115848>

92. L. Kong et al., Hepatitis C virus E2 envelope glycoprotein core structure. Science 342, 1090–1094 (2013). <10.1126/science.1243876>

93. A. Torrents de la Peña et al., Structure of the hepatitis C virus E1E2 glycoprotein complex. Science 378, 263–269 (2022). <10.1126/science.abn9884>

94. N. Frumento, A. I. Flyak, J. R. Bailey, Mechanisms of HCV resistance to broadly neutralizing antibodies. Curr Opin Virol 50, 23–29 (2021). <10.1016/j.coviro.2021.07.003>

95. D. Bankwitz et al., Hepatitis C virus hypervariable region 1 modulates receptor interactions, conceals the CD81 binding site, and protects conserved neutralizing epitopes. J Virol 84, 5751–5763 (2010). <10.1128/jvi.02200-09>

96. J. Prentoe et al., Hypervariable region 1 and N-linked glycans of hepatitis C regulate virion neutralization by modulating envelope conformations. Proc Natl Acad Sci USA 116, 10039–10047 (2019). <10.1073/pnas.1822002116>

97. V. Vigdorovich et al., Repertoire comparison of the B-cell receptor-encoding loci in humans and rhesus macaques by next-generation sequencing. Clin Transl Immunology 5, e93 (2016). <10.1038/cti.2016.42>

98. J. R. Willis et al., Vaccination with mRNA-encoded nanoparticles drives early maturation of HIV bnab precursors in humans. Science, eadr8382 (2025). <10.1126/science.adr8382>

99. T. U. J. Bruun, A. C. Andersson, S. J. Draper, M. Howarth, Engineering a rugged nanoscaffold to enhance plug-and-display vaccination. ACS Nano 12, 8855–8866 (2018). <10.1021/acsnano.8b02805>

100. J. Guenaga et al., HIV Env trimers elicit NHP apex cross-neutralizing antibodies mimicking human bNAbs. bioRxiv, (2025). <10.1101/2025.09.19.677470>

101. C. M. Ives et al., Restoring protein glycosylation with glycoshape. Nat Methods 21, 2117–2127 (2024). <10.1038/s41592-024-02464-7>

102. E. C. Meng et al., UCSF ChimeraX: Tools for structure building and analysis. Protein Sci 32, e4792 (2023). <10.1002/pro.4792>

103. J. H. Lee et al., Long-primed germinal centres with enduring affinity maturation and clonal migration. Nature 609, 998–1004 (2022). <10.1038/s41586-022-05216-9>

104. H. B. Gristick, et al., CD4 binding site immunogens elicit heterologous anti-HIV-1 neutralizing antibodies in transgenic and wild-type animals. Sci Immunol 8, eade6364 (2023). <10.1126/sciimmunol.ade6364>

105. A. H. Keeble et al., Approaching infinite affinity through engineering of peptide-protein interaction. Proc Natl Acad Sci USA 116, 26523–26533 (2019). <10.1073/pnas.1909653116>

106. L. Scharf et al., Broadly neutralizing antibody 8ANC195 recognizes closed and open states of HIV-1 Env. Cell 162, 1379–1390 (2015). <10.1016/j.cell.2015.08.035>

107. R. Rahikainen et al., Overcoming symmetry mismatch in vaccine nanoassembly through spontaneous amidation. Angew Chem Int Ed Engl 60, 321–330 (2021). <10.1002/anie.202009663>

108. A. A. Cohen et al., Mosaic nanoparticles elicit cross-reactive immune responses to zoonotic coronaviruses in mice. Science 371, 735–741 (2021). <10.1126/science.abf6840>

109. M. G. Alameh et al., Lipid nanoparticles enhance the efficacy of mRNA and protein subunit vaccines by inducing robust T follicular helper cell and humoral responses. Immunity 54, 2877–2892.e2877 (2021). <10.1016/j.immuni.2021.11.001>

110. Z. Mu et al., mRNA-encoded HIV-1 Env trimer ferritin nanoparticles induce monoclonal antibodies that neutralize heterologous HIV-1 isolates in mice. Cell Rep 38, 110514 (2022). <10.1016/j.celrep.2022.110514>

111. M. Baiersdörfer et al., A facile method for the removal of dsRNA contaminant from in vitro-transcribed mRNA. Mol Ther Nucleic Acids 15, 26–35 (2019). <10.1016/j.omtn.2019.02.018>

112. K. Karikó, M. Buckstein, H. Ni, D. Weissman, Suppression of RNA recognition by toll-like receptors: The impact of nucleoside modification and the evolutionary origin of RNA. Immunity 23, 165–175 (2005). <10.1016/j.immuni.2005.06.008>

113. M. Jayaraman et al., Maximizing the potency of siRNA lipid nanoparticles for hepatic gene silencing in vivo. Angew Chem Int Ed Engl 51, 8529–8533 (2012). <10.1002/anie.201203263>

114. M. A. Maier et al., Biodegradable lipids enabling rapidly eliminated lipid nanoparticles for systemic delivery of RNAi therapeutics. Mol Ther 21, 1570–1578 (2013). <10.1038/mt.2013.124>

115. H. Li et al., Envelope residue 375 substitutions in simian-human immunodeficiency viruses enhance CD4 binding and replication in rhesus macaques. Proc Natl Acad Sci USA 113, E3413–3422 (2016). <10.1073/pnas.1606636113>

116. X. Wei et al., Antibody neutralization and escape by HIV-1. Nature 422, 307–312 (2003). <10.1038/nature01470>

117. M. Sarzotti-Kelsoe et al., Optimization and validation of the TZM-bl assay for standardized assessments of neutralizing antibodies against HIV-1. J Immunol Methods 409, 131–146 (2014). <10.1016/j.jim.2013.11.022>

118. X. Brochet, M. P. Lefranc, V. Giudicelli, IMGT/V-quest: The highly customized and integrated system for ig and tr standardized V-J and V-D-J sequence analysis. Nucleic Acids Res 36, W503–508 (2008). <10.1093/nar/gkn316>

119. Q. Han et al., Neonatal rhesus macaques have distinct immune cell transcriptional profiles following HIV envelope immunization. Cell Rep 30, 1553–1569.e1556 (2020). <10.1016/j.celrep.2019.12.091>

120. S. J. Krebs et al., Longitudinal analysis reveals early development of three MPER-directed neutralizing antibody lineages from an HIV-1-infected individual. Immunity 50, 677–691.e613 (2019). <10.1016/j.immuni.2019.02.008>

121. N. Vázquez Bernat et al., High-quality library preparation for NGS-based immunoglobulin germline gene inference and repertoire expression analysis. Front Immunol 10, 660 (2019). <10.3389/fimmu.2019.00660>

122. V. Bhardwaj, M. Franceschetti, R. Rao, P. A. Pevzner, Y. Safonova, Automated analysis of immunosequencing datasets reveals novel immunoglobulin D genes across diverse species. PLoS Comput Biol 16, e1007837 (2020). <10.1371/journal.pcbi.1007837>

123. C. O. Barnes et al., SARS-CoV-2 neutralizing antibody structures inform therapeutic strategies. Nature 588, 682–687 (2020). <10.1038/s41586-020-2852-1>

124. D. N. Mastronarde, Automated electron microscope tomography using robust prediction of specimen movements. J Struct Biol 152, 36–51 (2005). <10.1016/j.jsb.2005.07.007>

125. A. Punjani, J. L. Rubinstein, D. J. Fleet, M. A. Brubaker, Cryosparc: Algorithms for rapid unsupervised cryo-EM structure determination. Nat Methods 14, 290–296 (2017). <10.1038/nmeth.4169>

126. A. Casañal, B. Lohkamp, P. Emsley, Current developments in coot for macromolecular model building of electron cryo-microscopy and crystallographic data. Protein Sci 29, 1069–1078 (2020). <10.1002/pro.3791>

127. P. V. Afonine et al., Real-space refinement in phenix for cryo-EM and crystallography. Acta Crystallogr D Struct Biol 74, 531–544 (2018). <10.1107/s2059798318006551>

128. V. B. Chen et al., Molprobity: All-atom structure validation for macromolecular crystallography. Acta Crystallogr D Biol Crystallogr 66, 12–21 (2010). <10.1107/s0907444909042073>

129. W. L. Delano, in *The PyMOL Molecular Graphics System*. (Schrödinger, LLC, New York, NY, USA, 2002).

130. E. Krissinel, K. Henrick, Inference of macromolecular assemblies from crystalline state. J Mol Biol 372, 774–797 (2007). <10.1016/j.jmb.2007.05.022>

131. A. Tareen, J. B. Kinney, Logomaker: Beautiful sequence logos in python. Bioinformatics 36, 2272–2274 (2020). <10.1093/bioinformatics/btz921>

132. P. Virtanen et al., Scipy 1.0: Fundamental algorithms for scientific computing in python. Nat Methods 17, 261–272 (2020). <10.1038/s41592-019-0686-2>

133. N. Pardi et al., Zika virus protection by a single low-dose nucleoside-modified mRNA vaccination. Nature 543, 248–251 (2017). <10.1038/nature21428>

134. S. K. Sharma et al., Cleavage-independent HIV-1 Env trimers engineered as soluble native spike mimetics for vaccine design. Cell Rep 11, 539–550 (2015). <10.1016/j.celrep.2015.03.047>

135. D. Wrapp et al., Structure-based stabilization of SOSIP Env enhances recombinant ectodomain durability and yield. J Virol 97, e0167322 (2023). <10.1128/jvi.01673-22>

136. M. J. Hogan et al., Increased surface expression of HIV-1 envelope is associated with improved antibody response in vaccinia prime/protein boost immunization. Virology 514, 106–117 (2018). <10.1016/j.virol.2017.10.013>

137. M. Silva, et al., A particulate saponin/TLR agonist vaccine adjuvant alters lymph flow and modulates adaptive immunity. Sci Immunol 6, eabf1152 (2021). <10.1126/sciimmunol.abf1152>

138. S. H. Bhagchandani, et al., Two-dose priming immunization amplifies humoral immunity by synchronizing vaccine delivery with the germinal center response. Sci Immunol 9, eadl3755 (2024). <10.1126/sciimmunol.adl3755>

