## Supplemental Information for "Consistent Induction of Broadly Neutralizing HIV Antibodies by a Novel Two-Step Mechanism Informs Immunogen Design"

#### Materials and Methods

Non-human primates

Indian rhesus macaques were housed at Bioqual, Inc., Rockville, MD, in accordance with guidelines of the Association for Assessment and Accreditation of Laboratory Animal Care standards. All experiments were approved by the University of Pennsylvania and Bioqual Institutional Animal Care and Use Committees. All incoming animals were healthy and naïve to HIV and SHIV exposure.

Animals in Group 1 were immunized with protein-nanoparticle versions of RC1-3fill (wk -16) and 11MUTB-3fill (wk -8). At each time point, a total dose of 100 ug of immunogen and 750 ug of SMNP adjuvant was given subcutaneously, divided evenly between the four limbs. Four macaques received bolus immunizations (V630, V631, V632, and V633), and four received escalating-dose immunizations (V634, V635, V636, and V637). The escalating dose regimen was administered over the course of two weeks, as described in ref (*103*). Animals in Group 2 were immunized with mRNA-LNP versions of RC1-3fill (wk -16) and 11MUTB-3fill (wk -8). A total dose of 100 ug of immunogen was given at each time point, with 25 ug administered subcutaneously in each upper limb and 25 ug administered intramuscularly in each lower limb.

At wk 0, all animals were infected with either SHIV.5MUT (Group 1-3) or SHIV.BG505.N332 (Group 4). All but two SHIV.5MUT-infected animals (AH82, AG99) received an intravenous infusion of 25 mg/kg of anti-CD8α (Nonhuman Primate Reagent Resource MT807R1) or anti-CD8β (Nonhuman Primate Reagent Resource CD8b255R1) 2-3 days prior to SHIV inoculation to transiently deplete CD8 T cells, with the goal of facilitating viral replication *in vivo* **(Supplementary Table 1)**. All SHIV.5MUT infections were performed intravenously. The SHIV.5MUT inoculum was generated by transfecting 293T cells with the respective infectious molecular clone (50 ng of p27 antigen in RPMI1640 with 10% heat-inactivated fetal bovine serum (FBS)). SHIV.BG505.N332-infected animals were inoculated as described in ref (*48*).

Processing and storage of clinical samples

Blood samples were collected from immunized and infected rhesus macaques into sterile vacutainers containing Anticoagulant Citrate Dextrose Solution A (ACD-A). 40 mL of anticoagulated blood was combined into a sterile 50 mL polypropylene conical tube and centrifuged at 1000 x g for 10 min at 20°C. The plasma phase was collected, clarified by centrifugation, aliquoted into 1 mL cryovials, and stored at -80°C. The WBC and RBC fractions were combined, resuspended in an equal volume of Hanks' Balanced Salt Solution (HBSS) without $Mg^{2+}$/$Ca^{2+}$ and containing 2 mM EDTA, and distributed evenly into four sterile 50 mL tubes. The volume of each tube was adjusted to 30mL with HBSS-EDTA (2mM), underlaid with 14 mL of 96% Ficoll-Paque (diluted with HBSS-EDTA), and centrifuged at 725 x g for 25 min at 20°C with slow acceleration and braking. Cells at the phase interface were collected, transferred to a sterile 50mL tube, diluted to a final volume of 50 mL with HBSS-EDTA (2mM), and centrifuged at 500 x g for 15 min at 20°C. The cell pellet was resuspended in 30 mL of HBSS with $Mg^{2+}$/$Ca^{2+}$ and 1% FBS, then centrifuged at 200 x g for 15 min at 20°C. If heavy RBC contamination was observed, lysis was performed using Biolegend RBC Lysis Buffer (420301). The cell pellet was resuspended in 30 mL of HBSS with $Mg^{2+}$/$Ca^{2+}$ and 1% FBS, and cell count and viability were determined with acridine orange / propidium iodide (AOPI) staining. Cells were centrifuged at 300 x g for 10 min at 20°C, resuspended at 5-10 × $10^6$ cells/mL in CryoStor CS5 cell cryopreservation media (Sigma C2999), and aliquoted into 1 mL cryovials (Thermo 374503).

Cells were stored in a CoolCell container (Corning 432008) at -80°C overnight and then transferred to vapor-phase liquid nitrogen for long-term storage. Cells collected from lymph nodes and bone marrow were processed in a similar manner. Lymph nodes were excised and immediately placed in RPMI1640 with 10% FBS on wet ice, cleaned of fat and connective tissue via dissection, quartered, and homogenized through a sterile 100 μm strainer. The homogenized cell suspension was transferred to a 50 mL tube, diluted to a final volume of 30 mL with RPMI with 10% FBS, and subjected to Ficoll density gradient purification as described above. Bone marrow was vigorously resuspended in 20 mL of HBSS without $Mg^{2+}/Ca^{2+}$, fat and connective tissue was removed, and the resulting single-cell suspension was purified by Ficoll density gradient purification as described above.

Protein expression and purification for nanoparticle preparations and structure determinations

Soluble SOSIP.664 versions (*53*) of BG505, 5MUT, del4, and del8 native-like gp140 Env trimers were expressed with MD39 stabilizing mutations (*24*) and potential N-linked glycosylation sites (PNGSs) introduced at positions $230_{gp120}$, $241_{gp120}$, and $344_{gp120}$ ("3fill" substitutions) to shield an immunodominant off-target glycan hole (*26, 69, 70*) as described (*104*). SpyTagged SOSIP Env trimers were prepared by adding a 16-residue SpyTag003 sequence (*105*) to the C-terminus of each soluble Env trimer. Soluble SOSIP Envs were expressed by transient transfection in Expi293 cells (Life Technologies). Proteins were harvested from culture supernatants and purified using PGT145 immunoaffinity chromatography followed by size-exclusion chromatography (SEC) on Superose 6 (GE Healthcare) in 20 mM sodium phosphate pH 7.5, 150 mM NaCl (PBS) as described (*104*). Final protein preparations were stored at 4°C. Fab fragments corresponding to the AJ09-21, AJ09-83, AJ09-110, AM12-340, AM12-347, AM12-351, AM12-352, NN39-25, V634-136, V634-136 UCA, V645-158, NN39-25, and NN39-171 IgGs were generated as described (*106*). In brief, Expi293 cells were transiently transfected with plasmids encoding an IgG light chain and the Fab region of an IgG heavy chain including a C-terminal 6x-His tag. Fabs secreted into the culture medium were isolated by nickel-nitrilotriacetic acid ($Ni^{2+}$-NTA) affinity chromatography (GE Healthcare) followed by purification via SEC using a Superdex 200 16/60 column (GE Healthcare). Purified Fabs were concentrated and stored at 4°C in a buffer containing 20 mM Tris pH 8.0, 150 mM NaCl (TBS) and 0.02% sodium azide.

Preparation of SOSIP-mi3 nanoparticles

SpyCatcher003-mi3 particles were prepared by purification from BL21 (DE3)-RIPL *E. coli* (Agilent) transformed with a pET28a SpyCatcher003-mi3 gene (*107*) (including an N-terminal 6x-His tag) as described (*108*). Briefly, bacterial cell pellets were lysed with a cell disruptor in the presence of 2.0 mM PMSF (Sigma) and then spun at 21,000 x g for 30 min and filtered with a 0.2 μm filter. mi3 particles were isolated by ammonium sulfate precipitation followed by SEC using a HiLoad 16/600 Superdex 200 (GE Healthcare) column equilibrated with TBS pH 8.0. SpyCatcher003-mi3 particles were stored at 4 °C and used for conjugations after filtering with a 0.2 μm filter and spinning for 30 min at 4 °C and 14,000 x g. For preparation of SOSIP-mi3 nanoparticles, SpyCatcher003-mi3 was incubated with a 2-fold molar excess (SOSIP protomer to mi3 subunit) of purified SpyTagged SOSIP overnight at room temperature in PBS. Conjugated SOSIP-mi3 particles were separated from free SOSIPs by SEC on a Superose 6 10/300 column (GE Healthcare) in PBS. Fractions corresponding to conjugated mi3 particles were identified by SDS-PAGE. Concentrations of SOSIP-mi3 particles were determined using the absorbance at 280 nm as measured on a Nanodrop spectrophotometer (Thermo Scientific).

mRNA design and production

Plasmids encoding membrane-anchored prefusion-stabilized Env trimers matching RC1 and 11MUTB protein immunogens were designed, cloned (Genscript), and used as a template for *in vitro* transcription (**fig. S1A**). Nucleoside-modified mRNA immunogens were produced using T7 RNA polymerase (Megascript) with co-transcriptional RNA capping (CleanCap 3OMe, Trilink) and incorporation of N1-methylpseudouridine in place of uridine (Trilink) as previously reported (*109, 110*). A plasmid-encoded polyadenylation sequence (101 nt) was incorporated to improve mRNA stability within the cell. Contaminating double stranded byproducts were removed with cellulose purification as previously described (*111*) and innate immune sensing was assessed as interferon-alpha release following transfection of monocyte-derived dendritic cells (*112*). mRNA was evaluated for appropriate size using gel electrophoresis and stored at -20°C.

mRNA-LNP Synthesis

mRNA-LNPs were prepared in 4-component lipid nanoparticles using previously described methods (*113, 114*). LNP formulations contain ionizable lipid (proprietary to Acuitas) / phosphatidylcholine / cholesterol / PEG-lipid. Lipid and LNP compositions are described in US patent US10,221,127.

SHIV and pseudovirus production

The construction of SHIV.BG505.N332 was described previously and includes an S375Y mutation that is required for efficient entry and replication in rhesus CD4 T cells (*115*). SHIV.5MUT was generated by mutagenizing four residues (V134Y, N136P, I138L, and D140N) in the plasmid encoding this parental strain using the Q5 Site-Directed Mutagenesis Kit (NEB E0554S). Mutants in this and other backbones were generated using the same strategy. Mutagenized plasmids were sequenced to confirm their integrity.

Infectious SHIV stocks were produced as previously described (*115*). Briefly, $5 \times 10^6$ HEK 293T cells (ATCC CRL-3216) were plated in a 100 mm tissue culture dish in 10 mL of Dulbecco's Modified Eagle Medium (DMEM) supplemented with 10% FBS and 100 U/mL penicillin-streptomycin (Gibco 10378-016) and cultured at 37°C with 5% $CO_2$. After 24 hours, 6 ug of SHIV plasmid DNA (for SHIV generation) or 4.5 ug of HIV-1 backbone (SG3ΔEnv) plasmid plus 1.5 ug of Env plasmid (for pseudovirus generation) was diluted in 500 uL of DMEM, combined with 18 uL of FuGENE 6 transfection reagent (Promega E2692), and added dropwise to the cells. Cells were cultured for 48 hours at 37°C with 5% $CO_2$. Virus-containing media was then harvested and centrifuged at 2000 x g for 8 min at 20°C. Supernatant was divided into 500 uL aliquots and stored at -80°C. SHIV infection stocks were quantified by measuring p27 antigen using an ELISA (ZeptoMetrix 0801169). Pseudovirus stocks were titrated in TZM-bl cells (*116*). Briefly, $1.5 \times 10^4$ TZM-bl cells were seeded in flat-bottom 96-well tissue culture plates in DMEM with 10% FBS and 100 U/mL penicillin-streptomycin and cultured at 37°C with 5% $CO_2$. Virus stocks were harvested 24 hours later, serially five-fold diluted in DMEM with 6% FBS, 100 U/mL penicillin-streptomycin, and 40 ug/mL DEAE-dextran, and then added to the cells with four technical replicates. 48 hours later, cells were fixed in phosphate-buffered saline (PBS) containing 0.8% glutaraldehyde and 2.2% formaldehyde for 10 min, washed three times with PBS, and stained with PBS containing 4 uM magnesium chloride, 4 uM, potassium ferricyanide, 4 uM potassium ferrocyanide, and 400 ug/mL X-gal for 3 hours at 37°C. Cells were washed three times with PBS and imaged on an ELISPOT analyzer (CTL ImmunoSpot 7.0.34.0 Professional Analyzer DC).

Infectious virus titers were determined by averaging the number of spots per virus dilution and dividing by the viral input volume.

Plasma viral RNA quantification

SHIV plasma viral load was quantified using standardized, quality-controlled real-time PCR assays (Applied Biosystems) at the CLIA-certified, NIH/NIAID-sponsored Non-Human Primate Virology Core Laboratory at the Duke Human Vaccine Institute. High-throughput sample processing and PCR setup was facilitated by use of QIAsymphony SP and QIAgility automation platforms (QIAGEN). Viral RNA was isolated from plasma and reverse-transcribed into cDNA using a target-specific primer. cDNA was treated with RNase, added to a custom real-time PCR master mix containing target-specific primers and a fluorescently-labeled hydrolysis probe, and amplified on a QuantStudio3 (Thermo) real-time quantitative PCR machine. Raw data were quality-controlled (including confirmation of positive and negative controls) and mean viral RNA copies/mL was calculated. The lower limit of quantification of this assay was 62 RNA copies/mL.

Neutralization assay

Neutralization assays were performed using TZM-bl indicator cells as previously described (*115-117*). Briefly, $1.5 \times 10^4$ TZM-bl cells were plated in flat-bottom 96-well plates in 100 uL of Dulbecco's Modified Eagle Medium (DMEM) supplemented with 10% FBS and 100 U/mL penicillin-streptomycin and cultured at 37°C with 5% $CO_2$. 24 hours later, plasma samples were heat-inactivated at 56°C for 1 hour and serially five-fold diluted starting at 1:20. To maintain a constant plasma concentration of 5% v/v across all wells, media was supplemented with 10% human serum (Millipore Sigma H5667) at all but the highest dilution. Alternatively, monoclonal antibodies were serially five-fold diluted starting at 50 or 250 ug/mL in media that was not supplemented with human serum. Virus was diluted to achieve a final multiplicity of infection (MOI) of 0.3 when added to the TZM-bl cells. Virus was incubated with plasma or mAb dilutions at 37°C for 1 hour, and this mixture was then added to the adherent TZM-bl cells and cultured at 37°C with 5% $CO_2$. 48 hours later, cells were lysed with 0.5% Triton-X 100 in PBS and luciferase activity was quantified using a luciferase assay reagent (Promega E1501) and a BioTek Synergy Neo2 plate reader (Agilent).

Generation of soluble Env trimers for flow cytometry

Stabilized HIV Env trimers were expressed and purified as previously described. A V3 knockout construct was generated by introducing D325A, R327A, and S334A mutations into the Q23 backbone. Plasmids encoding Env trimers with C-terminal AviTags were transfected into HEK293F cells (Thermo R79007) cultured in Gibco FreeStyle 293 Expression Medium (Gibco 12338018) using PEI-MAX 4000 transfection reagent (Polysciences 23765). *Escherichia coli* biotin ligase (BirA) was co-transfected to site-specifically biotinylate the biotin-acceptor AviTag peptide on Env. Cells were maintained at 37°C with 8% $CO_2$ and shaking at 125 rpm. Four days post-transfection, cultures were harvested by centrifugation at 2700 x g for 15 min and clarified supernatants containing secreted Env trimers were collected. Env trimers were purified by affinity chromatography using either agarose-bound Galanthus nivalis lectin (GNL; Vector Laboratories AL-1243-5) or Toyopearl AF-Tresyl-650M resin (Tosoh Bioscience 0014472) conjugated to the broadly neutralizing antibody PGT145. Antibody conjugation to the resin was performed according to the manufacturer's protocol. Eluted proteins were further purified by size-exclusion chromatography (SEC) using a Superdex 200 Increase 10/300 GL column (Cytiva 28-9909-44) equilibrated in PBS or Tris-buffered saline (TBS). Biolayer interferometry (BLI) was used to assess the efficiency of biotinylation with an Octet RH96 instrument (Sartorius). Biotinylated Env proteins were diluted in PBS supplemented with 0.1% Tween-20 and analyzed using streptavidin biosensors (Sartorius 18-5019) pre-equilibrated in the same buffer. Successful biotinylation was confirmed by a reproducible and saturable loading response within the expected range for mono-biotinylated proteins. Purified biotinylated Envs were conjugated to streptavidin-BV421 (Biolegend 405225), streptavidin-AF647 (Biolegend 405237), or streptavidin-PE (Invitrogen SA10044) by incubating together at a 4:1 molar ratio at 4°C.

<u>B cell isolation</u>

5-10 × $10^6$ cryopreserved PBMCs were thawed and resuspended in 10mL of RPMI supplemented with 10% FBS, 100 U/mL penicillin-streptomycin, and *** Benzonase (Millipore Sigma 70664-3). The cell suspension was centrifuged at 300 x g for 5 min at 4°C, washed once with PBS, and stained with Zombie-NIR viability dye (Biolegend 77184) for 30 min at 4°C. Cells were washed with PBS supplemented with 5% FBS and stained for 1 hr at 4°C with a cocktail of antibodies against CD3 (clone SP34-2, BD 557757), CD8a (clone SK1, Invitrogen 47-0087-41), CD14 (clone 61D3, Invitrogen 47-0149-42), CD16 (clone eBioCB16, Invitrogen 47-0168-41), CD20 (clone L27, BD 335793), and IgG (clone G18-145, BD 550931). For isolation of bulk B cells for lineage tracing analysis, antibodies against IgM (clone G20-127, BD 562618) and IgD (polyclonal, Dako F0189) were also used. For isolation of single, Env-specific B cells, cells were washed once with PBS supplemented with 5% FBS and then were stained for 1 hr at 4°C with 1 ug of BV421-, AF647, and/or PE-conjugated soluble Env trimer in PBS supplemented with 5% FBS. All cells were then washed three times with PBS supplemented with 5% FBS, filtered through a sterile 100 µm strainer, and sorted on a BD FACSMelody Cell Sorter. Naïve ($CD3^-CD8a^-CD14^-CD16^-CD20^+IgM^+IgD^+$) or memory ($CD3^-CD8a^-CD14^-CD16^-CD20^+IgM^-IgD^-IgG^+$) B cells were sorted in bulk into filtered FBS, centrifuged at 300 x g for 5 min, resuspended in 600 uL of cold RNAzol RT (Molecular Research Center RN190), transferred into 2 mL vials (Sarstedt 72.694.396), and stored at -80°C. Single, Env-specific B cells ($CD3^-CD8a^-CD14^-CD16^-CD20^+IgG^+Env^{++}$) were sorted into individual wells of a 96-well PCR plate containing 20 uL of lysis buffer containing 0.5 uL RNAse OUT (40 U/uL, Invitrogen 10777019), 5 uL 5X First Strand Buffer (Invitrogen 18080044), 1.25 uL DTT (0.1 M, Invitrogen 18080044), 0.0625 uL IGEPAL CA-630 detergent (Sigma I8896), and 13.25 uL nuclease-free water. Cell lysate was centrifuged at 300 x g for 5 min at 4°C and stored at -20°C. Bone marrow plasma cells were sorted in a similar manner, with the following changes. After viability staining cells were washed, resuspended in PBS supplemented with 5% FBS and 1:20 anti-FcR (Invitrogen 14-9156-42), and incubated for 15 min at 4°C. Without washing, cells were then stained with a cocktail of antibodies against CD3, CD8a, CD14, CD16, CD20, CD102 (clone CBR-IC2/2, Biolegend 328506), and CD31 (clone C31.7, Novus NBP2-33136AF647). Plasma cells ($CD3^-CD8a^-CD14^-CD16^-CD20^-CD102^+CD31^+$) were sorted into 20 uL of RPMI supplemented with 10% FBS and 100 U/mL penicillin-streptomycin, counted, and immediately carried forward for single-cell analysis using the 10X Genomics platform.

<u>Single-cell BCR sequencing from PBMCs</u>

Immunoglobulin heavy (IgH) and light (IgK, IgL) chain genes were amplified as previously described (*39*). Briefly, single B cell RNA was reverse transcribed by adding 2.3 uL random hexamers (200 ng/uL, Thermo SO142), 2 uL dNTP mix (10 mM per nucleotide, Invitrogen 18427088), 1 uL Superscript III reverse transcriptase (200 U/uL, Invitrogen 18080044), and 0.7

uL nuclease-free water and incubating in a thermocycler at 42°C for 10 min, 25°C for 10 min, 50°C for 1 hr, and then 94°C for 5 min. IgH, IgK, and IgL genes were amplified by two rounds of nested PCR, each using 3 uL of template, 5 uL 10X buffer (Qiagen 203207), 1uL dNTP mix (Qiagen 201901), 0.5 uL $MgCl_2$ (Qiagen 203207), 0.5 uL carrier RNA (Qiagen 1017647), 0.75 uL forward primer mix (50 uM per primer), 0.5 uL of each reverse primer (25 uM), 0.4 uL HotStart Taq polymerase (100 mM per nucleotide, Qiagen 203207), and 39.7 uL nuclease-free water. PCR conditions were as follows: 95°C for 15 min, then 50 cycles of 95°C for 30 sec, 51°C (Round 1) or 58°C (Round 2) for 30 sec, 72°C for 30 sec, with a final extension of 72°C for 7 min. Amplicons were visualized by agarose gel electrophoresis, Sanger sequenced (Genewiz), and analyzed computationally using IMGT V-QUEST (*118*).

Single-cell BCR sequencing from bone marrow-derived plasma cells

$CD102^+CD31^+$ bone marrow cells suspension was loaded on a Chromium X instrument (10X Genomics) to generate a single-cell bead emulsion, at a loading concentration for a target recovery of 20,000 cells per reaction. Single-cell RNA-seq libraries were then prepared using Chromium Next GEM Single Cell 5' Kit v3 bead and library construction kit (10X Genomics). BCR libraries were constructed using Chromium Single Cell Human BCR Amplification Kit (10X Genomics) and macaque-specific primers targeting the constant regions of heavy chain IgM, IgG, and IgA gene isotypes and light chain IgK and IgL genes, as described (*119*). BCR libraries were indexed using Dual Index Kit TT Set A kit (10X Genomics) and sequenced on an Illumina NextSeq 2000 instrument at a minimum read depth of 5000 reads/cell. Sequencing reads were analyzed using Cell Ranger v7.2 (10X Genomics) and heavy chain BCR sequences were recovered from a total of 21,220 cells. AM12-352 lineage members were identified based on CDRH3 length and amino acid sequence homology, heavy chain V, D and J gene usage, and light chain V and J gene usage.

Antibody cloning and expression

Immunoglobulin heavy and light chain variable regions of interest were synthesized and cloned (Genscript) into rhesus IgG1, IgK, or IgL expression vectors immediately upstream of the constant region using AgeI/NheI, AgeI/BsiWI, or AgeI/ScaI restriction sites, respectively, as previously described (*39*). Recombinant mAbs were expressed by co-transfecting paired heavy and light chain plasmids into 293F cells at a 1:2 ratio using ExpiFectamine transfection reagent (Gibco A14525), purified using rProtein A/Protein G Sepharose Gravitrap kit (Cytiva 28985256), concentrated in 30 kDa centrifugal filters (Sigma UFC903024), and buffer exchanged into PBS. Antibody concentration was measured fluorometrically using a Qubit Protein Assay kit (Invitrogen Q33212).

Bulk BCR sequencing and lineage tracing

Immunoglobulin heavy and light chain sequences were amplified and analyzed from bulk $IgM^+IgD^+$ or $IgG^+$ B cells as previously described (*48, 87, 120*). Briefly, RNA was extracted using RNAzol RT (Molecular Research Center RN190) according to the manufacturer's instructions. cDNA was synthesized using 5' rapid amplification of cDNA ends (5' RACE) with SMARTer template switching and SuperScript II reverse transcriptase (Invitrogen 18064071) and purified using AMPure XP beads (Beckman Coulter A63881). BCR library construction was performed as previously described (*48, 120*) using KAPA HiFi HotStart ReadyMix PCR Kit (Roche KK2601) and primers annealing in the IgG, IgK, or IgL constant region. Additional PCR steps were used to append barcodes and Illumina P5 and P7 sequencing adapters. All libraries were sequenced on an Illumina MiSeq sequencer (with 2 × 300 bp runs using the 600-cycle MiSeq Reagent V3 kit) or

NextSeq 2000 sequencer (with P1 or P2 600-cycle kits). Sequences from the naïve IgM$^+$IgD$^+$ B cell population were used to construct an individualized germline immunoglobulin repertoire for each macaque of interest using IgDiscover (*57, 121*) and MINING-D (*122*) computational pipelines, as previously described (*48*). Sequences from the IgG$^+$ B cell population were used to trace lineages of interest and infer unmutated common ancestors using the SONAR computational pipeline (*58*), as previously described (*48*).

Identification of homologous human V gene segment alleles

Homologous human V gene segment alleles were found using global alignments between the rhesus UCA V gene segment alleles and the orgdb database (https://ogrdb.airr-community.org/germline_sets/Human release data 2024-10-12) after removal of the novel alleles (designated with i) with BLOSUM62 scoring and gap opening and extension penalties of 11 and 1, respectively (*60*). The best human V gene segment allele was identified based on protein identity, and ties were broken using biochemical similarity.

Viral RNA sequencing

The SHIV *rev-vpu-env* cassette was sequenced using single-genome sequencing (SGS) approach as previously described (*61, 62*). Briefly, reverse transcription of RNA was performed by using reverse transcriptase (Superscript III) and reverse primer SIVmac251.R.R1 5′-CACTAGCTTACTTCTAAAATGGCAGC-3′ (nt 10,138–10163, SIVmac239) according to the manufacturer's instructions. The *rev-vpu-env* cassette was amplified by nested PCR with the following primers: first round forward primer SHIV.RevEnv.F1 5′-CGAAAGGCTTAGGCATCTCCTATG-3′ (nt 5949 - 5972, HXB2), second round forward primer SHIV.RevEnv.F2 5′-TTAGGCATCTCCTATGGCAGGAAGA-3′ (nt 5957-5981, HXB2), first round reverse primer SIVmac251.R.R1 5′-CACTAGCTTACTTCTAAAATGGCAGC-3′ (nt 10,138 - 10,163, SIVmac239), and second reverse primer SIVmac251.R.R2 5′-TACTTCTAAAATGGCAGCTTTATTGAAGAGG-3′ (nt 10,125 - 10,155, SIVmac239). A total of 8406 *env* gp140 sequences were aligned using the MUSCLE alignment method and manually inspected using Geneious Prime to improve the alignment results based on codon translation. The sequence alignment was then analyzed using Pixel plot (https://www.hiv.lanl.gov/content/sequence/pixel/pixel.html) and the LASSIE program (*80*).

Cryo-EM sample preparation

Purified Fabs and SOSIP Envs were incubated at a Fab:Env protomer molar ratio of ~4:1 at room temperature (for mature bNAb Fabs) or a ratio of ~1.1:1 at 37˚C (for V634-136 UCA Fab). Resulting mature Fab-Env complexes were purified by SEC using a Superdex 200 Increase 10/300 GL analytical column (GE Healthcare Life Sciences) in TBS pH 8.0, and fractions corresponding to a Fab-Env complex were concentrated to a final concentration of 2-3 mg/mL using a 50 kDa molecular weight cutoff spin concentrator (Millipore). The V634-136 UCA Fab-Env complex was not subjected to SEC purification.

Immediately prior to grid preparation, a fluorinated octyl maltoside solution (Anatrace) was added to each sample from a 0.5% (w/v) stock to a final concentration of 0.02% (v/v) as described (*123*), and 3 μL of each complex was applied to freshly glow-discharged Quantifoil R1.2/1.3 grids (300 mesh Cu; Electron Microscopy Sciences) that had been treated for 1 minute at 20 mA using a PELCO easiGlow device (Ted Pella). Grids were plunge-frozen using a Vitrobot Mark IV (Thermo Fisher) at 22°C and 100% humidity. Blotting was performed with Whatman

No. 1 filter paper for 3 s using a blot force of 0. Samples were vitrified by rapid plunging into liquid ethane cooled by liquid nitrogen.

Cryo-EM data collection

Single-particle cryo-EM data acquisition was performed on two transmission cryo-electron microscopes: A Titan Krios (Thermo Fisher) operating at 300 kV was used for imaging the AJ09-21, AJ09-83, and AJ09-110 Fabs in complex a SOSIP Env, and a Talos Arctica (Thermo Fisher) operating at 200 kV was used for the remaining Fab-Env complexes involving the AM12-340, AM12-347, AM12-351, AM12-352, NN39-25, V634-136, V634-136 UCA, V645-158, NN39-25, and NN39-171 Fabs. Automated data collection was carried out using SerialEM software (*124*), employing beam-image shift across a 3×3 grid of 1.2 μm holes with one exposure per hole. For the AJ09-21, AJ09-83 or AJ09-110–Env datasets recorded on the Krios, 40-frame movies were captured in super-resolution mode using a Gatan K3 camera positioned behind a BioQuantum energy filter (Gatan) with a 20 eV slit and a pixel size of 0.416 Å (105,000x magnification). For datasets collected on the Talos Arctica, 40-frame movies were captured in a super-resolution mode using Gatan K3 camera with a pixel size of 0.465Å (45,000x magnification). Data processing was carried out for all datasets using cryoSPARC v.4 (*125*). Data collection parameters are summarized in **Table S2**.

Cryo-EM data processing

Briefly, cryo-EM movies were patch motion corrected in cryoSPARC v4 (*125*) to account for beam-induced motion, including dose weighting, following the binning of super-resolution frames. For CTF parameter estimations, non-dose-weighted micrographs were processed using the Patch CTF job in cryoSPARC. Micrographs displaying poor CTF fits or evidence of crystalline ice in their power spectra were excluded from further analysis. Particle picking was performed in cryoSPARC v4 (*125*) using Blob picker (minimum particle diameter = 120 Å, maximum particle diameter = 180 Å) for reference-free selection. Particle extraction was carried out using a box size of 360 Å. Particles were subjected to several rounds of 2D classification. The best class averages representing different views were used to generate ab initio models, which were further refined using homogenous refinement (with minimize over per-particle scale) or non-uniform refinement by applying C1 or C3 symmetry. The particles and 3D volumes from homogenous refinement jobs combined with micrographs from the CTF estimation for reference-based motion correction before further refining using homogenous refinement or non-uniform refinement to generate the final refined 3D volume for all structures.

Cryo-EM structure modelling and refinement

To generate initial coordinates of the complex, individual chains from reference structures of the fully glycosylated BG505 SOSIP.664 HIV-1 Env trimer (PDB: 5T3X or 5T3Z) were independently docked into cryo-EM density maps using UCSF ChimeraX (*102*). Preliminary models were subjected to rigid-body fitting and subsequently refined in real space against the EM maps. Manual adjustments and sequence corrections were performed using Coot v.0.8.9 (*126*), followed by iterative cycles of model rebuilding in Coot and refinement using Phenix (*127*). N-linked glycans were incorporated at predicted glycosylation sites by interpreting low-resolution ("blurred") maps filtered with varying B-factors. Final models were assessed for stereochemical quality using MolProbity (*128*).

Structural analyses

Structure figures were created using ChimeraX (*102*) and PyMOL (*129*). To highlight Fab densities within EM reconstructions, maps were segmented, with each region visualized in a unique color. The gp41 region was rendered using a color scheme that corresponded to the fitted SOSIP coordinates from lower-resolution reconstructions. Estimations of local resolution were performed using cryoSPARC v4 (*125*). To quantify interfacial interactions, buried surface areas were analyzed via PDBePISA v1.48.65 (*130*), employing a 1.4 Å solvent probe radius. Hydrogen bonds were defined as interactions involving distances below 4.0 Å and an angle between donor, hydrogen, and acceptor atoms greater than 90°. Van der Waals contacts were considered when interatomic distances were under 4.0 Å. These interaction assessments are considered approximate due to limitations in resolution. Structural comparisons, including root-mean-square deviation (r.m.s.d.) values from pairwise Cα alignments, were calculated in PyMOL (*129*) without applying outlier exclusion. Criteria used to define epitope regions are provided in the respective figure legends.

The angles of approach of V3 Abs were calculated as described (*31*). The Env trimer 3-fold symmetry axis was aligned on the z axis, with the center of mass of one monomer's $N332_{gp120}$ epitope placed on the x axis. The $N332_{gp120}$ epitope was defined by residues $G324_{gp120}$, $D325_{gp120}$, $V326_{gp120}$, $R327_{gp120}$, $M328_{gp120}$, $A329_{gp120}$, $H330_{gp120}$, $I415_{gp120}$, $L416_{gp120}$, and $P417_{gp120}$. The center of mass of a gp41 residue ($L587_{gp41}$ in all three protomers) was aligned to the negative z axis. The latitudinal angle was that formed by the z axis and a vector from the $N332_{gp120}$ epitope center of mass to the HC center of mass (carbon-a atoms for six b-strands; HC residues 21-24, 34-39, 46-52, 67-71, 77-82, and 89-92) in the x-z plane. The longitudinal angle was that formed by the x axis and the $N332_{gp120}$ epitope-HC vector in the x-y plane. The HC-LC twist angle was the angle between the x axis and a vector from the LC center of mass (carbon-a atoms for six b-strands, LC residues 18-24, 34-37, 45-48, 62-66, 70-76, and 84-88) to the HC center of mass in the x-y plane.

Site-specific glycan analysis

100 µg aliquots of each sample were denatured for 1h in 50 mM Tris/HCl, pH 8.0 containing 6 M of urea and 5 mM dithiothreitol (DTT). Next, Env samples were reduced and alkylated by adding 20 mM iodoacetamide (IAA) and incubated for 1h in the dark, followed by a 1h incubation with 20 mM DTT to eliminate residual IAA. The alkylated Env samples were buffer exchanged into 50 mM Tris/HCl, pH 8.0 using Vivaspin columns (10 kDa) and two of the aliquots were digested separately overnight using Trypsin (Mass Spectrometry Grade, Promega), chymotrypsin (Promega), or alpha-lytic protease (Sigma Aldrich) at a ratio of 1:30 (w/w). The next day, the peptides were dried and extracted using an Oasis HLB µElution Plate (Waters).

The peptides were dried again, re-suspended in 0.1% formic acid, and analyzed by nanoLC-ESI MS with a Vanquish Neo (Thermo Fisher Scientific) system coupled to an Orbitrap Eclipse Tribrid mass spectrometer (Thermo Fisher Scientific) using stepped higher energy collision-induced dissociation (HCD) fragmentation. Peptides were separated using a µPAC Neo HPLC Column (180 µm × 110 cm). A trapping column (PepMap 100 C18 3µM 75µM × 2cm) was used in line with the LC prior to separation with the analytical column. The LC conditions were as follows: 280-minute linear gradient consisting of 4-32% acetonitrile in 0.1% formic acid over 260 minutes followed by 20 minutes of alternating 76% acetonitrile in 0.1% formic acid and 4% ACN in 0.1% formic acid, used to ensure all the sample had eluted from the column. The flow rate was set to 300 nL/min. The spray voltage was set to 2.5 kV and the temperature of the heated capillary was set to 55 °C. The ion transfer tube temperature was set to 275 °C. The scan range was 350−2000 m/z. The stepped HCD collision energies were set to 15, 25 and 45% and the MS2 for each energy was combined. Precursor and fragment detection was performed using an Orbitrap at a resolution $MS^1$ = 120,000, $MS^2$ = 30,000. A standard AGC target for $MS^1$ ($4e^5$) and $MS^2$ ($1e^4$) and auto injection times ($MS^1$ =50ms $MS^2$ =54ms) were used.

Glycopeptide fragmentation data were extracted from the raw file using Byos (Version 5.5; Protein Metrics Inc.). The glycopeptide fragmentation data were evaluated manually for each glycopeptide; the peptide was scored as true-positive when the correct b and y fragment ions were observed along with oxonium ions corresponding to the glycan identified. The MS data was searched using the Protein Metrics 305 N-glycan library. The relative amounts of each glycan at each site as well as the unoccupied proportion were determined by comparing the extracted chromatographic areas for different glycotypes with an identical peptide sequence. All charge states for a single glycopeptide were summed. The precursor mass tolerance was set at 4 ppm and 10 ppm for fragments. A 1% false discovery rate (FDR) was applied. The relative amounts of each glycan at each site as well as the unoccupied proportion were determined by comparing the extracted ion chromatographic areas for different glycopeptides with an identical peptide sequence. Glycans were categorized according to the composition detected.

HexNAc(2)Hex(9−3) was classified as M9 to M3. Any of these structures containing a fucose were categorized as FM (fucosylated mannose). Complex-type glycans were classified according to the number of HexNAc subunits and the presence or absence of fucose. Core glycans refer to truncated structures smaller than M3. As this fragmentation method does not provide linkage information, compositional isomers are grouped.

To obtain data for sites that frequently present low intensity glycopeptides, the glycans present on the glycopeptides were homogenized to boost the intensity of these peptides. This analysis loses fine processing information but enables the ratio of high mannose: complex: unoccupied to be determined. The remaining glycopeptides were first digested with Endo H (New England Biolabs) to deplete oligomannose- and hybrid-type glycans and leave a single GlcNAc or GlcNAcFuc residue at the corresponding site. The reaction mixture was then dried completely and resuspended in a mixture containing 50 mM ammonium bicarbonate and PNGase F (New England Biolabs) using only $H_2O^{18}$ (Sigma-Aldrich) throughout. This second reaction cleaves the remaining complex-type glycans but leaves the GlcNAc residues remaining after Endo H cleavage intact. The use of $H_2O^{18}$ in this reaction enables complex glycan sites to be differentiated from unoccupied glycan sites as the hydrolysis of the glycosidic bond by PNGaseF leaves a heavy oxygen isotope on the resulting aspartic acid residue. The resultant peptides were purified as outlined above and subjected to reverse-phase (RP) nanoLC-MS. Instead of the extensive N-glycan library used above, three modifications were searched for: +203 Da corresponding to a single GlcNAc, or +349 corresponding to a GlcNAcFuc, a remnant of an oligomannose/hybrid glycan (+/- fucose), and +3 Da corresponding to the $O^{18}$ deamidation product of a complex glycan. Data acquisition and analysis was performed as above and the relative amounts of each glycoform were determined, including unoccupied peptides.

Longitudinal Env evolution analyses

Our previously established algorithm LASSIE was used to identify Env sites under natural selection in each RM using the threshold of ≥80% TF loss at any timepoint (*80*), as was done previously for SHIV-infected macaques (*87*). To identify selected Env sites significantly enriched in SHIV.5MUT-infected macaques that developed bNAbs, the contingency tables of number of macaques that developed or failed to develop bNAbs with or without a site under selection were subjected to Fisher's exact tests and all significant associations (with uncorrected $p < 0.05$) were identified. All significantly enriched sites also met Benjamini-Hochberg multiple test correction with false discovery rate (FDR) < 20%. Hypervariable V1 loop characteristics of length and number of glycans were also tested for statistical enrichment using maximum increase in each characteristic's timepoint average over that of TF for each macaque. For calculation of hypervariable V1 loop number of PNGS for a given sequence, if two sequons were separated by a Pro (e.g. NNTPNAT), only one was assumed to be glycosylated based on 5MUT site-specific glycan analyses (**Fig. 4E**). Mutations at LASSIE selected sites and V3 loop sites were visualized using the Python software Logomaker (*131*).

Env-bNAb coevolution analysis

To assess whether plasma recognition of heterologous viruses required quasispecies mutation away from the infecting SHIV.5MUT (termed transmitted founder or TF virus) at key sites, we determined how often mutations emerged in the viral quasispecies at these sites prior to or contemporaneously with the earliest observed neutralization of each heterologous virus. Of the ten key sites identified as being preferentially selected in macaques that developed bNAbs (15, 87, 308, 325, 330, 332, 343, 359, 363, and 440), we determined the subset relevant to recognition of a particular heterologous virus on a per-animal basis. We assumed that the bNAb lineage in a given animal would select the most relevant resistance sites, and thus the subsets of sites differ between animals. For each heterologous virus in our screening panel, we only analyzed sites that differed between that virus and SHIV.5MUT, assuming that variation at those sites must be acquired before the bNAb lineage can recognize that virus. Additionally, sites 15 and 343 were not considered to be relevant to recognition of viruses encoding M-group consensus residues Q15 or Q343, as these residues were never sampled by the quasispecies in any SHIV.5MUT-infected macaque that developed bNAbs and thus are unlikely to be associated with resistance.

We determined the earliest timepoint at which mutations were observed at each of the relevant key sites in the evolving quasispecies of a given animal, even if the mutant was rare at that timepoint. While the mutant residue often matched that carried by the heterologous virus, we interpreted any variant residue as potentially contributing to recognition. This is particularly pertinent for position 363, which is a PNGS in SHIV.5MUT but is highly variable in the heterologous virus panel. The loss of this glycan in the quasispecies, which can be achieved by mutation to any non-asparagine residue at 363, is likely more important than the particular residue sampled. Similarly, while the heterologous viruses exhibit high variability at position 440, none of them carry an arginine. Nevertheless, Q440R was by far the most common escape mutation observed in the quasispecies of SHIV.5MUT-infected macaques that developed bNAbs, suggesting that the loss of the TF glutamine was more important than accommodation of a particular residue at this site. An example is shown in **Fig. 6A**, where nine of the ten key sites are under selection in animal V634. Only six of these differ between SHIV.5MUT and T250, including Q15 and Q343 **(Fig. 7A)**. So only the remaining 4 sites (325, 330, 363, and 440) were considered to be relevant for the bNAb lineage in this animal to recognize T250 **(Fig. 7B)**. Consistent with our hypothesis, mutations emerged at all 4 of these sites before or at the timepoint at which plasma neutralization of T250 was first detected. Indeed, in animal V634 this was the case for 20 of 21 residues across all heterologous viruses and relevant selected sites. The one exception was for site 440 and recognition of 25710, where the plasma acquired neutralization activity against 25710 at week 20 even though mutations at site 440 were not detected until week 24. A summary of all results is shown in **Fig. 7C-D**, and the individual analyses for each macaque are shown in **fig. S18-31**.

Given that increased V1h length and number of glycans were also significantly enriched in the quasispecies of SHIV.5MUT-infected macaques that developed bNAbs, we queried their contribution to the acquisition of plasma neutralization breadth in each macaque. V1h elongation was considered a key relevant feature for recognition of six of eight heterologous viruses that had longer V1h regions than SHIV.5MUT (>13 AA) **(Fig. 7A)**. In addition, all eight heterologous viruses had more V1h glycans than SHIV.5MUT (>1 glycan) **(Fig. 7A)**, and so an increased number of V1h PNGS was considered a relevant key feature for all viruses. In our analyses, we thus assumed that any increase in these parameters selected for antibodies capable of tolerating longer, more-glycosylated V1h regions. Indeed, the number of V1h glycans began to increase early in most SHIV.5MUT-infected macaques, and variants with >1 PNGS were observed in the quasispecies of animal V634 by week 4 post-infection **(Fig. 7B)**. V1h elongation was observed in the quasispecies at week 32, contemporaneous with plasma neutralization of 2/6 of the heterologous viruses for which V1h length is relevant (CAP256SU and BJOX02000). While there was low-titer neutralization of the other 4 viruses (T250, Ce1176, TRO11, and CH119) prior to this timepoint, we observed a substantial increase in potency upon emergence of the V1-elongated variants, suggesting that these variants may have selected for bNAb lineage members with enhanced affinity.

<u>Statistical analyses</u>

All statistical tests were calculated in GraphPad Prism 10 (version 10.4.2) or using the Stats module from SciPy (version 0.18.0) (*132*).

#### Supplementary Text

<u>Immunogen design</u>

To retain immunogen Envs in their prefusion conformation, we stabilized protein and mRNA constructs using MD39 mutations (*24*) and introduced potential N-linked glycosylation sites (PNGS) at gp120 positions 230, 241, and 344 (“3fill”) to shield immunodominant off-target epitopes (*27, 69, 70*). Envs expressed by mRNA constructs were further engineered to include an HLA-DR signal peptide (*133*), a flexible $(GGGGS)_2$ linker in place of the furin cleavage site (*134*), two helix-breaking prolines (*135*), and a truncated SIVmac cytoplasmic tail with an endocytosis knockout mutation (*136*) **(fig. S1A)**. Both RC1-3fill and 11MUTB-3fill mRNA constructs expressed well on the surface of 293F cells and exhibited favorable antigenic profiles **(fig. S1B)**. The protein immunogens were multimerized on mi3 nanoparticles (*99, 104*) and delivered with saponin/MPLA nanoparticle (SMNP) (*137*) adjuvant as either a bolus (n=4) or an escalating dose regimen over the course of two weeks (n=4) (*103, 138*). The mRNA immunogens were encapsulated in lipid nanoparticles (LNPs) and delivered as a bolus.

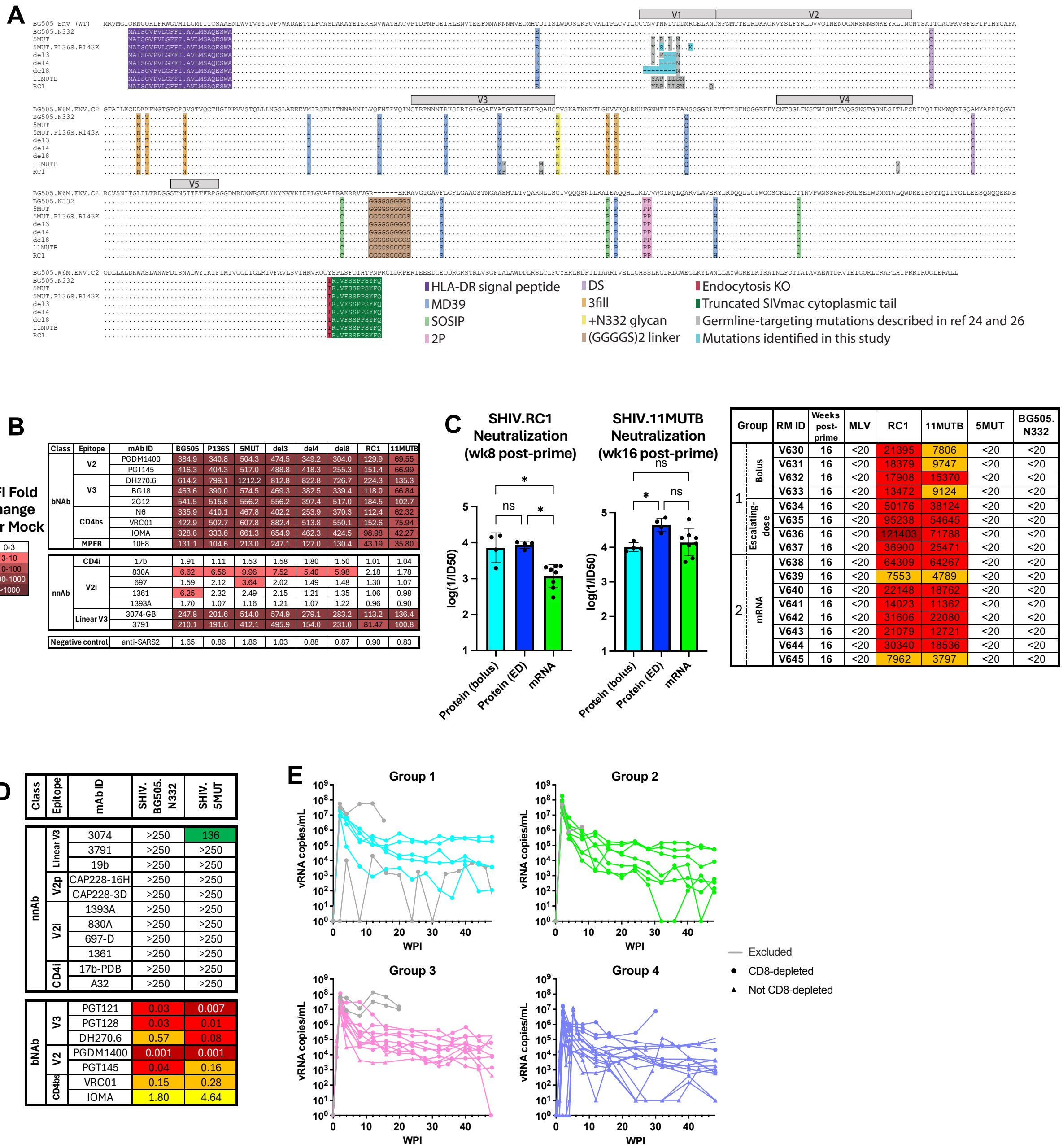


**Fig. S1. RC1 and 11MUTB immunization induces robust autologous neutralizing responses.** **(A)** mRNA constructs used in this study. Amino acid sequences are aligned to the BG505 wildtype Env sequence (dots indicate identity and dashes indicate deletions). Stabilization, glycan filling, and germline-targeting mutations are indicated. The RC1 and 11MUTB mRNA constructs were used as immunogens in this study **(Fig. 1A)**. **(B)** Antigenicity of all mRNA-expressed, stabilized, membrane-bound Envs shown in **(A)**. mRNA was transfected into 293F cells and expressed Envs were tested by flow cytometric analysis for binding to a panel of bNAbs (top) as well as conformation-sensitive non-neutralizing antibodies (nnAb, bottom). Binding was quantified as fold-change in mean fluorescence intensity (MFI) over mock-transfected cells, and data points with a >3-fold increase are colored as indicated. **(C)** Plasma neutralization of the indicated virus at 8 and 16 weeks post RC1 immunization, represented as log(1/$ID_{50}$) (left) or reciprocal $ID_{50}$

(right). Protein was given as bolus or escalating dose (ED). Kruskal-Wallis test with Dunn's correction for multiple comparisons. Not significant (ns) $P > 0.05$; $^{*}P < 0.05$. **(D)** Antigenicity of SHIV.5MUT and SHIV.BG505.N332. Neutralization sensitivity was tested using a panel of conformation-sensitive non-neutralizing antibodies (nnAb) and broadly neutralizing antibodies (bNAb). Titers are expressed as $IC_{50}$ in µg/mL. **(E)** Longitudinal viral load in macaques infected with SHIV.5MUT (Groups 1-3) or SHIV.BG505.N332 (Group 4), represented as vRNA copies/mL. Circles indicate animals that were CD8-depleted 2-3 days prior to infection, and triangles indicate animals that were not. Grey lines indicate animals that were excluded from subsequent analysis due to rapid progression to AIDS or abnormally low viral load. WPI, weeks post-infection.

| Group | RM ID | Weeks post-infection | Weeks post-imm | MLV | Tier-2 Autologous: 5MUT A1 | 5MUT. S334A A1 | 5MUT. D325N A1 | BG505 A1 | Tier-2 Heterologous: T250 02_AG | CAP256 SU C | Q23 A1 | 25710 C | BJOX02000 07_BC | Ce1176 C | TRO11 B | CH119 07_BC |
|---|---|---|---|---|---|---|---|---|---|---|---|---|---|---|---|---|
| 1 | V631 | 20 | 36 | <20 | 898 | 1,908 | 894 | <20 | <20 | <20 | <20 | <20 | <20 | <20 | <20 | <20 |
| | | 32 | 48 | <20 | 505 | 1,273 | 1,068 | 90 | <20 | <20 | <20 | <20 | <20 | <20 | <20 | <20 |
| | | 40 | 56 | <20 | 879 | 1,247 | 362 | 101 | <20 | <20 | <20 | <20 | <20 | <20 | <20 | <20 |
| | | 44 | 60 | <20 | 903 | 1,016 | 464 | 158 | <20 | <20 | <20 | <20 | <20 | <20 | <20 | <20 |
| | | 48 | 64 | <20 | 639 | 933 | 369 | 40 | <20 | <20 | <20 | <20 | <20 | <20 | <20 | <20 |
| | V632 | 20 | 36 | <20 | 1,267 | 1,637 | 772 | 134 | <20 | <20 | <20 | <20 | <20 | <20 | <20 | <20 |
| | | 32 | 48 | <20 | 1,844 | 685 | 2,247 | 646 | <20 | <20 | <20 | <20 | <20 | <20 | <20 | <20 |
| | | 40 | 56 | <20 | 3,666 | 895 | 2,279 | 216 | <20 | <20 | <20 | <20 | <20 | <20 | <20 | <20 |
| | | 44 | 60 | <20 | 1,834 | 511 | 1,300 | 304 | <20 | <20 | <20 | <20 | <20 | <20 | <20 | <20 |
| | | 48 | 64 | <20 | 2,838 | 265 | 1,814 | 78 | <20 | <20 | <20 | <20 | <20 | <20 | <20 | <20 |
| | V633 | 20 | 36 | <20 | 5,750 | 2,366 | 5,133 | 450 | <20 | <20 | <20 | 48 | <20 | <20 | <20 | <20 |
| | | 32 | 48 | <20 | 3,077 | 1,187 | 2,692 | 691 | 24 | 37 | 368 | 225 | 28 | 43 | 85 | 100 |
| | | 40 | 56 | <20 | 2,677 | 1,380 | 2,562 | 416 | 85 | 32 | 163 | 99 | 25 | <20 | 32 | 55 |
| | | 44 | 60 | <20 | 2,723 | 1,342 | 1,443 | 567 | 82 | 23 | 230 | 127 | 34 | <20 | 43 | 100 |
| | | 48 | 64 | <20 | 1,998 | 1,032 | 1,148 | 568 | 91 | <20 | 244 | 37 | 53 | <20 | <20 | <20 |
| | | 65 | 81 | <20 | 657 | 572 | 494 | 298 | 33 | 20 | 163 | <20 | 28 | <20 | <20 | <20 |
| | | 74 | 90 | <20 | 495 | 461 | 392 | 225 | 26 | <20 | 298 | <20 | 37 | <20 | <20 | <20 |
| | V634 | 20 | 36 | <20 | 6,406 | 8,525 | 2,630 | 919 | <20 | <20 | <20 | 33 | <20 | <20 | <20 | <20 |
| | | 24 | 40 | <20 | 2,658 | 4,591 | 1,707 | 663 | 29 | <20 | 99 | 38 | <20 | <20 | <20 | <20 |
| | | 28 | 44 | <20 | 3,828 | 5,063 | 2,008 | 670 | 97 | <20 | 326 | 277 | <20 | 50 | 138 | 54 |
| | | 32 | 48 | <20 | 4,092 | 8,000 | 3,224 | 1,724 | 239 | 22 | 887 | 629 | 69 | 98 | 422 | 93 |
| | | 36 | 52 | <20 | 2,995 | 5,621 | 2,572 | 1,393 | 407 | 66 | 590 | 431 | 108 | 204 | 628 | 120 |
| | | 40 | 56 | <20 | 6,402 | 7,102 | 3,575 | 2,459 | 542 | 64 | 1,241 | 743 | 137 | 202 | 1,076 | 160 |
| | | 44 | 60 | <20 | 4,914 | 3,614 | 2,928 | 1,109 | 783 | 42 | 1,590 | 1,117 | 193 | 238 | 803 | 122 |
| | | 48 | 64 | <20 | 5,780 | 5,107 | 3,855 | 1,631 | 1,092 | 29 | 2,094 | 1,110 | 142 | 291 | 913 | 278 |
| | | 65 | 81 | <20 | 953 | 5,522 | 1,081 | 861 | 491 | 27 | 435 | 105 | 70 | 40 | 183 | 40 |
| | | 74 | 90 | <20 | 952 | 3,894 | 1,054 | 573 | 209 | <20 | 265 | 92 | 47 | 24 | 91 | 35 |
| | V635 | 20 | 36 | <20 | 6,064 | 10,416 | 2,698 | 657 | <20 | <20 | <20 | <20 | <20 | <20 | <20 | <20 |
| | | 24 | 40 | <20 | 2,340 | 6,042 | 1,576 | 478 | <20 | <20 | <20 | <20 | <20 | <20 | <20 | <20 |
| | | 28 | 44 | <20 | 6,472 | 12,101 | 853 | 365 | <20 | <20 | 36 | <20 | <20 | <20 | <20 | <20 |
| | | 32 | 48 | <20 | 7,364 | 10,476 | 1,285 | 585 | <20 | 22 | 135 | <20 | 20 | <20 | <20 | <20 |
| | | 40 | 56 | <20 | 7,042 | 7,593 | 3,328 | 428 | <20 | 120 | 1,225 | <20 | 474 | 58 | 128 | 116 |
| | | 44 | 60 | <20 | 7,593 | 5,263 | 1,716 | 538 | <20 | 324 | 1,981 | 45 | 878 | 93 | 198 | 260 |
| | | 48 | 64 | <20 | 4,202 | 6,658 | 3,229 | 870 | 92 | 297 | 1,689 | 48 | 498 | 117 | 181 | 252 |
| | | 65 | 81 | <20 | 6,519 | 3,815 | 1,410 | 723 | 104 | 211 | 700 | 62 | 338 | 35 | 112 | 71 |
| | | 74 | 90 | <20 | 2,465 | 3,914 | 1,825 | 391 | 41 | 130 | 497 | 44 | 577 | 56 | 105 | 125 |
| 2 | V638 | 20 | 36 | <20 | 2,275 | 3,441 | 1,826 | 546 | <20 | <20 | <20 | <20 | <20 | <20 | <20 | <20 |
| | | 32 | 48 | <20 | 2,350 | 1,601 | 1,923 | 815 | <20 | <20 | <20 | <20 | <20 | <20 | <20 | <20 |
| | | 40 | 56 | <20 | 1,300 | 343 | 2,513 | 354 | <20 | <20 | <20 | <20 | <20 | 31 | <20 | <20 |
| | | 44 | 60 | <20 | 1,843 | 531 | 1,754 | 406 | <20 | <20 | <20 | <20 | <20 | 52 | <20 | <20 |
| | | 52 | 68 | <20 | 470 | 426 | 366 | 34 | <20 | <20 | <20 | <20 | <20 | <20 | <20 | <20 |
| | | 65 | 81 | <20 | 754 | 420 | 292 | 60 | <20 | <20 | <20 | <20 | <20 | <20 | <20 | <20 |
| | | 74 | 90 | <20 | 293 | 349 | 228 | 70 | 24 | <20 | <20 | 33 | <20 | <20 | <20 | <20 |
| | V639 | 20 | 36 | <20 | 2,900 | 5,501 | 1,984 | 455 | 30 | <20 | 33 | <20 | <20 | <20 | <20 | <20 |
| | | 32 | 48 | <20 | 6,993 | 8,354 | 4,088 | 1,326 | 34 | 46 | 127 | <20 | <20 | <20 | <20 | <20 |
| | | 40 | 56 | <20 | 3,765 | 5,467 | 6,075 | 852 | 64 | <20 | 97 | <20 | <20 | <20 | <20 | <20 |
| | | 44 | 60 | <20 | 3,220 | 3,885 | 3,101 | 251 | 71 | 132 | 402 | <20 | <20 | <20 | <20 | <20 |
| | | 52 | 68 | <20 | 2,896 | 4,490 | 4,082 | 249 | 20 | <20 | 126 | <20 | <20 | <20 | <20 | <20 |
| | | 56 | 72 | <20 | 3,514 | 6,684 | 5,345 | 978 | <20 | <20 | 233 | <20 | <20 | <20 | <20 | <20 |
| | | 65 | 81 | <20 | 2,160 | 3,617 | 1,735 | 701 | 50 | <20 | 591 | <20 | 26 | <20 | <20 | <20 |
| | | 74 | 90 | <20 | 2,459 | 4,831 | 1,421 | 463 | 30 | <20 | 251 | 23 | 26 | <20 | <20 | <20 |
| | V640 | 20 | 36 | <20 | 5,510 | 7,570 | 4,023 | 1,455 | <20 | <20 | <20 | <20 | <20 | <20 | <20 | <20 |
| | | 32 | 48 | <20 | 5,615 | 3,808 | 3,908 | 1,220 | <20 | <20 | <20 | <20 | <20 | <20 | <20 | <20 |
| | | 40 | 56 | <20 | 4,230 | 2,068 | 4,671 | 986 | <20 | <20 | 75 | <20 | <20 | 26 | 40 | <20 |
| | | 44 | 60 | <20 | 4,496 | 2,067 | 2,543 | 421 | <20 | <20 | 41 | <20 | <20 | <20 | <20 | <20 |
| | | 52 | 68 | <20 | 2,367 | 953 | 2,088 | 174 | <20 | <20 | 45 | <20 | <20 | <20 | <20 | <20 |
| | | 65 | 81 | <20 | 1,087 | 954 | 854 | 446 | <20 | <20 | 95 | <20 | <20 | <20 | <20 | <20 |
| | | 74 | 90 | <20 | 1,001 | 789 | 552 | 195 | <20 | <20 | 50 | <20 | <20 | <20 | <20 | <20 |
| | V641 | 20 | 36 | <20 | 2,416 | 3,478 | 4,202 | 128 | <20 | <20 | <20 | <20 | <20 | <20 | <20 | <20 |
| | | 24 | 40 | <20 | 1,754 | 6,329 | 3,499 | 110 | 30 | <20 | <20 | 26 | <20 | <20 | <20 | <20 |
| | | 28 | 44 | <20 | 3,319 | 8,150 | 3,471 | 191 | 81 | 23 | 63 | 56 | <20 | 24 | 28 | <20 |
| | | 32 | 48 | <20 | 2,259 | 3,791 | 2,901 | 220 | 27 | 47 | 152 | 422 | <20 | <20 | 33 | 24 |
| | | 40 | 56 | <20 | 4,769 | 1,132 | 3,428 | 444 | 36 | 826 | 2,257 | 1,178 | <20 | 131 | 192 | 153 |
| | | 44 | 60 | <20 | 6,601 | 9,276 | 9,141 | 401 | 399 | 1,789 | 2,038 | 1,420 | <20 | 375 | 185 | 611 |
| | | 52 | 68 | <20 | 2,863 | 1,364 | 3,289 | 396 | 563 | 177 | 424 | 344 | <20 | 76 | 72 | 38 |
| | | 56 | 72 | <20 | 1,992 | 2,380 | 2,886 | 298 | 1,834 | 86 | 766 | 245 | <20 | 23 | 50 | 43 |
| | | 65 | 81 | <20 | 2,885 | 2,042 | 8,489 | 328 | 2,683 | 165 | 1,667 | 110 | 50 | 54 | 21 | 53 |
| | | 74 | 90 | <20 | 3,105 | 945 | 4,655 | 190 | 3,891 | 457 | 805 | 193 | 42 | 88 | 63 | 110 |
| | V642 | 20 | 36 | <20 | 2,741 | 3,627 | 1,075 | 502 | <20 | <20 | <20 | <20 | <20 | <20 | <20 | <20 |
| | | 32 | 48 | <20 | 2,846 | 5,999 | 1,153 | 509 | <20 | <20 | <20 | <20 | <20 | <20 | <20 | <20 |
| | | 40 | 56 | <20 | 1,330 | 7,052 | 6,325 | 718 | <20 | <20 | <20 | <20 | <20 | <20 | <20 | <20 |
| | | 44 | 60 | <20 | 4,188 | 7,315 | 1,089 | 725 | <20 | <20 | <20 | <20 | <20 | <20 | <20 | <20 |
| | V644 | 20 | 36 | <20 | 508 | 661 | 356 | 50 | <20 | <20 | <20 | <20 | <20 | <20 | <20 | <20 |
| | | 32 | 48 | <20 | 404 | 1,055 | 453 | 42 | <20 | <20 | <20 | <20 | <20 | <20 | <20 | <20 |
| | | 40 | 56 | <20 | 182 | 449 | 398 | 43 | <20 | <20 | <20 | <20 | <20 | <20 | <20 | <20 |
| | | 44 | 60 | <20 | 455 | 669 | 228 | 24 | <20 | <20 | <20 | <20 | <20 | <20 | <20 | <20 |
| | V645 | 12 | 28 | <20 | 695 | 710 | 217 | 42 | <20 | <20 | <20 | <20 | <20 | <20 | <20 | <20 |
| | | 20 | 36 | <20 | 10,621 | 12,311 | 9,970 | 223 | <20 | <20 | 360 | <20 | <20 | <20 | <20 | <20 |
| | | 24 | 40 | <20 | 8,467 | 10,724 | 11,025 | 229 | 27 | 21 | 237 | <20 | <20 | <20 | <20 | <20 |
| | | 28 | 44 | <20 | 16,461 | 16,532 | 13,378 | 489 | 374 | 85 | 377 | 21 | <20 | <20 | <20 | <20 |
| | | 32 | 48 | <20 | 12,561 | 17,153 | 19,759 | 1,369 | 425 | 288 | 551 | 42 | 56 | <20 | <20 | <20 |
| | | 40 | 56 | <20 | 11,430 | 12,229 | 9,690 | 1,365 | 1,508 | 403 | 845 | 82 | 165 | 37 | 122 | 24 |
| | | 44 | 60 | <20 | 11,947 | 13,646 | 17,699 | 494 | 2,465 | 1,385 | 1,769 | 148 | 402 | 96 | 272 | 79 |
| | | 52 | 68 | <20 | 6,623 | 10,190 | 7,955 | 714 | 529 | 225 | 608 | 47 | 169 | 25 | 121 | 28 |
| | | 56 | 72 | <20 | 13,667 | 16,827 | 14,654 | 730 | 2,153 | 370 | 1,182 | 63 | 197 | 39 | 98 | 21 |
| | | 65 | 81 | <20 | 3,662 | 7,369 | 3,849 | 484 | 1,292 | 241 | 978 | 42 | 202 | 26 | 139 | 35 |
| | | 74 | 90 | <20 | 2,563 | 4,310 | 1,116 | 362 | 812 | 477 | 473 | <20 | 28 | 24 | 50 | <20 |
| 3 | V650 | 16 | N/A | <20 | 339 | 475 | 212 | <20 | <20 | <20 | <20 | <20 | <20 | <20 | <20 | <20 |
| | | 40 | N/A | <20 | 3,315 | 655 | 928 | 108 | <20 | <20 | 227 | <20 | <20 | <20 | <20 | <20 |
| | | 44 | N/A | <20 | 7,199 | 3,220 | 3,237 | 941 | <20 | <20 | 1,771 | <20 | 142 | <20 | <20 | <20 |
| | | 48 | N/A | <20 | 11,573 | 2,340 | 2,470 | 561 | <20 | 26 | 5,241 | <20 | 508 | 334 | 356 | 139 |
| | | 52 | N/A | <20 | 4,182 | 995 | 1,513 | 232 | <20 | <20 | 1,228 | <20 | 369 | 47 | 89 | 35 |
| | | 60 | N/A | <20 | 956 | 439 | 398 | 170 | <20 | <20 | 261 | <20 | 79 | 25 | 22 | <20 |
| | | 68 | N/A | <20 | 1,099 | 258 | 502 | 162 | <20 | <20 | 171 | <20 | 61 | 22 | 29 | <20 |
| | | 72 | N/A | <20 | 993 | 461 | 298 | 629 | <20 | <20 | 90 | <20 | 61 | <20 | <20 | <20 |
| | | 80 | N/A | <20 | 547 | 229 | 265 | 30 | <20 | <20 | 20 | <20 | 49 | <20 | <20 | <20 |
| | | 85 | N/A | <20 | 313 | 951 | 683 | 73 | <20 | <20 | 68 | <20 | <20 | <20 | <20 | <20 |
| | V651 | 16 | N/A | <20 | 4,062 | 1,677 | 1,529 | <20 | <20 | <20 | <20 | <20 | <20 | <20 | <20 | <20 |
| | | 40 | N/A | <20 | 15,788 | 7,634 | 9,425 | 32 | <20 | <20 | <20 | <20 | <20 | <20 | <20 | <20 |
| | | 44 | N/A | <20 | 16,469 | 8,718 | 10,047 | 36 | <20 | <20 | <20 | <20 | <20 | <20 | <20 | <20 |
| | | 48 | N/A | <20 | 13,721 | 8,826 | 10,571 | 64 | <20 | <20 | <20 | <20 | <20 | <20 | <20 | <20 |
| | | 52 | N/A | <20 | 15,444 | 7,843 | 8,251 | <20 | <20 | <20 | <20 | <20 | <20 | <20 | <20 | <20 |
| | | 60 | N/A | <20 | 10,801 | 7,072 | 8,453 | 36 | <20 | <20 | <20 | <20 | <20 | <20 | <20 | <20 |
| | | 68 | N/A | <20 | 8,425 | 7,278 | 7,740 | 39 | <20 | <20 | <20 | <20 | <20 | <20 | <20 | <20 |
| | | 72 | N/A | <20 | 16,258 | 4,815 | 8,244 | 44 | <20 | <20 | <20 | <20 | <20 | <20 | <20 | <20 |
| | | 80 | N/A | <20 | 5,112 | 1,857 | 3,720 | <20 | <20 | <20 | <20 | <20 | <20 | <20 | <20 | <20 |
| | | 85 | N/A | <20 | 6,605 | 4,596 | 4,198 | 121 | <20 | <20 | <20 | <20 | <20 | <20 | <20 | <20 |
| | CM03 | 16 | N/A | <20 | 3,196 | 1,493 | 1,622 | 486 | <20 | <20 | <20 | <20 | <20 | <20 | <20 | <20 |
| | | 20 | N/A | <20 | 3,459 | 935 | 2,184 | 158 | <20 | <20 | 32 | <20 | <20 | <20 | 25 | <20 |
| | | 24 | N/A | <20 | 3,033 | 1,115 | 2,091 | 418 | <20 | <20 | 91 | <20 | <20 | <20 | 44 | <20 |
| | | 28 | N/A | <20 | 2,259 | 1,223 | 2,022 | 180 | <20 | 23 | 65 | <20 | <20 | <20 | 37 | <20 |
| | | 32 | N/A | <20 | 2,409 | 998 | 1,746 | 387 | <20 | 31 | 100 | <20 | <20 | <20 | 40 | <20 |
| | | 36 | N/A | <20 | 1,970 | 1,459 | 3,478 | 177 | <20 | 31 | 43 | <20 | <20 | <20 | 44 | <20 |
| | | 40 | N/A | <20 | 1,800 | 2,131 | 2,386 | 380 | <20 | <20 | 37 | <20 | <20 | <20 | 31 | <20 |
| | | 44 | N/A | <20 | 2,225 | 1,648 | 2,821 | 209 | <20 | <20 | 49 | 20 | <20 | <20 | <20 | <20 |
| | | 48 | N/A | <20 | 1,368 | 1,311 | 2,664 | 62 | <20 | <20 | 23 | <20 | <20 | <20 | <20 | <20 |
| | | 52 | N/A | <20 | 1,970 | 1,459 | 3,478 | 177 | <20 | 31 | 43 | <20 | <20 | <20 | 44 | <20 |
| | | 56 | N/A | <20 | 521 | 558 | 878 | 42 | <20 | 73 | 169 | 26 | 26 | <20 | 41 | 27 |
| | | 60 | N/A | <20 | 217 | 266 | 570 | 26 | <20 | <20 | 24 | <20 | <20 | <20 | <20 | <20 |
| | CK36 | 16 | N/A | <20 | 115 | 152 | 125 | <20 | <20 | <20 | <20 | <20 | <20 | <20 | <20 | <20 |
| | | 20 | N/A | <20 | 433 | 276 | 148 | <20 | <20 | <20 | <20 | <20 | <20 | <20 | <20 | <20 |
| | | 24 | N/A | <20 | 419 | 267 | 307 | 26 | <20 | <20 | <20 | <20 | <20 | <20 | <20 | <20 |
| | | 28 | N/A | <20 | 535 | 453 | 532 | 99 | <20 | <20 | <20 | <20 | <20 | <20 | <20 | <20 |
| | | 32 | N/A | <20 | 891 | 944 | 1,108 | 347 | <20 | <20 | <20 | <20 | <20 | <20 | <20 | <20 |
| | | 36 | N/A | <20 | 721 | 1,131 | 1,136 | 705 | <20 | <20 | <20 | <20 | <20 | <20 | <20 | <20 |
| | | 40 | N/A | <20 | 1,402 | 1,318 | 1,096 | 448 | <20 | <20 | <20 | <20 | <20 | <20 | <20 | <20 |
| | | 44 | N/A | <20 | 876 | 703 | 951 | 323 | <20 | <20 | <20 | <20 | <20 | <20 | <20 | <20 |
| | | 48 | N/A | <20 | 903 | 724 | 667 | 158 | <20 | <20 | <20 | <20 | <20 | <20 | <20 | <20 |
| | | 52 | N/A | <20 | 721 | 1,131 | 1,136 | 705 | <20 | <20 | <20 | <20 | <20 | <20 | <20 | <20 |
| | | 56 | N/A | <20 | 547 | 638 | 611 | 118 | <20 | <20 | <20 | <20 | <20 | <20 | <20 | <20 |
| | | 60 | N/A | <20 | 457 | 487 | 617 | 229 | <20 | <20 | <20 | <20 | <20 | <20 | <20 | <20 |
| | NN58 | 16 | N/A | <20 | 6,394 | 7,174 | 3,979 | 174 | <20 | <20 | <20 | <20 | <20 | <20 | <20 | <20 |
| | | 20 | N/A | <20 | 4,726 | 8,190 | 3,381 | 1,268 | <20 | <20 | <20 | <20 | <20 | <20 | <20 | <20 |
| | | 24 | N/A | <20 | 9,681 | 5,429 | 4,021 | 1,192 | <20 | <20 | 122 | 26 | <20 | <20 | <20 | <20 |
| | | 28 | N/A | <20 | 7,536 | 5,914 | 5,063 | 1,934 | <20 | <20 | 550 | <20 | <20 | 26 | <20 | <20 |
| | | 32 | N/A | <20 | 6,341 | 4,502 | 2,869 | 1,092 | <20 | <20 | 611 | <20 | <20 | 35 | <20 | <20 |
| | | 36 | N/A | <20 | 3,628 | 4,011 | 2,406 | 912 | <20 | <20 | 350 | <20 | <20 | 43 | <20 | <20 |
| | | 40 | N/A | <20 | 3,554 | 2,437 | 2,064 | 518 | <20 | <20 | 195 | 30 | <20 | 41 | <20 | <20 |
| | | 44 | N/A | <20 | 2,481 | 2,045 | 2,244 | 627 | <20 | <20 | 701 | 34 | <20 | 67 | <20 | <20 |
| | | 48 | N/A | <20 | 2,430 | 1,695 | 1,790 | 255 | <20 | <20 | 451 | 48 | <20 | 68 | <20 | <20 |
| | | 52 | N/A | <20 | 3,628 | 4,011 | 2,406 | 912 | <20 | <20 | 350 | 54 | <20 | 82 | <20 | <20 |
| | | 56 | N/A | <20 | 1,619 | 1,996 | 2,251 | 220 | <20 | <20 | 218 | 43 | <20 | 31 | <20 | <20 |
| | | 60 | N/A | <20 | 903 | 867 | 861 | 113 | <20 | <20 | 140 | <20 | <20 | <20 | <20 | <20 |
| | NN39 | 16 | N/A | <20 | 4,699 | 5,435 | 4,854 | 482 | <20 | <20 | <20 | <20 | <20 | <20 | <20 | <20 |
| | | 20 | N/A | <20 | 11,101 | 11,380 | 7,716 | 1,218 | <20 | <20 | 42 | <20 | <20 | <20 | <20 | <20 |
| | | 24 | N/A | <20 | 13,891 | 6,969 | 5,640 | 1,310 | <20 | <20 | 1,436 | <20 | 126 | 51 | <20 | <20 |
| | | 28 | N/A | <20 | 6,636 | 3,614 | 5,734 | 1,290 | <20 | 38 | 1,169 | <20 | 200 | 63 | 40 | <20 |
| | | 32 | N/A | <20 | 8,071 | 4,699 | 16,340 | 1,721 | 32 | 354 | 2,404 | 20 | 980 | 481 | 426 | 395 |
| | | 36 | N/A | <20 | 13,161 | 5,444 | 35,714 | 1,323 | 193 | 1,000 | 4,630 | 139 | 3,448 | 1,680 | 1,082 | 1,380 |
| | | 40 | N/A | <20 | 9,025 | 4,785 | 55,648 | 1,413 | 307 | 171 | 1,723 | 106 | 493 | 216 | 411 | 317 |
| | AM12 | 12 | N/A | <20 | 52 | 113 | 55 | <20 | <20 | <20 | <20 | <20 | <20 | <20 | <20 | <20 |
| | | 16 | N/A | <20 | 375 | 277 | 176 | <20 | <20 | <20 | <20 | <20 | <20 | <20 | <20 | <20 |
| | | 24 | N/A | <20 | 7,364 | 572 | 1,633 | 3,472 | 65 | 558 | 923 | 53 | 125 | 73 | <20 | <20 |
| | | 32 | N/A | <20 | 4,352 | 2,077 | 2,859 | 721 | 347 | 536 | 696 | 353 | 202 | 369 | 184 | 35 |
| | | 36 | N/A | <20 | 13,495 | 2,149 | 5,373 | 1,069 | 5,464 | 2,332 | 2,189 | 1,465 | 741 | 552 | 227 | 124 |
| | | 40 | N/A | <20 | 4,836 | 1,196 | 3,515 | 607 | 3,034 | 1,308 | 1,179 | 581 | 345 | 321 | 172 | 52 |
| | | 44 | N/A | <20 | 9,398 | 2,817 | 6,435 | 846 | 5,602 | 2,471 | 2,255 | 1,400 | 995 | 623 | 446 | 250 |
| | | 48 | N/A | <20 | 1,856 | 180 | 1,482 | 356 | 2,413 | 1,362 | 730 | 517 | 391 | 336 | 114 | 81 |
| | | *51 | N/A | <20 | 1,513 | 694 | 775 | 119 | 1,412 | 345 | 407 | 238 | 159 | 171 | 48 | <20 |
| | AJ09 | 12 | N/A | <20 | 1,301 | 763 | 710 | <20 | <20 | <20 | <20 | <20 | <20 | <20 | <20 | <20 |
| | | 16 | N/A | <20 | 3,018 | 3,233 | 183 | <20 | <20 | <20 | <20 | <20 | <20 | <20 | <20 | <20 |
| | | 24 | N/A | <20 | 2,307 | 2,822 | 1,103 | 286 | <20 | 21 | 24 | <20 | <20 | <20 | <20 | <20 |
| | | 32 | N/A | <20 | 1,239 | 1,780 | 974 | 342 | 41 | <20 | <20 | <20 | <20 | <20 | <20 | <20 |
| | | 36 | N/A | <20 | 644 | 996 | 362 | 148 | 23 | <20 | <20 | <20 | <20 | <20 | <20 | <20 |
| | | 40 | N/A | <20 | 518 | 1,869 | 534 | 97 | 26 | <20 | 149 | <20 | <20 | <20 | <20 | <20 |
| | | 44 | N/A | <20 | 404 | 5,222 | 405 | 116 | 101 | <20 | 96 | <20 | <20 | <20 | <20 | <20 |
| | | 48 | N/A | <20 | 1,285 | 7,587 | 880 | 110 | 57 | 48 | 166 | 29 | <20 | <20 | <20 | 32 |
| | | 52 | N/A | <20 | 3,181 | 10,555 | 1,491 | 143 | 145 | 159 | 627 | 31 | <20 | 25 | <20 | 75 |
| | | 56 | N/A | <20 | 2,608 | 3,609 | 1,306 | 123 | 167 | 216 | 682 | 123 | 64 | 41 | 62 | 120 |
| | | 60 | N/A | <20 | 3,435 | 9,042 | 1,559 | 60 | 44 | 24 | 181 | 32 | <20 | <20 | 32 | 27 |
| | | 64 | N/A | <20 | 15,326 | 9,452 | 980 | 102 | 85 | 126 | 259 | 68 | <20 | <20 | 83 | 81 |
| | | 68 | N/A | <20 | 3,519 | 4,407 | 1,256 | 158 | 73 | 196 | 962 | 32 | 25 | <20 | 61 | 51 |
| | | 76 | N/A | <20 | 4,250 | 6,596 | 1,086 | 220 | 60 | 37 | 395 | 27 | <20 | <20 | <20 | <20 |
| | | 80 | N/A | <20 | 2,492 | 3,628 | 768 | 132 | 49 | 64 | 365 | 51 | <20 | <20 | <20 | <20 |
| | | 84 | N/A | <20 | 2,125 | 3,348 | 757 | 97 | 40 | 24 | 184 | <20 | <20 | <20 | <20 | <20 |
| | AH82 | 12 | N/A | <20 | 85 | <20 | <20 | <20 | <20 | <20 | <20 | <20 | <20 | <20 | <20 | <20 |
| | | 16 | N/A | <20 | 3,352 | 250 | 623 | 31 | <20 | <20 | <20 | <20 | <20 | <20 | <20 | <20 |
| | | 24 | N/A | <20 | 4,598 | 402 | 1,309 | 435 | <20 | 20 | 40 | <20 | <20 | <20 | <20 | <20 |
| | | 32 | N/A | <20 | 4,378 | 606 | 3,256 | 388 | 25 | 21 | 32 | <20 | <20 | <20 | <20 | <20 |
| | | 36 | N/A | <20 | 4,097 | 546 | 2,917 | 459 | <20 | <20 | <20 | <20 | <20 | <20 | <20 | <20 |
| | | 40 | N/A | <20 | 2,473 | 382 | 2,007 | 328 | 34 | 52 | 28 | <20 | <20 | <20 | <20 | <20 |
| | | 44 | N/A | <20 | 1,633 | 245 | 1,127 | 225 | 53 | 43 | 34 | <20 | <20 | <20 | <20 | <20 |
| | | 48 | N/A | <20 | 3,274 | 1,361 | 2,124 | 218 | 37 | 33 | <20 | <20 | <20 | <20 | <20 | <20 |
| | | 52 | N/A | <20 | 2,335 | 262 | 1,204 | 265 | 26 | <20 | 26 | <20 | <20 | <20 | <20 | <20 |
| | | 56 | N/A | <20 | 1,203 | 159 | 622 | 158 | <20 | <20 | <20 | <20 | <20 | <20 | <20 | <20 |
| | | 60 | N/A | <20 | 1,076 | 217 | 596 | 151 | <20 | <20 | <20 | <20 | <20 | <20 | <20 | <20 |
| | | 80 | N/A | <20 | 824 | 401 | 500 | 220 | <20 | <20 | <20 | 21 | <20 | <20 | <20 | <20 |
| | | 84 | N/A | <20 | 1,040 | 373 | 719 | 339 | <20 | <20 | <20 | <20 | <20 | <20 | <20 | <20 |
| | AG99 | 12 | N/A | <20 | 46 | 48 | 24 | <20 | <20 | <20 | <20 | <20 | <20 | <20 | <20 | <20 |
| | | 16 | N/A | <20 | 300 | 364 | 223 | <20 | <20 | <20 | <20 | <20 | <20 | <20 | <20 | <20 |
| | | 24 | N/A | <20 | 606 | 771 | 337 | <20 | <20 | <20 | <20 | <20 | <20 | <20 | <20 | <20 |
| | | 32 | N/A | <20 | 333 | 402 | 273 | <20 | <20 | <20 | <20 | <20 | <20 | <20 | <20 | <20 |
| | | 56 | N/A | <20 | 156 | 211 | 144 | <20 | <20 | <20 | <20 | <20 | <20 | <20 | <20 | <20 |
| | | 80 | N/A | <20 | 84 | 174 | 74 | <20 | <20 | <20 | <20 | 51 | <20 | <20 | <20 | <20 |
| | | 112 | N/A | <20 | 75 | 76 | 61 | <20 | <20 | <20 | <20 | <20 | <20 | <20 | <20 | <20 |
| | | 116 | N/A | <20 | 42 | 60 | 40 | <20 | <20 | <20 | <20 | <20 | <20 | <20 | <20 | <20 |

**Reciprocal $ID_{50}$**

- <20
- 20-100
- 100-1000
- 1000-10000
- >10000

**Fig. S2. SHIV.5MUT-infected rhesus macaques develop heterologous plasma neutralization breadth by week 20-32 post-infection. (A)** Longitudinal plasma neutralization against the indicated viruses, expressed as reciprocal $ID_{50}$.

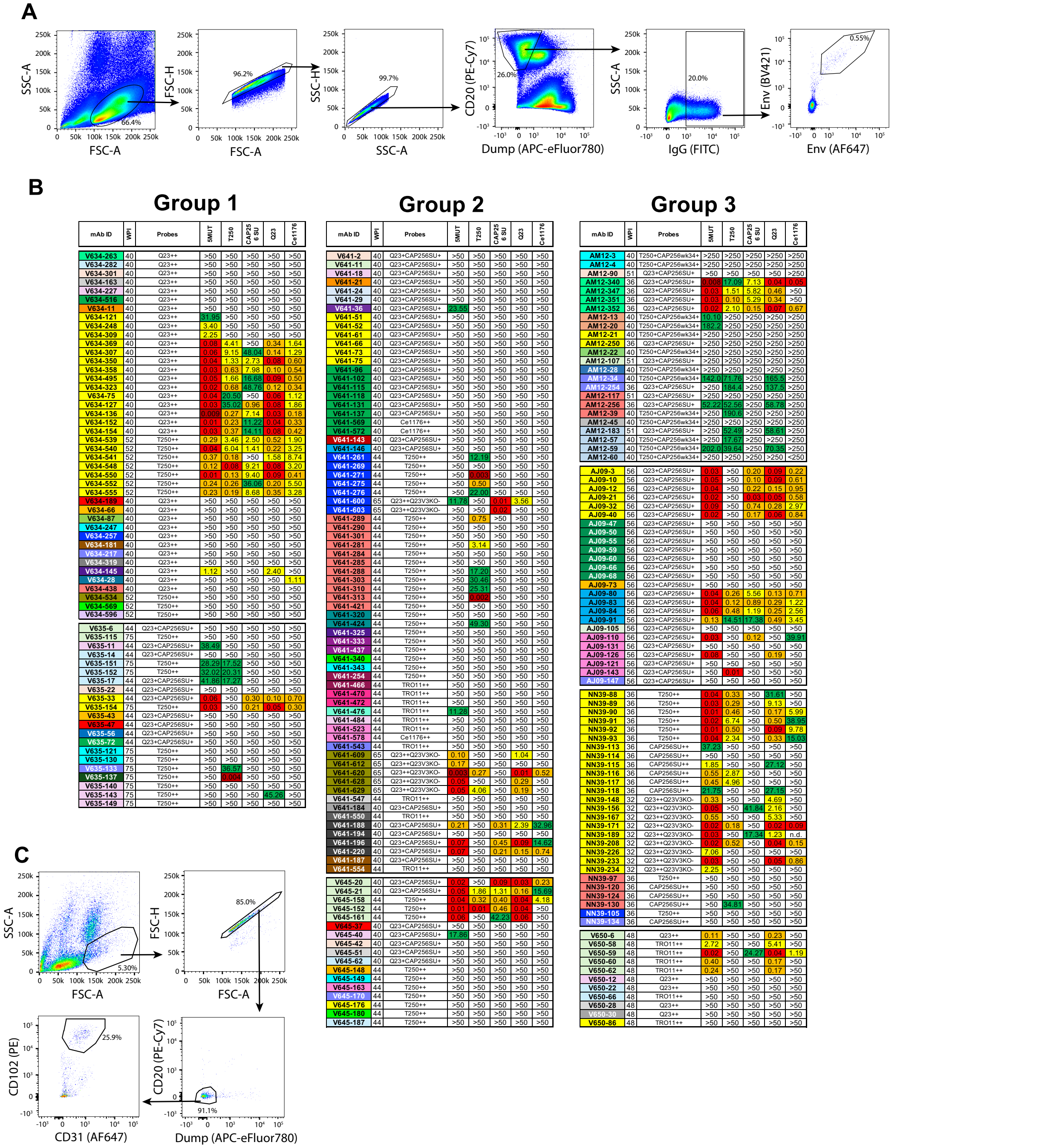


**Fig. S3. Isolation of candidate bNAbs from SHIV.5MUT-infected macaques. (A)** Representative gating strategy for the fluorescence-activated cell sorting used to isolate Env-binding B cells. Live $CD3^{-}CD8a^{-}CD14^{-}CD16^{-}CD20^{+}IgG^{+}Env^{++}$ B cells were sorted using Env trimer probes. **(B)** Monoclonal antibodies were synthesized and screened for neutralization activity against the autologous (SHIV.5MUT) and heterologous (T250, CAP256SU, Q23, and Ce1176) viruses. Titers are expressed as $IC_{50}$ in μg/mL. The timepoint and Env probes used to isolate each B cell are listed. Within each animal, antibody names are color-coded by lineage. **(C)** Gating strategy for the isolation of bone marrow plasma cells. WPI, weeks post-infection.

| | Clone Name | Subtype | V1 length (C-to-C) | Glycan 332 | 324-336 | AJ09-21 | AJ09-83 | AJ09-110 | AM12 lineage | | | | V635-33 | V645-158 | V650-59 | V634 lineage | | | NN39-171 | V641-220 | V641-600 | V641-620 |
|---|---|---|---|---|---|---|---|---|---|---|---|---|---|---|---|---|---|---|---|---|---|---|
| | | | | | | | | | AM12-347 | AM12-340 | AM12-352 | AM12-BM5 | | | | V634-136 | V634-540 | V634-555 | | | | |
| 5MUT | 5MUT | A | 17 | 332 | GDIRQAHCNVSKA | 0.030 | 0.040 | 0.030 | 0.033 | 0.008 | 0.015 | 0.009 | 0.061 | 0.035 | 0.018 | 0.009 | 0.044 | 0.232 | 0.016 | 0.065 | 11.780 | 0.003 |
| A1.KE.1993.BG505_W6M_C2.DQ208458 T332N | BG505.T332N | A | 17 | 332 | GDIRQAHCNVSKA | 4.580 | 0.270 | 2.030 | 2.010 | 0.039 | 0.100 | 0.077 | 0.092 | 1.030 | >50 | 0.191 | 1.064 | 1.400 | 0.102 | 0.384 | >50 | 0.504 |
| B.US.1995.6535_3.AY835438 | 6535.3 | B | 27 | 332 | GDIRQAHCNISRA | 0.019 | 0.084 | 0.013 | >50 | 0.105 | 0.084 | 0.027 | 0.020 | 0.011 | 0.110 | 0.182 | 0.031 | 0.159 | 0.014 | >50 | >50 | 0.300 |
| B.US.1995.6240_08_TA5_4622.EU289190 | 6240_08_TA5_4622 | B | 31 | 332 | GDIRQAHCNLSSK | 39.505 | 2.134 | 0.863 | >50 | 0.401 | 3.773 | 0.512 | 0.230 | 0.963 | 33.220 | 0.553 | 2.462 | >50 | 0.012 | >50 | >50 | >50 |
| B.IT.1995.TRO_11.AY835445 | TRO.11 | B | 30 | 332 | GDIRQAHCNISRT | 4.514 | 0.566 | >50 | 1.713 | 0.529 | 0.429 | 0.135 | 0.222 | 4.998 | >50 | 0.154 | 0.908 | 0.222 | 0.020 | >50 | >50 | >50 |
| B.US.1998.1056_10_TA11_1826.EU289186 | 1056_10_TA11_1826 | B | 25 | 332 | GDIRQAHCNISGE | >50 | 2.623 | >50 | 0.128 | 0.017 | 0.042 | 0.009 | 0.192 | 2.742 | >50 | 1.223 | 0.881 | >50 | >50 | >50 | >50 | 0.192 |
| B.IT.1996.PVO_4.AY835444 | PVO.4 | B | 29 | 332 | GDIRQAHCNLSRA | >50 | 9.601 | >50 | >50 | 3.956 | 3.817 | 0.200 | 0.218 | 2.917 | >50 | 6.444 | 8.122 | 1.751 | 3.972 | >50 | >50 | >50 |
| B.TT.1994.QH0692_42.AY835439 | QH0692.42 | B | 29 | 332 | GDIRQAHCNLSSV | >50 | 4.478 | >50 | >50 | 1.483 | 2.006 | 0.266 | 0.245 | >50 | >50 | 0.756 | 13.664 | 35.161 | >50 | >50 | >50 | >50 |
| B.US.1998.AC10_29.AY835446 | AC10.0.29 | B | 25 | 332 | GDIRQAHCNISRQ | >50 | 40.245 | >50 | >50 | 1.476 | 49.062 | 0.145 | 0.021 | >50 | >50 | >50 | 13.356 | >50 | 0.357 | >50 | >50 | 42.939 |
| B.US.2000.RHPA4259_7.AY835447 | RHPA4259.7 | B | 21 | 332 | GDIRQAHCNISRE | 0.061 | 0.220 | >50 | >50 | >50 | 7.693 | 0.023 | 0.057 | 0.463 | >50 | 0.128 | 0.467 | 1.494 | >50 | >50 | >50 | >50 |
| B.US.1990.WEAU_D15_410.EU289202 | WEAU_d15_410_787 | B | 30 | 332 | GDIRRAHCNLSRT | >50 | 13.319 | 0.545 | >50 | >50 | 0.616 | 0.090 | 0.178 | 1.320 | >50 | 8.348 | 3.269 | 15.185 | 0.076 | >50 | >50 | >50 |
| B.US.2004.CAAN5342.AY835452 | CAAN5342.A2 | B | 22 | 332 | GDIRKAHCNISGE | >50 | 49.129 | 1.580 | >50 | >50 | 0.146 | 0.010 | 0.615 | 0.176 | >50 | >50 | 0.838 | >50 | 3.771 | >50 | >50 | 7.248 |
| B.X2278_C2_B6.FJ817366 | X2278 | B | 27 | 332 | GDIRKAYCNISAT | >50 | >50 | 2.150 | >50 | >50 | 2.947 | 3.505 | >50 | 0.520 | 12.900 | >50 | >50 | 20.290 | 0.028 | >50 | >50 | >50 |
| B.US.1996.6244_13_B5_4567.EU289191 | 6244_13_B5_4576 | B | 37 | 332 | GDIRQAHCNISRA | >50 | 4.096 | >50 | >50 | >50 | 0.422 | 0.097 | >50 | 6.721 | >50 | 0.534 | 5.487 | 20.968 | 21.780 | >50 | >50 | >50 |
| B.US.2001.TRJO4551_58.AY835450 | TRJO4551.58 | B | 32 | 332 | GDIRKAYCNISEA | >50 | >50 | >50 | >50 | >50 | 0.655 | 0.076 | >50 | 1.359 | >50 | >50 | >50 | >50 | >50 | >50 | >50 | >50 |
| B.TT.1993.SC05_8C11_2344.EU289200 | SC05_8C11_2344 | B | 20 | 332 | GDIRQAHCNLSNT | 2.910 | 1.078 | 1.455 | 37.288 | >50 | >50 | 0.454 | 0.053 | 0.125 | 17.166 | 1.890 | 2.958 | 47.689 | 0.017 | 10.822 | >50 | >50 |
| B.US.1997.1012_11_TC21_3257.EU289184 | 1012_11_TC21_3257 | B | 28 | 332 | GNIRQAHCNISRS | >50 | 2.852 | >50 | >50 | >50 | >50 | >50 | >50 | >50 | >50 | 1.019 | 7.217 | >50 | >50 | >50 | >50 | >50 |
| B.TT.1995.SC422_8.AY835441 | SC422661.8 | B | 22 | 332 | GDIRQVHCNLSSA | 13.155 | >50 | >50 | >50 | >50 | >50 | >50 | 0.100 | >50 | >50 | 47.796 | >50 | >50 | >50 | >50 | >50 | >50 |
| B.US.2000.WITO4160_33.AY835451 | WITO4160.33 | B | 23 | 332 | GDIRKAHCNISTE | >50 | >50 | >50 | >50 | >50 | >50 | 2.341 | >50 | >50 | >50 | >50 | >50 | >50 | >50 | >50 | >50 | >50 |
| B.US.1997.1006_11_C3_1601.EU289183 | 1006_11_C3_1601 | B | 21 | 332 | GDIKRAYCNISRT | >50 | >50 | >50 | >50 | >50 | >50 | >50 | >50 | 0.044 | >50 | 1.626 | >50 | 0.486 | >50 | >50 | 0.012 | >50 |
| B.US.1997.1054_07_TC4_1499.EU289185 | 1054_07_TC4_1499 | B | 29 | 332 | GDIRKAYCNISRA | >50 | >50 | >50 | >50 | >50 | >50 | 0.187 | >50 | >50 | >50 | >50 | >50 | >50 | >50 | >50 | >50 | >50 |
| B.US.1997.62357_14_D3_4589.EU289189 | 62357_14_D3_4589 | B | 18 | 332 | GDIRQAYCNISRA | >50 | >50 | >50 | >50 | >50 | >50 | >50 | >50 | >50 | >50 | >50 | >50 | >50 | >50 | >50 | >50 | >50 |
| B.US.2001.REJO4541_67.AY835449 | REJO4541.67 | B | 27 | 334 | GDIRKAYCTINES | >50 | >50 | >50 | >50 | >50 | >50 | >50 | >50 | >50 | >50 | >50 | >50 | >50 | >50 | >50 | >50 | >50 |
| B.US.2000.THRO4156.AY835448 | THRO4156.18 | B | 33 | 334 | GDIRKAYCTVNGT | >50 | >50 | >50 | >50 | >50 | >50 | >50 | >50 | >50 | >50 | >50 | >50 | >50 | >50 | >50 | >50 | >50 |
| C.ZA.1999.DU156_12.DQ411852 | Du156.12 | C | 23 | 332 | GDIRQAHCNISRN | 0.108 | 0.352 | 6.054 | >50 | 0.099 | 3.078 | 0.280 | 0.187 | 0.913 | 1.477 | 0.246 | 0.649 | 8.116 | 0.017 | 3.328 | >50 | 4.623 |
| C.ZA.2005.CAP256_SU.KF241776 | CAP256.SU | C | 20 | 332 | GDIRQAHCNISEI | 0.020 | 1.270 | 0.050 | 5.819 | 7.529 | 0.146 | 0.046 | 0.300 | 0.399 | 24.270 | 7.137 | 1.406 | 8.680 | >50 | 0.205 | 0.011 | >50 |
| C.ZA.1998.DU422_1.DQ411854 | Du422.1 | C | 25 | 332 | GDIREAHCNISRE | 2.107 | 26.124 | >50 | 45.467 | 41.187 | 0.501 | 0.085 | 0.326 | 6.319 | 1.842 | 39.098 | >50 | >50 | 0.022 | >50 | >50 | 17.447 |
| C.MW.2004.CE1176_A3.FJ444437 | Ce1176_A3 | C | 21 | 332 | GDIRQAHCNVSKQ | 1.279 | 1.077 | >50 | >50 | 0.058 | 0.762 | 0.088 | 0.244 | 5.050 | 3.163 | 0.452 | 3.186 | 3.944 | 0.042 | 1.926 | >50 | 21.202 |
| C.IN.1999.25710_2_43.EF117271 | 25710 | C | 14 | 332 | GDIRQAHCNISKD | 33.780 | 0.820 | >50 | >50 | 0.095 | 0.111 | 0.052 | 14.390 | >50 | >50 | 0.094 | 0.387 | 0.488 | 6.398 | 0.140 | >50 | 7.876 |
| C.ZA.1998.DU172_17.DQ411853 | Du172.17 | C | 28 | 332 | GDIRQAHCNISRK | >50 | 15.663 | >50 | >50 | 0.023 | 2.788 | 0.501 | 17.283 | >50 | >50 | 2.825 | 45.341 | >50 | >50 | >50 | >50 | >50 |
| C.MW.2004.CE1172_H1.FJ444434 | Ce1172_H1 | C | 23 | 332 | GDIRQAHCNISRE | >50 | 0.742 | >50 | >50 | 0.038 | 2.268 | 0.123 | 0.106 | >50 | >50 | 0.239 | 1.718 | 2.822 | 0.158 | >50 | >50 | >50 |
| C.ZM.2003.ZM247_V1.FJ496204 | ZM247v1(Rev-) | C | 20 | 332 | GDIREAHCNITKH | >50 | 8.305 | >50 | >50 | 44.267 | 1.618 | 0.138 | 0.388 | >50 | >50 | 0.743 | >50 | >50 | >50 | >50 | >50 | >50 |
| C.MW.2004.1394_C9G1.FJ444527 | 1394C9G1(Rev-) | C | 32 | 332 | GDIRQAHCNISKA | >50 | 0.268 | >50 | 0.673 | 0.107 | 0.150 | 0.019 | 0.156 | >50 | >50 | 4.210 | >50 | >50 | 1.757 | 1.266 | >50 | 0.477 |
| C.ZA.2007.704809221_1B3.FJ444103 | Ce704809221_1B3 | C | 34 | 332 | GDIRQAHCNISEE | >50 | 9.764 | >50 | >50 | 0.122 | 0.844 | 0.086 | 2.121 | >50 | >50 | 28.757 | >50 | >50 | 7.738 | >50 | >50 | 0.717 |
| C.ZM.2003.ZM214_15.DQ388516 | ZM214M.PL15 | C | 26 | 332 | GDIRQAHCNISKD | >50 | >50 | >50 | 48.187 | 0.677 | 1.245 | 0.118 | 2.406 | >50 | >50 | >50 | >50 | >50 | >50 | 0.312 | >50 | >50 |
| C.CN.2007.CNE17.HM215403 | CNE17 | C | 44 | 332 | GDIRQAHCNISKG | >50 | 33.732 | >50 | >50 | 0.633 | 5.629 | 0.596 | >50 | >50 | >50 | 11.929 | >50 | >50 | >50 | 30.875 | >50 | >50 |
| C.CH1012 | CH1012 | C | 27 | 332 | GDIRRAYCNISAE | >50 | >50 | >50 | >50 | 23.490 | 0.610 | 0.314 | >50 | >50 | >50 | >50 | >50 | >50 | >50 | >50 | 1.828 | >50 |
| C.MW.2003.CE0682_E4.FJ444325 | Ce0682_E4 | C | 27 | 332 | GDIRQAHCNISDE | >50 | >50 | >50 | 0.048 | >50 | 0.020 | 0.006 | >50 | >50 | >50 | >50 | >50 | >50 | >50 | >50 | >50 | >50 |
| C.CE703010217_B6.FJ443575 | Ce0217 | C | 15 | 332 | GDIRRAYCNISEK | >50 | >50 | >50 | >50 | >50 | 8.332 | 0.264 | >50 | 27.550 | 12.390 | >50 | >50 | 17.020 | 0.026 | >50 | >50 | >50 |
| C.IN.2000.001428_2_42.EF117266 | HIV-001428-2.42 | C | 25 | 332 | GNIREAHCNISEK | >50 | 0.588 | >50 | >50 | 12.382 | >50 | 0.279 | >50 | >50 | >50 | 11.423 | >50 | >50 | >50 | >50 | >50 | >50 |
| C.IN.1999.16845_2_22.EF117269 | HIV-16845-2.22 | C | 31 | 332 | GDIRQAHCNISEA | >50 | 26.393 | >50 | >50 | 22.610 | >50 | 29.934 | >50 | >50 | >50 | 27.530 | >50 | >50 | >50 | >50 | >50 | >50 |
| C.ZM.2003.ZM246F.FJ496194 | 246F C1G | C | 12 | 332 | GDIREAHCNISST | >50 | >50 | >50 | >50 | 0.029 | >50 | >50 | >50 | >50 | >50 | >50 | 25.824 | >50 | >50 | >50 | >50 | >50 |
| C.CH848_3_d0017_10_02.KX216883 | CH848 | C | 32 | 332 | GDIRQAHCNISER | >50 | >50 | >50 | >50 | 16.220 | >50 | 36.590 | >50 | >50 | >50 | >50 | >50 | >50 | >50 | >50 | >50 | >50 |
| C.MW.2007.703010200_1E5.FJ443999 | 7030102001E5(Rev-) | C | 21 | 332 | GDIRQAHCNISKK | 3.196 | 0.058 | 0.301 | >50 | >50 | >50 | 0.097 | 0.007 | 0.048 | >50 | 0.039 | 0.029 | 0.126 | 0.413 | 32.201 | 0.014 | 0.357 |
| C.TZ.2003.3301_V1_C24.HM215294 | 3301.v1.c24 | C | 19 | 332 | GDIRQAHCNITRN | >50 | 0.975 | >50 | >50 | >50 | 3.946 | 0.347 | 0.209 | >50 | >50 | 18.696 | 0.822 | 0.932 | 0.543 | >50 | >50 | >50 |
| C.ZM.2002.ZM233_6.DQ388517 | ZM233M.PB6 | C | 15 | 332 | GNIREAHCNISAS | >50 | 0.168 | >50 | >50 | >50 | >50 | >50 | >50 | >50 | >50 | 0.654 | >50 | 1.142 | >50 | 42.564 | >50 | >50 |
| C.ZM.1998.ZM135_10A.AY424079 | ZM135M.PL10a | C | 21 | 332 | GDIRQAHCNISGE | >50 | 47.228 | >50 | >50 | >50 | >50 | 41.736 | 25.306 | >50 | >50 | >50 | >50 | >50 | 0.810 | >50 | >50 | >50 |
| C.ZA.2005.CAP210_E8.DQ435683 | CAP210.2.00.E8 | C | 17 | 332 | GNIREAHCNISEK | >50 | 46.928 | >50 | >50 | >50 | >50 | >50 | >50 | >50 | >50 | >50 | >50 | >50 | >50 | >50 | >50 | >50 |
| C.CN.2006.CNE58.HM215421 | CNE58 | C | 37 | 332 | GDIRQAHCNITKS | >50 | >50 | >50 | >50 | >50 | >50 | >50 | >50 | >50 | >50 | >50 | >50 | >50 | >50 | >50 | >50 | >50 |
| C.ZM.2003.ZM249_1.DQ388514 | ZM249M.PL1 | C | 21 | 332 | GKIREAHCNISKE | >50 | >50 | >50 | >50 | >50 | >50 | >50 | >50 | >50 | >50 | >50 | >50 | >50 | >50 | >50 | >50 | >50 |
| C.MW.2005.CE2060_G9.FJ444607 | Ce2060_G9 | C | 32 | 332 | GNIRQAYCNITKG | >50 | >50 | >50 | >50 | >50 | >50 | >50 | >50 | >50 | >50 | >50 | >50 | >50 | >50 | >50 | >50 | >50 |
| C.IN.2000.0013095_2_11.EF117267 | HIV-0013095-2.11 | C | 22 | 332 | GDIRQARCNISEE | >50 | >50 | >50 | >50 | >50 | >50 | >50 | >50 | >50 | >50 | >50 | >50 | >50 | >50 | >50 | >50 | >50 |
| C.ZM.2003.249M_B10.EU166862 | 249M B10 | C | 21 | 332 | GKIREAHCNISKE | >50 | >50 | >50 | >50 | >50 | >50 | >50 | >50 | >50 | >50 | >50 | >50 | >50 | >50 | >50 | >50 | >50 |
| C.MW.2002.BF1266_431A.HM215360 | BF1266.431a | C | 24 | 334 | GDTRQAYCTINGT | >50 | >50 | >50 | >50 | >50 | 0.631 | 0.389 | >50 | >50 | >50 | >50 | >50 | >50 | >50 | >50 | >50 | >50 |
| C.MW.2008.CH505_TF.KC247556 | CH505 | C | 16 | 334 | GDIREAYCNINES | >50 | >50 | >50 | >50 | >50 | 2.106 | 1.428 | >50 | >50 | >50 | >50 | >50 | >50 | >50 | >50 | >50 | >50 |
| C.ZM.2000.ZM109_4.AY424138 | ZM109F.PB4 | C | 15 | 334 | GDIRKAYCKINGS | >50 | >50 | >50 | >50 | >50 | >50 | >50 | >50 | >50 | >50 | >50 | >50 | >50 | >50 | >50 | >50 | >50 |
| C.ZA.2005.CAP45_G3.DQ435682 | CAP45.2.00.G3 | C | 15 | 334 | GDIRQAHCNINNS | >50 | >50 | >50 | >50 | >50 | >50 | >50 | >50 | >50 | >50 | >50 | >50 | >50 | >50 | >50 | >50 | >50 |
| C.MW.2004.CE1086_B2.FJ444395 | Ce1086_B2 | C | 16 | 334 | GNIRQAHCNINES | >50 | >50 | >50 | >50 | >50 | >50 | >50 | >50 | >50 | >50 | >50 | >50 | >50 | >50 | >50 | >50 | >50 |
| C.MW.2005.CE2010_F5.FJ444561 | Ce2010_F5 | C | 31 | 334 | GNIRQAHCTINAT | >50 | >50 | >50 | >50 | >50 | >50 | >50 | >50 | >50 | >50 | >50 | >50 | >50 | >50 | >50 | >50 | >50 |
| C.MW.2007.703010054_2A2.FJ443808 | Ce703010054_2A2 | C | 15 | 334 | GDIRQAHCDINGT | >50 | >50 | >50 | >50 | >50 | >50 | >50 | >50 | >50 | >50 | >50 | >50 | >50 | >50 | >50 | >50 | >50 |
| C.ZM.2000.ZM53_12.AY423984 | ZM53M.PB12 | C | 13 | none | GDIRQAYCIINKA | >50 | >50 | >50 | >50 | >50 | 0.818 | 0.045 | >50 | >50 | >50 | >50 | >50 | >50 | >50 | >50 | >50 | >50 |
| C.ZM.2002.ZM197_7.DQ388515 | ZM197M.PB7 | C | 37 | none | GDIRQAHCDLSKS | >50 | >50 | >50 | >50 | >50 | >50 | >50 | >50 | >50 | >50 | >50 | >50 | >50 | >50 | >50 | >50 | >50 |
| C.IN.1999.16055_2_3.EF117268 | HIV-16055-2.3 | C | 22 | none | GNIRQAYCNIKKD | >50 | >50 | >50 | >50 | >50 | >50 | >50 | >50 | >50 | >50 | >50 | >50 | >50 | >50 | >50 | >50 | >50 |
| C.MW.2003.CE0393_C3.FJ444215 | Ce0393_C3 | C | 32 | none | GDIRQASCNIDEK | >50 | >50 | >50 | >50 | >50 | >50 | >50 | >50 | >50 | >50 | >50 | >50 | >50 | >50 | >50 | >50 | >50 |
| 07_BC.CN.2004.CH119_10.EF117261 | CH119 | BC | 19 | 332 | GDIRQAHCNISKW | 0.080 | 0.240 | >50 | >50 | 3.325 | 2.971 | 0.422 | 0.037 | 45.080 | 7.068 | 0.062 | 0.437 | 0.947 | 0.029 | 12.000 | >50 | 17.320 |
| 07_BC.CN.2007.BJOX002000_03_2.HM215364 | BJOX002000 | BC | 20 | 332 | GDIRQAHCNISGK | 5.510 | 1.210 | 0.450 | >50 | 14.580 | 0.484 | 0.065 | 0.012 | 0.940 | 12.760 | 0.040 | 2.436 | 7.846 | 0.018 | >50 | >50 | 38.880 |
| 07_BC.CN.2007.CNE21.HM215407 | CNE21 | BC | 24 | 332 | GDIRQAHCNISET | >50 | 0.305 | >50 | >50 | 0.024 | 0.666 | 0.028 | 13.382 | >50 | >50 | 0.037 | 0.411 | 0.402 | 1.036 | 30.481 | >50 | 9.944 |
| 07_BC.CN.2007.CNE20.HM215406 | CNE20 | BC | 15 | 332 | GNIRQAHCNISEN | 31.652 | 0.027 | >50 | >50 | >50 | 45.624 | 0.011 | >50 | >50 | >50 | 0.007 | 0.111 | 0.032 | >50 | 0.027 | >50 | 1.958 |
| 08_BC.CN.2007.CNE53.HM215417 | CNE53 | BC | 18 | 332 | GDIRQAHCNISES | 0.291 | 0.558 | >50 | >50 | 0.585 | >50 | 0.819 | 0.319 | 15.932 | 19.139 | 0.153 | 0.549 | 0.326 | 0.037 | 2.035 | >50 | >50 |
| 08_BC.CN.2007.CNE30.HM215411 | CNE30 | BC | 23 | 332 | GDIRQAHCNISRN | >50 | 21.972 | >50 | >50 | >50 | >50 | >50 | 4.944 | >50 | >50 | 4.875 | 38.840 | >50 | 1.582 | >50 | >50 | >50 |
| 08_BC.CN.2007.CNE52.HM215416 | CNE52 | BC | 23 | 332 | GDIRQAHCNISKE | >50 | >50 | >50 | >50 | >50 | >50 | >50 | >50 | >50 | >50 | >50 | >50 | >50 | >50 | >50 | >50 | >50 |
| 07_BC.CN.2007.CNE19.HM215405 | CNE19 | BC | 20 | 332 | GDIRQAYCNISKD | >50 | >50 | >50 | >50 | >50 | >50 | 1.872 | >50 | >50 | >50 | >50 | >50 | 1.432 | >50 | >50 | 0.022 | >50 |
| A1.KE.1994.Q23_17.AF004885 | Q23.17 | A | 17 | 332 | GDIRQAHCNVTRS | 0.050 | 0.234 | 0.006 | 0.193 | 0.018 | 0.077 | 0.010 | 0.019 | 0.011 | 0.016 | 0.061 | 0.130 | 0.158 | 0.003 | 0.057 | 0.222 | 0.075 |
| A1.UG.2007.9004SS_A3_4.HM215350 | 9004SS_A3_4 | A | 31 | 332 | GNIRQAHCNVSKA | >50 | 0.204 | >50 | 38.799 | 8.051 | 1.944 | 0.018 | 25.787 | >50 | >50 | >50 | 10.368 | 25.113 | >50 | >50 | >50 | >50 |
| A1.TZ.2005.0260_V5_C36.HM215256 | 0260.v5.c36 | A | 32 | 332 | GDIRQAHCNVSKA | >50 | 20.377 | 9.558 | >50 | >50 | 3.821 | 0.778 | >50 | 8.242 | >50 | 3.181 | 15.435 | >50 | 0.107 | >50 | >50 | >50 |
| A1.TZ.2004.3365_V2_C20.HM215298 | 3365.v2.c2 | A | 20 | 332 | EDIREAHCNVSQT | >50 | 2.149 | >50 | >50 | >50 | 1.168 | 0.028 | 0.535 | >50 | >50 | >50 | 12.049 | >50 | 0.061 | 3.622 | >50 | >50 |
| A1D.KE..MS208_A1.DQ187010 | MS208.A1 | A | 21 | 332 | GDIRQAHCNVSGS | >50 | >50 | >50 | >50 | >50 | 0.148 | 0.031 | >50 | >50 | >50 | >50 | >50 | >50 | >50 | >50 | >50 | >50 |
| A1.UG.2007.191084_B7_19.HM215266 | 191084 B7-19 | A | 25 | 332 | GDIRQAHCNVSES | >50 | 12.385 | 1.407 | >50 | >50 | >50 | 3.840 | 0.029 | 0.096 | 2.853 | >50 | 23.794 | >50 | 0.015 | 1.803 | >50 | >50 |
| A1.TZ.2003.3415_V1_C1.HM215299 | 3415.v1.c1 | A | 22 | 332 | GGIRQAHCNVSRK | >50 | 1.576 | >50 | >50 | >50 | >50 | >50 | >50 | >50 | >50 | 0.623 | 20.837 | 21.036 | >50 | >50 | >50 | >50 |
| A1.KE.1994.Q842_D12.AF407160 | Q842.d12 | A | 17 | 334 | GDIRQAHCNVNRT | >50 | >50 | >50 | >50 | 1.404 | 3.652 | 0.919 | >50 | >50 | >50 | >50 | >50 | >50 | >50 | >50 | >50 | >50 |
| A1.KE.1994.Q259_17.AF407152 | Q259.d2.17 | A | 19 | 334 | GNIRQAYCTVNRT | >50 | >50 | >50 | >50 | >50 | 1.932 | 0.021 | >50 | >50 | >50 | >50 | >50 | >50 | >50 | >50 | >50 | >50 |
| A1.UG.2007.191955_A11.HM215272 | 191955_A11 | A | 21 | 334 | GDIRQAYCKVNKS | >50 | >50 | >50 | >50 | >50 | >50 | >50 | >50 | >50 | >50 | >50 | >50 | >50 | >50 | >50 | >50 | >50 |
| A1.KE.1995.Q461_E2.AF407156 | Q461.e2 | A | 25 | 334 | GDIRNAHCVVNRT | >50 | >50 | >50 | >50 | >50 | >50 | 8.823 | >50 | >50 | >50 | >50 | >50 | >50 | >50 | >50 | >50 | >50 |
| A1.KE.1996.Q769_D22.AF407158 | Q769.d22 | A | 22 | none | GDIRKAHCNVNRQ | >50 | >50 | >50 | >50 | >50 | 2.710 | >50 | >50 | >50 | >50 | >50 | >50 | >50 | >50 | >50 | >50 | >50 |
| 02_AG.CI.1999.928_28.EU513199 | 928-28 | AG | 23 | 332 | GDIRQAHCNVSKI | >50 | >50 | >50 | >50 | >50 | >50 | >50 | >50 | >50 | >50 | >50 | >50 | >50 | >50 | >50 | >50 | >50 |
| 02_AG.CM.2004.263_8.EU513182 | 263-8 | AG | 27 | 332 | GDIRQAYCNVSKG | >50 | >50 | >50 | >50 | >50 | >50 | >50 | >50 | 0.642 | >50 | >50 | >50 | 0.652 | >50 | >50 | >50 | >50 |
| 02_AG.CM.2004.T250_4.EU513189 | T250-4 | AG | 20 | 332 | GNIRKAYCNVSKR | >50 | 0.301 | >50 | 13.903 | 3.330 | 22.602 | 0.012 | >50 | 0.525 | >50 | 0.813 | 22.163 | 0.173 | >50 | >50 | 0.004 | >50 |
| 02_AG.CM.2004.T251_18.EU513196 | T251-18 | AG | 22 | 332 | GDIRQAHCNVSRG | >50 | 4.881 | >50 | >50 | >50 | >50 | 0.317 | >50 | >50 | >50 | >50 | >50 | >50 | >50 | >50 | >50 | >50 |
| 02_AG.CM.2005.T278_50.EU513198 | T278-50 | AG | 24 | 332 | GDIRQAYCNVSGT | >50 | >50 | >50 | >50 | >50 | >50 | 1.267 | >50 | 1.188 | >50 | >50 | >50 | >50 | >50 | >50 | >50 | >50 |
| 02_AG.CM.2004.T255_34.EU513184 | T255-34 | AG | 26 | 332 | GDIRQAHCNVSK | >50 | 0.162 | >50 | >50 | >50 | 0.061 | 0.008 | >50 | >50 | >50 | >50 | 31.629 | 0.280 | >50 | >50 | >50 | >50 |
| 02_AG.CM.2004.211_9.EU513187 | 211-9 | AG | 32 | 332 | GNIRQAHCNVSRA | >50 | >50 | >50 | >50 | >50 | >50 | >50 | >50 | >50 | >50 | >50 | >50 | >50 | >50 | >50 | >50 | >50 |
| 02_AG.CM.2004.235_47.EU513195 | 235-47 | AG | 20 | 332 | GDIRQAHCNVSST | >50 | >50 | >50 | >50 | >50 | >50 | >50 | >50 | >50 | 6.484 | >50 | >50 | >50 | 5.762 | >50 | >50 | >50 |
| 02A1.CM.2004.T257_31.EU513185 | T257-31 | AG | 31 | 334 | GDIRQAHCKVNKT | >50 | >50 | >50 | >50 | 40.754 | 10.319 | 6.449 | >50 | >50 | >50 | >50 | >50 | >50 | >50 | >50 | >50 | >50 |
| 01_AE.CN.2007.BJOX028000_10_3.HM215389 | BJOX028000.10.3 | AE | 29 | 332 | GDIRQAYCNISRT | >50 | >50 | >50 | >50 | >50 | >50 | 0.960 | >50 | >50 | >50 | >50 | >50 | >50 | >50 | >50 | >50 | >50 |
| 01_AE.TH.1999.C1080_C3.JN944660 | C1080.c03 | AE | 26 | 334 | GDIRKAYCEINGT | >50 | >50 | >50 | >50 | >50 | 5.819 | 0.798 | >50 | >50 | >50 | >50 | >50 | >50 | >50 | >50 | >50 | >50 |
| 01_AE.TH.1999.C2101_C1.JN944661 | C2101.c01 | AE | 26 | 334 | GDIRAVYCEINET | >50 | >50 | >50 | >50 | >50 | 1.271 | 0.250 | >50 | >50 | >50 | >50 | >50 | >50 | >50 | >50 | >50 | >50 |
| 01_AE.CN.2006.CNE5.HM215415 | CNE5 | AE | 28 | 334 | GDIRKAYCEINGT | >50 | >50 | >50 | >50 | >50 | 8.675 | 4.108 | >50 | >50 | >50 | >50 | >50 | >50 | >50 | >50 | >50 | >50 |
| 01_AE.CN.2006.CNE8.HM215427 | CNE8 | AE | 24 | 334 | GDIRKAYCEINRT | >50 | >50 | >50 | >50 | >50 | 0.975 | 0.231 | >50 | >50 | >50 | >50 | >50 | >50 | >50 | >50 | >50 | >50 |
| 01_AE.CN.2007.BJOX009000_02_4.HM215372 | BJOX009000.02.4 | AE | 25 | 334 | GDIRKAYCEINGT | >50 | >50 | >50 | >50 | >50 | 0.640 | 0.389 | >50 | >50 | >50 | >50 | >50 | >50 | >50 | >50 | >50 | >50 |
| 01_AE.CN.2007.BJOX010000_06_2.HM215373 | BJOX010000.06.2 | AE | 32 | 334 | GDIRKAYCKINAT | >50 | >50 | >50 | >50 | >50 | 0.304 | 0.216 | >50 | >50 | >50 | >50 | >50 | >50 | >50 | >50 | >50 | >50 |
| 01_AE.CN.2007.BJOX015000_11_5.HM215377 | BJOX015000.11.5 | AE | 29 | 334 | GDIRKAYCEINGT | >50 | >50 | >50 | >50 | >50 | 0.318 | 0.135 | >50 | >50 | >50 | >50 | >50 | >50 | >50 | >50 | >50 | >50 |
| 01_AE.CN.2007.BJOX025000_01_1.HM215386 | BJOX025000.01.1 | AE | 26 | 334 | GDIRQAYCEINGT | >50 | >50 | >50 | >50 | >50 | 29.184 | 3.422 | >50 | >50 | >50 | >50 | >50 | >50 | >50 | >50 | >50 | >50 |
| 01_AE.TH.1999.C3347_C11.JX512902 | C3347.c11 | AE | 30 | 334 | GNIRKAYCEINGT | >50 | >50 | >50 | >50 | >50 | >50 | 0.166 | >50 | >50 | >50 | >50 | >50 | >50 | >50 | >50 | >50 | >50 |
| 01_AE.TH.1999.R3265_C6.JX512903 | R3265.c06 | AE | 27 | 334 | GDIRQAYCEINQT | >50 | >50 | >50 | >50 | >50 | >50 | 7.964 | >50 | >50 | >50 | >50 | >50 | >50 | >50 | >50 | >50 | >50 |
| 01_AE.TH.2001.R2184_C4.JN944665 | R2184.c04 | AE | 28 | 334 | GNIRKAYCVINGT | >50 | >50 | >50 | >50 | >50 | >50 | >50 | >50 | >50 | >50 | >50 | >50 | >50 | >50 | >50 | >50 | >50 |
| 01_AE.TH.2005.620345_C1.JX512900 | 620345.c01 | AE | 27 | 334 | TSIRKAYCEINGT | >50 | >50 | >50 | >50 | >50 | >50 | >50 | >50 | >50 | >50 | >50 | >50 | >50 | >50 | >50 | >50 | >50 |
| 01_AE.TH.1998.R1166_C1.JQ352783 | R1166.c01 | AE | 39 | 334 | GDIRRAYCEINRT | >50 | >50 | >50 | >50 | >50 | >50 | >50 | >50 | >50 | >50 | >50 | >50 | >50 | >50 | >50 | >50 | >50 |
| 01_AE.CN.2007.CNE55.HM215418 | CNE55 | AE | 23 | none | GDIRKAYCEIDGT | >50 | >50 | >50 | >50 | >50 | 7.196 | 2.139 | >50 | >50 | >50 | >50 | >50 | >50 | >50 | >50 | >50 | >50 |
| 01_AE.TH.2000.C4118_9.JQ352782 | C4118.c09 | AE | 27 | none | GDIRNAYCEIDGT | >50 | >50 | >50 | >50 | >50 | >50 | >50 | >50 | >50 | >50 | >50 | >50 | >50 | >50 | >50 | >50 | >50 |
| G.ES.2008.P1981_C5_3.FJ817369 | P1981_C5_3 | G | 19 | 332 | GDIRQAHCNISKS | 0.084 | 0.147 | 0.014 | >50 | 0.069 | 0.836 | 0.034 | 0.033 | 0.080 | >50 | 2.832 | 0.118 | 0.055 | 0.039 | 0.021 | >50 | 0.072 |
| G.ES.2003.X1254_C3.EU885762 | X1254_c3 | G | 24 | 332 | GDIRQAHCNISGT | >50 | 2.169 | >50 | >50 | 0.084 | 1.319 | 0.122 | 4.303 | >50 | >50 | >50 | 1.875 | 1.853 | 0.286 | 46.377 | >50 | 0.336 |
| G.ES.2002.X1193_C1.EU885761 | X1193_c1 | G | 28 | 332 | GDIRQAHCNISKT | >50 | 2.436 | >50 | >50 | 0.109 | 0.506 | 0.080 | 27.541 | >50 | >50 | 0.649 | 10.408 | >50 | >50 | 0.236 | >50 | 1.868 |
| G.ES.2005.X2131_C1_B5.FJ817368 | X2131_C1_B5 | G | 28 | 332 | GDIRQAHCNISKT | >50 | 1.099 | >50 | >50 | 0.251 | 0.847 | 0.207 | 1.907 | >50 | >50 | 0.272 | 0.655 | 0.765 | 0.209 | 0.090 | >50 | 28.042 |
| G.ES.2002.P0402_C2_11.EU885759 | P0402_c2_11 | G | 22 | 332 | GNIRQAHCNISGA | >50 | 0.365 | >50 | >50 | 4.191 | >50 | 0.059 | 47.673 | >50 | >50 | >50 | 39.917 | >50 | >50 | >50 | >50 | >50 |
| G.ES.2006.X2088_C9.EU885764 | X2088_c9 | G | 13 | 332 | GDIRQAHCNVTRG | >50 | 9.522 | 4.193 | >50 | >50 | >50 | >50 | 0.065 | >50 | 0.018 | >50 | >50 | >50 | 0.002 | >50 | >50 | >50 |
| G.ES.2004.X1632_S2_B10.FJ817370 | X1632_S2_B10 | G | 25 | 334 | GDTRQAHCNINGS | >50 | >50 | >50 | >50 | >50 | >50 | >50 | >50 | >50 | >50 | >50 | >50 | >50 | >50 | >50 | >50 | >50 |
| D.UG.1999.A07412M1_VRC12.HM215357 | A07412M1.vrc12 | D | 23 | 332 | GNIRQAHCNVSKV | >50 | 21.609 | >50 | >50 | >50 | >50 | 16.774 | >50 | >50 | >50 | 10.641 | >50 | >50 | >50 | >50 | >50 | >50 |
| D.TZ.2004.6405_V4_C34.HM215327 | 6405.v4.c34 | D | 28 | 332 | GDIRQAHCNVSRV | >50 | 1.888 | >50 | >50 | >50 | >50 | 0.202 | 3.092 | >50 | >50 | 1.760 | 3.041 | 8.007 | >50 | >50 | >50 | >50 |
| D.UG.2001.231965_C1.JQ361079 | 231965.c01 | D | 34 | 332 | GNIRLAYCNISRK | >50 | >50 | >50 | >50 | >50 | >50 | >50 | >50 | >50 | >50 | 14.227 | >50 | >50 | >50 | >50 | 1.597 | >50 |
| D.UG.2001.231966_C2.JX512899 | 231966.c02 | D | 30 | 332 | GDIRQAYCNISET | >50 | >50 | >50 | >50 | >50 | >50 | 2.463 | >50 | >50 | >50 | >50 | >50 | >50 | >50 | >50 | >50 | >50 |
| D.TZ.2005.3016_V5_C45.HM215283 | 3016.v5.c45 | D | 21 | none | GDIRQAYCTIDGT | >50 | >50 | >50 | >50 | >50 | >50 | >50 | >50 | >50 | >50 | >50 | >50 | >50 | >50 | >50 | >50 | >50 |
| CD.TZ.2006.6811_V7_C18.HM215340 | 6811.v7.c18 | CD | 11 | 332 | GDIRQAHCNISES | 0.153 | 0.275 | 0.042 | >50 | 0.147 | 0.130 | 0.053 | 0.012 | 0.486 | 0.008 | 0.444 | 0.263 | 11.736 | 0.002 | 0.558 | >50 | 13.046 |
| CD.TZ.2004.6480_V4_C25.HM215329 | 6480.v4.c25 | CD | 19 | 332 | GDIRKAHCNISKD | >50 | 10.315 | 0.182 | >50 | 4.024 | 0.068 | 0.020 | 0.050 | 0.129 | 0.708 | 5.593 | 0.786 | 1.453 | 0.008 | >50 | >50 | >50 |
| CD.TZ.2004.3817_V2_C59.HM215310 | 3817.v2.c59 | CD | 21 | 332 | GDIRQAHCNISRG | >50 | >50 | >50 | >50 | 43.877 | 8.067 | 0.151 | 9.703 | >50 | >50 | >50 | >50 | >50 | 23.449 | >50 | >50 | >50 |
| 41_CD.TZ.2003.6952_V1_C20.HM215343 | 6952.v1.c20 | CD | 28 | 332 | GDIRQAHCNISGT | >50 | 0.648 | >50 | >50 | >50 | >50 | >50 | 11.070 | >50 | >50 | 0.080 | 5.668 | 0.792 | >50 | 2.668 | >50 | >50 |
| CD.TZ..89_F1_2_25.HM215349 | 89-F1_2_25 | CD | 14 | none | GDIRKAYCTISKG | >50 | >50 | >50 | >50 | >50 | >50 | >50 | >50 | >50 | >50 | >50 | >50 | >50 | >50 | >50 | >50 | >50 |
| A1CD.TZ.2004.3103_V3_C10.HM215288 | 3103.v3.c10 | A1CD | 23 | 332 | GDIRQAHCNISEA | 44.839 | 0.392 | >50 | 0.626 | 0.539 | 1.106 | 0.184 | 0.158 | 4.763 | 29.943 | 0.080 | 0.311 | 0.313 | 0.054 | >50 | >50 | 14.604 |
| A1D.TZ.2004.0815_V3_C3.HM215260 | 0815.v3.c3 | A1D | 26 | 332 | GDIRQAHCNVSRA | >50 | >50 | 2.838 | >50 | >50 | 3.735 | 0.172 | 0.806 | >50 | 1.163 | >50 | 7.882 | >50 | 0.089 | >50 | >50 | >50 |
| A1C.TZ.2004.6041_V3_C23.HM215321 | 6041.v3.c23 | A1C | 26 | 334 | GNIRQAHCDINRT | >50 | >50 | >50 | >50 | >50 | >50 | >50 | >50 | >50 | >50 | >50 | >50 | >50 | >50 | >50 | >50 | >50 |
| A1C.TZ.2004.6540_V4_C1.HM215330 | 6540.v4.c1 | A1C | 23 | 334 | GNIRQAYCTVNRT | >50 | >50 | >50 | >50 | >50 | >50 | 1.174 | >50 | >50 | >50 | >50 | >50 | >50 | >50 | >50 | >50 | >50 |
| A1C.TZ.2005.6545_V4_C1.HM215332 | 6545.v4.c1 | A1C | 23 | 334 | GNIRQAYCTVNRT | >50 | >50 | >50 | >50 | >50 | >50 | 14.385 | >50 | >50 | >50 | >50 | >50 | >50 | >50 | >50 | >50 | >50 |
| A1C.TZ.246_F3_C10_2.HM215279 | 246F3 | A1C | 20 | 334 | GNIRQAHCTVNKT | >50 | >50 | >50 | >50 | >50 | >50 | 7.004 | >50 | >50 | >50 | >50 | >50 | >50 | >50 | >50 | >50 | >50 |
| MuLV | | | | | | >50 | >50 | >50 | >50 | >50 | >50 | >50 | >50 | >50 | >50 | >50 | >50 | >50 | >50 | >50 | >50 | >50 |

| | AJ09-21 | AJ09-83 | AJ09-110 | AM12-347 | AM12-340 | AM12-352 | AM12-BM5 | V635-33 | V645-158 | V650-59 | V634-136 | V634-540 | V634-555 | NN39-171 | V641-220 | V641-600 | V641-620 |
|---|---|---|---|---|---|---|---|---|---|---|---|---|---|---|---|---|---|
| Whole Panel | 15.4% | 44.6% | 13.8% | 9.2% | 32.3% | 50.0% | 67.7% | 36.9% | 24.6% | 15.4% | 37.7% | 36.2% | 30.0% | 30.8% | 17.7% | 6.2% | 16.9% |
| GMT IC50 (μg/mL) | 0.97 | 1.80 | 0.41 | 2.79 | 0.79 | 1.16 | 0.25 | 0.41 | 0.81 | 2.13 | 0.97 | 2.25 | 1.62 | 0.13 | 1.48 | 0.06 | 2.80 |
| N332 Panel | 22.5% | 65.2% | 20.2% | 13.5% | 44.9% | 55.1% | 75.3% | 53.9% | 36.0% | 22.5% | 55.1% | 52.8% | 43.8% | 44.9% | 25.8% | 9.0% | 24.7% |
| GMT IC50 (μg/mL) | 0.97 | 1.80 | 0.41 | 2.79 | 0.71 | 0.95 | 0.17 | 0.41 | 0.81 | 2.13 | 0.97 | 2.25 | 1.62 | 0.13 | 1.48 | 0.06 | 2.80 |

**Fig. S4. Breadth and potency of SHIV.5MUT-elicited V3-glycan bNAbs.** Neutralization breadth and potency of the SHIV.5MUT-elicited V3-glycan bNAbs was assessed using a panel of 130 tier-2 viruses consisting of a 119-virus global panel (blue) plus 11 additional viruses (purple).

Viruses are separated by subtype, and sequence features including V1 length, N332/N334 glycan status, and V3 epitope sequence are shown. Neutralization titers are expressed as $IC_{50}$ in µg/mL.

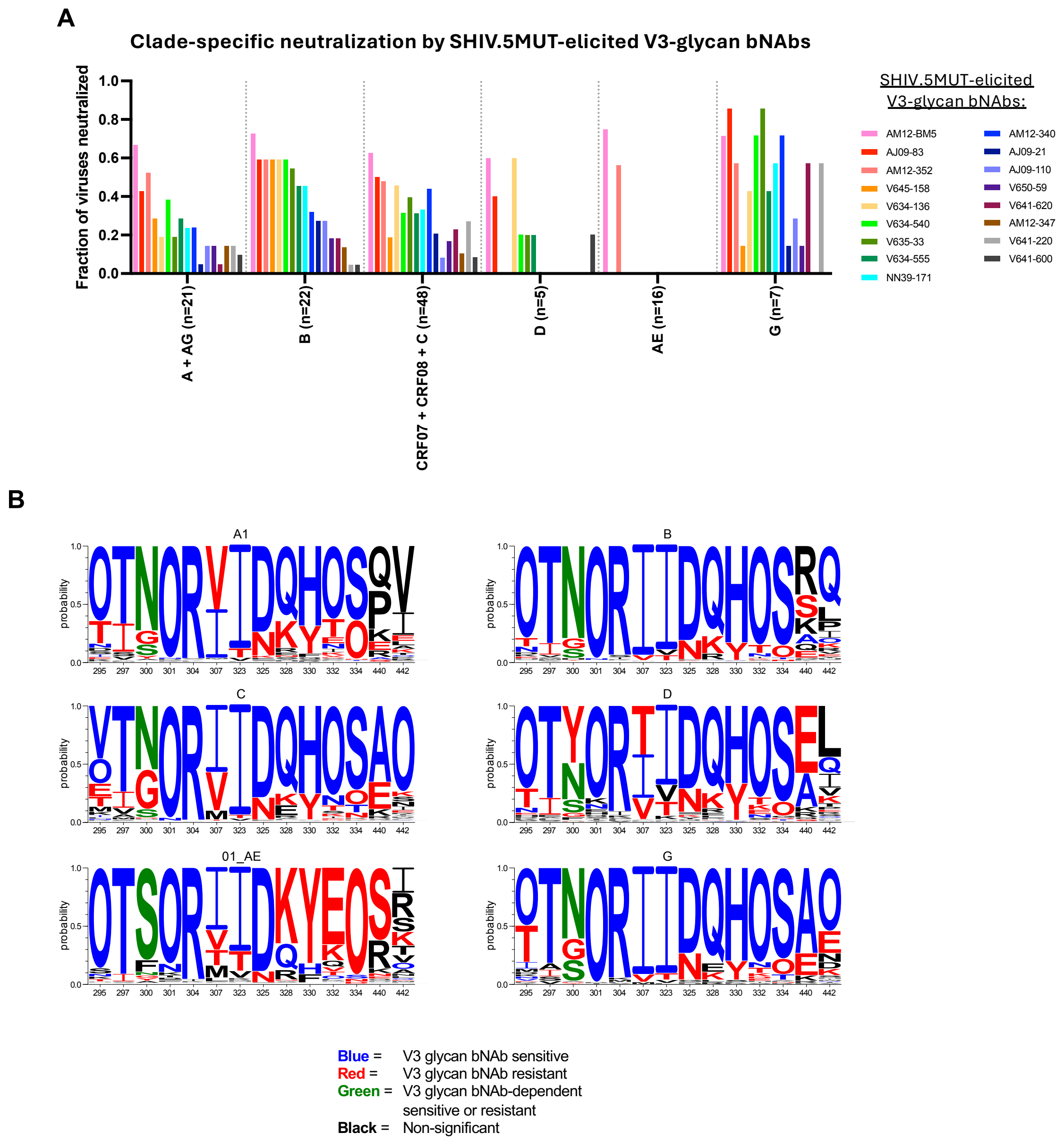


**Fig. S5. Clade-specific breadth of SHIV.5MUT-elicited V3-glycan bNAbs. (A)** Summary bar graph highlighting the fraction of heterologous viruses in each clade that is neutralized by the indicated antibody. CRF07 & CRF08 are clade C in Env, and CRF02 is clade A in Env, and hence these CRFs are grouped with the parental clades. **(B)** Sequence diversity at major V3-glycan bNAb epitope signature sites shown for six M-group clades (*21*). Logos were produced using the AnalyzeAlign tool on the Los Alamos HIV Database (www.hiv.lanl.gov) using the database alignment of M-group Envs.

**A**

| mAb ID | MLV | Q23 | | | | | | | CAP256SU | | | | |
|---|---|---|---|---|---|---|---|---|---|---|---|---|---|
| | | WT | D325N | R327K | H330Y | T334A | T334N | T303A | WT | D325A | H330Y | S334A | T303A |
| AM12-352 | >50 | 0.09 | 6.99 | >50 | 0.38 | 0.16 | 0.14 | >50 | 0.15 | >50 | 1.02 | 1.31 | >50 |
| AM12-340 | >50 | 0.05 | 1.49 | 18.46 | 0.88 | >50 | >50 | 0.01 | 7.53 | >50 | >50 | >50 | 1.63 |
| AM12-347 | >50 | 0.38 | 15.13 | >50 | >50 | >50 | >50 | 0.02 | 5.82 | >50 | >50 | >50 | 3.33 |

**B**

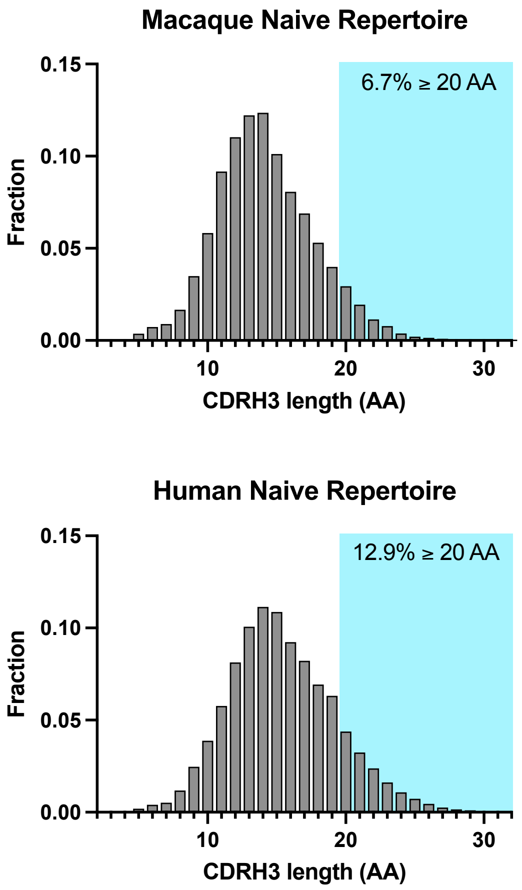


**Fig. S6. Representatives of three AM12 V3-glycan bNAb sub-lineages exhibit diversity in epitope recognition. (A)** Epitope mapping of representative members of three AM12 V3-glycan bNAb sub-lineages. Neutralization activity against a panel of Q23 and CAP256SU mutant viruses with mutations at key residues in the V3-glycan epitope is shown, with titers expressed as $IC_{50}$ in μg/mL. **(B)** Distribution of CDRH3 lengths in naïve $IgM^+$ peripheral B cells in humans (top) and macaques (bottom), identified by bulk BCR repertoire sequencing. Sequences with CDRH3 ≥20 AA are highlighted. The human dataset was taken from ref (*52*), and sequences from outlier participant 1070 were excluded. The rhesus dataset was generated in this study and includes 2,172,207 IgM sequences from eight macaques.

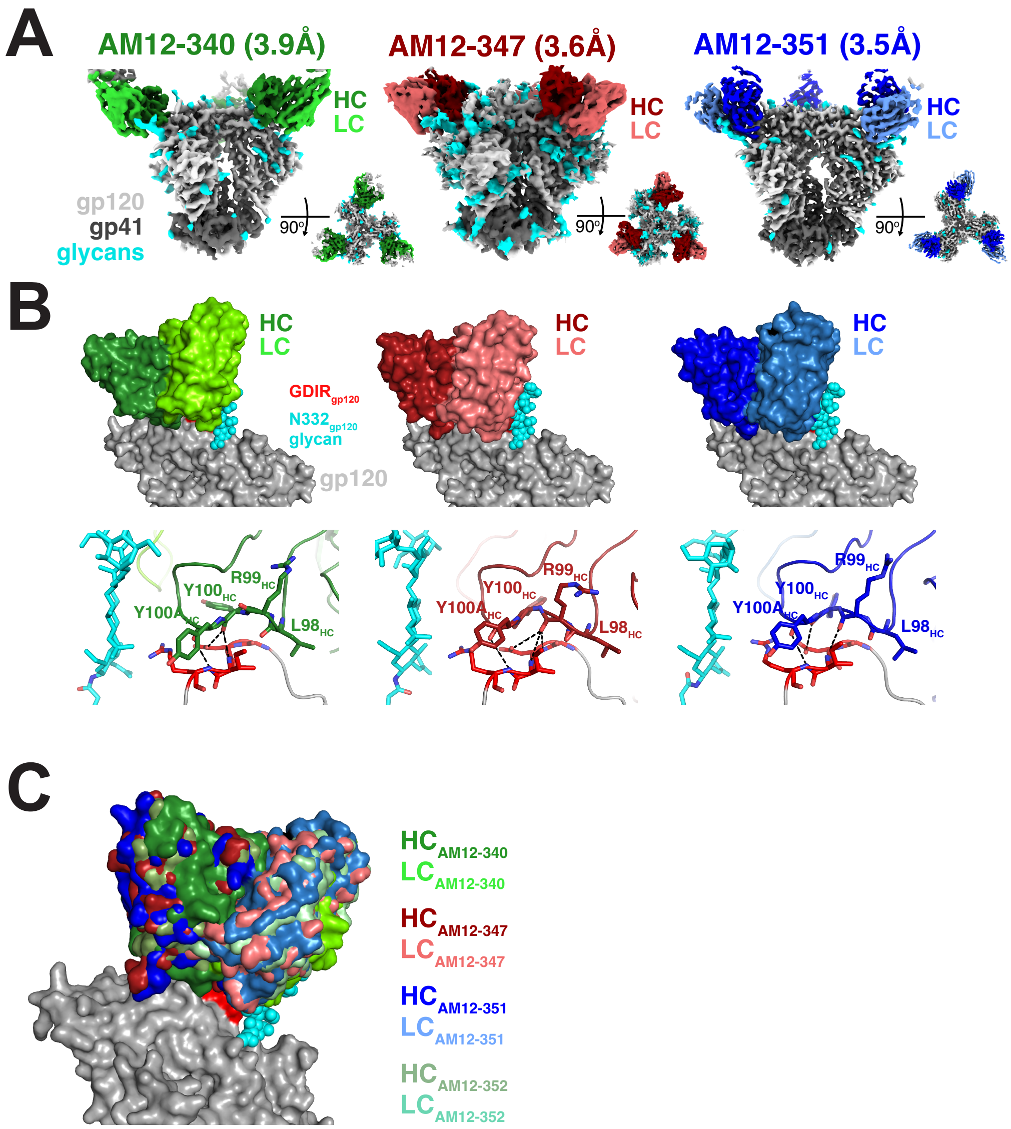


**Fig. S7. Structural characterization of AM12-352 lineage members. (A)** Cryo-EM maps of AM12-340 (left), AM12-347 (middle), and AM12-351 (right) bNAbs in complex with the 5MUT-3fill SOSIP. The heavy chain of each bNAb is shown in a darker shade, and the light chain is shown in a lighter shade. The N332 glycan is modeled as cyan spheres and the conserved GDIR peptide motif is red. **(B)** Top, surface representations of the V3-glycan bNAbs from **(A)** bound to gp120, with the N332 glycan and GDIR motif as depicted in **(A)**. Bottom, interfaces of the predicted interactions between the V3-glycan bNAbs and the GDIR motif. **(C)** Structural overlay of AM12 lineage members demonstrating nearly identical approach angles for binding to Env.

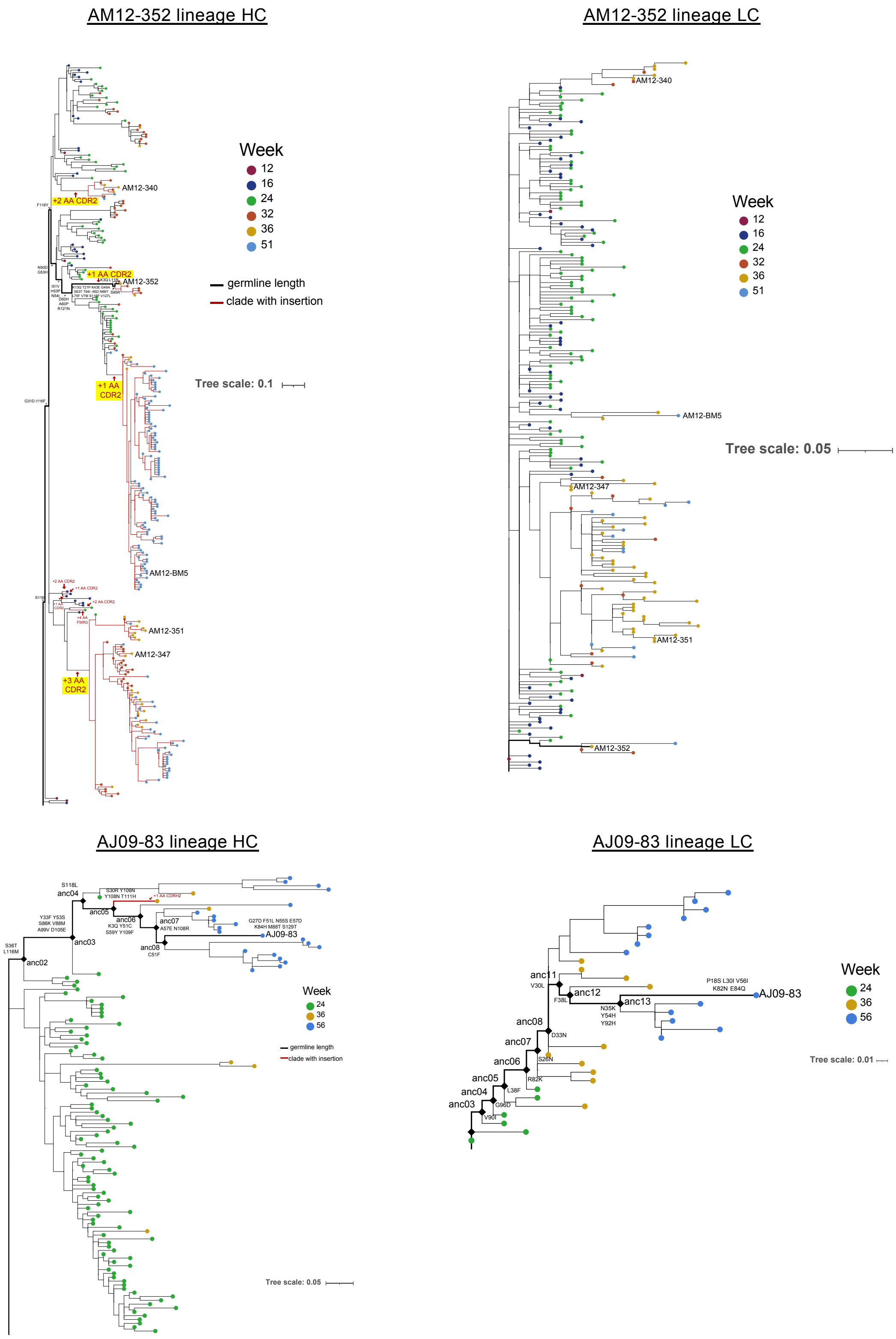


**Fig. S8. Evolution of the AM12-352 and AJ09-83 bNAb lineages.** Phylogenetic trees of heavy and light chain sequences derived from NGS and isolated mAbs are shown for two rhesus bNAb lineages whose UCAs did not bind to common V1 escape mutants **(Fig. 5F)**. Ancestors were inferred at each node by analyzing longitudinal bulk BCR sequencing data using the SONAR computational analysis pipeline. Sequences are color coded according to their week of isolation and the position of mature bNAbs on the trees are shown. Branches in the heavy chain tree on which amino acid insertions occurred are indicated in red, with four particular CDRH2 insertions highlighted. The bold black line highlights intermediary nodes to the mature bNAb. The tree was constructed from heavy and light chain nucleotide sequences using IgPhyML and the scale bar indicates the number of estimated mutations per site.

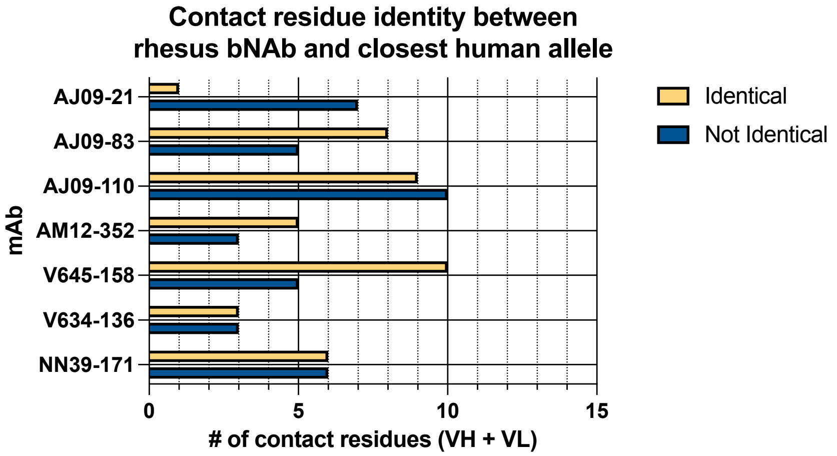


**Fig. S9. SHIV.5MUT-elicited bNAbs share the majority of their contact residues with the closest human V allele.** The number of contact residues that are (yellow) or are not (blue) identical in the closest human $V_H$ and $V_K$ alleles are shown for the indicated bNAbs.

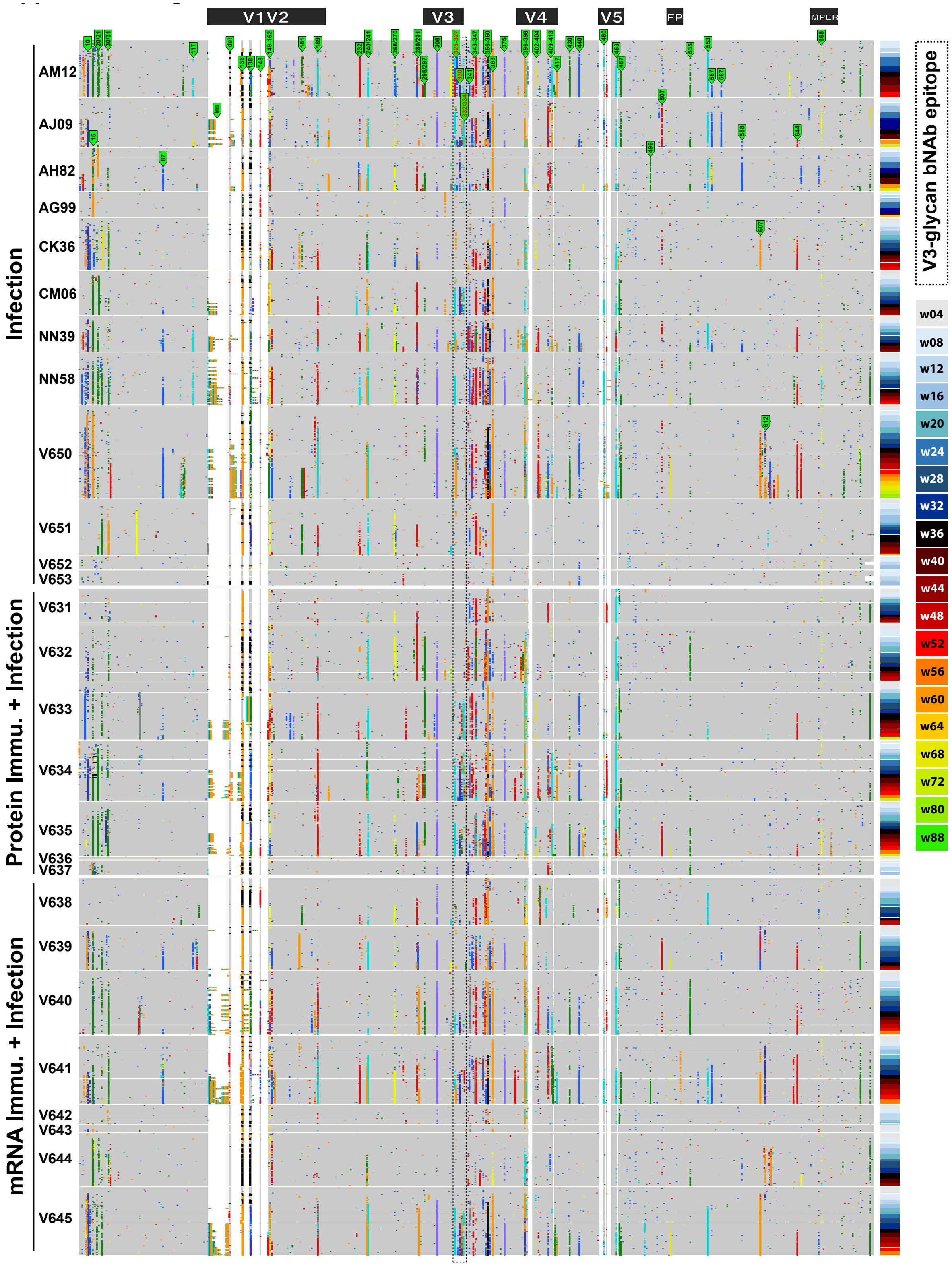


**Fig. S10. Immune selection in V1 and V3 regions in SHIV.5MUT-infected rhesus macaques.** A highlighter plot of longitudinal Env amino acid sequences obtained by single-genome sequencing of plasma viral RNA is shown for each macaque. Sequences are grouped first by macaque (left) and then by timepoint (right). Each row represents a single sequence, depicted as a string of horizontal pixels with each pixel representing a single amino acid in the alignment. Sequences are shown for the entire Env ectodomain, with a schematic map depicted at top. A single alignment was generated from sequences of all SHIV.5MUT-infected macaques at all timepoints, such that the vertical "stripes" indicate mutations at the same site in different Envs. All sequences are aligned to the SHIV.5MUT Env sequence. Residues with substantial selection are indicated, with colors denoting different amino acids. Timepoints are color-coded on the right. Deletions are denoted by black boxes.

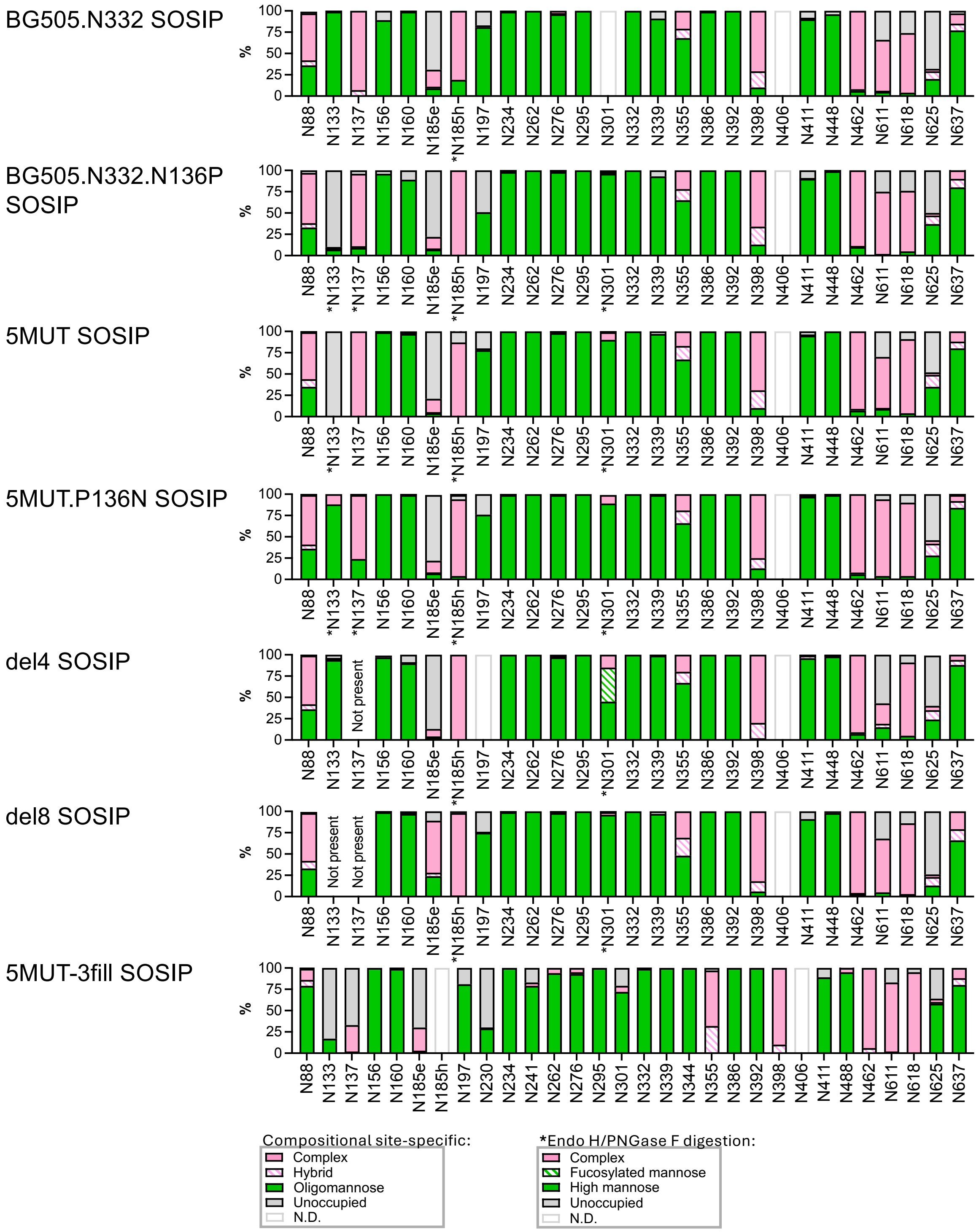


**Fig. S11. Hypoglycosylation of position N133 in the 5MUT Env trimer.** Mass spectroscopy-based site-specific glycan analysis of MD39-stabilized BG505.N332, BG505.N332.N136P, 5MUT, 5MUT.P136N, del4, del8, and 5MUT-3fill SOSIP proteins. Glycan compositions are grouped into their corresponding categories, with complex-type glycans displayed in pink, hybrid in hatched pink, oligomannose in green, and unoccupied in gray. For sites in which the site-specific data was deduced by employing glycosidase digestion by Endo H followed by PNGase F in O18 water (indicated by asterisks), the peptide modifications which correspond to distinct glycan types are grouped into high mannose in green, fucosylated mannose in hatched green, complex in pink, and unoccupied in gray. Glycan sites that could not be determined are denoted as “N.D.”

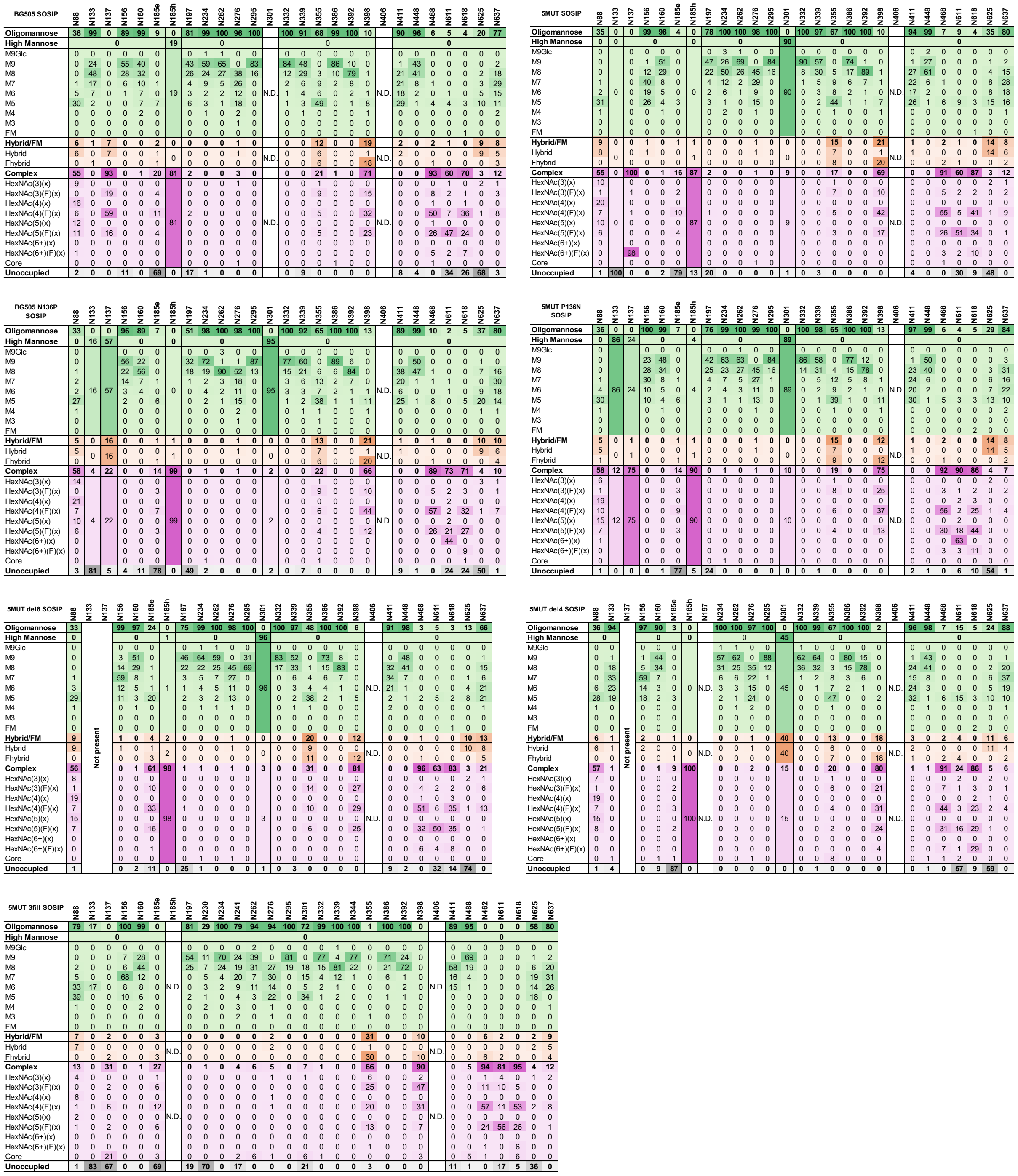


**Fig. S12. Compositional site-specific glycan analysis of various BG505- and 5MUT-derived trimers.** Quantification of site-specific occupancy and compositional glycan analysis of MD39-stabilized BG505.N332, BG505.N332.N136P, 5MUT, 5MUT.P136N, del4, del8, and 5MUT-3fill SOSIP proteins. The datasets show the glycoforms found at each potential N-linked glycosylation site (PNGS). Data for oligomannose glycans are shaded in green, hybrid-type glycans in orange, fully processed complex-type glycans in magenta, and the absence of a glycan from some PNGS in gray. Oligomannose-type glycans are categorized according to the number of mannose residues present, hybrid-type glycans by the presence/absence of fucose, and complex-type glycans by the number of processed antenna and the presence/absence of fucose. For further information see **Methods**. Data that was obtained using Endo H and PNGase F in O18 water cannot be allocated into the above categories. They are merged to cover all oligomannose/hybrid compositions or complex-type glycans.

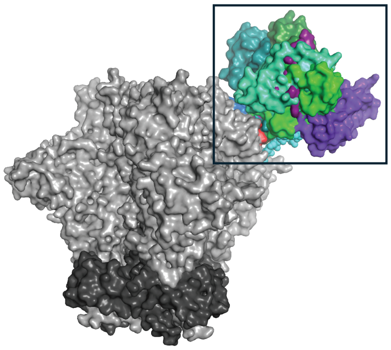

| gp120 region | gp120 residue | (wk 12) NN39-25 | (wk 32) NN39-171 | (5T3Z) 10-1074 |
|---|---|---|---|---|
| V1 | 134 | Y | | |
| | 135 | T | | T |
| | 136 | P | | |
| | 137 | N | N | N |
| | 138 | L | | |
| | 139 | T | T | |
| | 140 | N | N | |
| | 141 | | D | |
| | 142 | | | |
| | 143 | R | | |
| V3 | 321 | D | | |
| | 322 | | I | |
| | 323 | I | | |
| | 324 | G | G | G |
| | 325 | D | D | D |
| | 326 | I | I | I |
| | 327 | | R | R |
| | 328 | | Q | Q |
| | 329 | | | |
| | 330 | | H | H |
| V4 | 415 | | T | |
| | 416 | | | |
| | 417 | | P | |

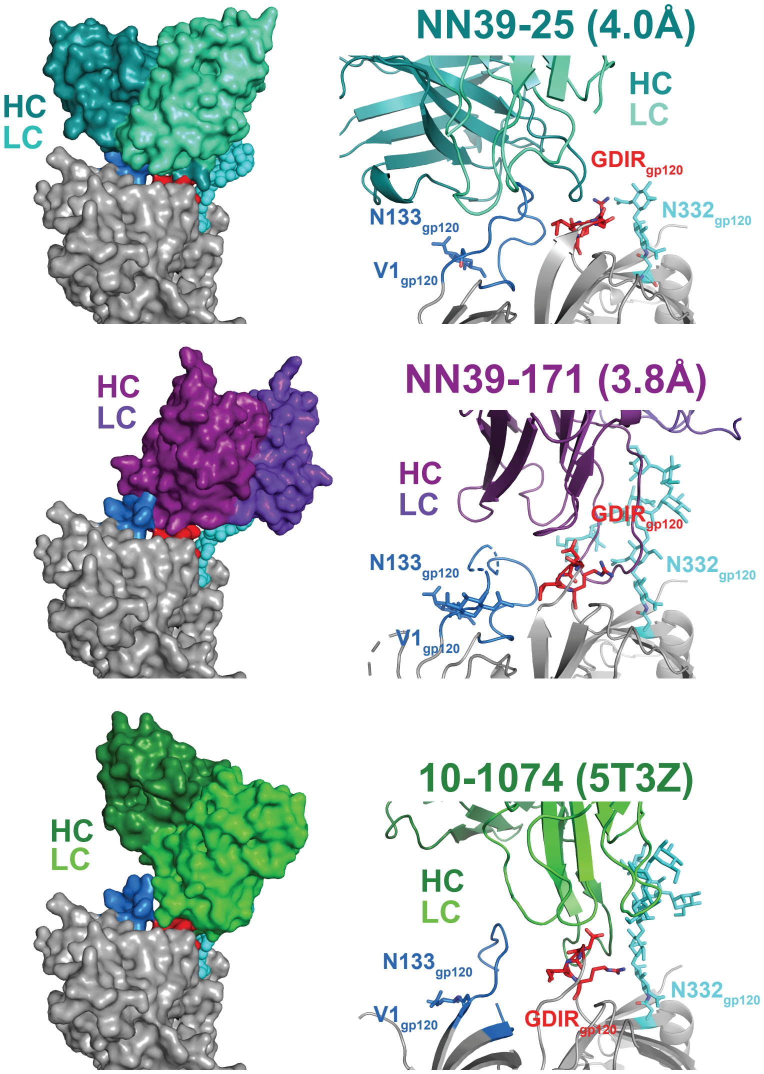


**Fig. S13. Comparative structural analysis of SHIV.5MUT-induced V1- and V3-directed neutralizing antibodies.** Cryo-EM maps of two SHIV.5MUT-induced neutralizing antibodies (NN39-25 and NN39-171) in complex with 5MUT-3fill SOSIP, overlaid with a previously-published crystal structure of the human V3-glycan bNAb 10-1074 in complex with BG505 SOSIP. The heavy chain of each antibody is shown in a darker shade, while the light chain is shown in a lighter shade. The $N332_{gp120}$ glycan is shown in cyan, the conserved GDIR peptide motif in red, and the V1 loop (including the $N133_{gp120}$ glycan) in dark blue. NN39-25 exhibits strong V1 loop dependence, whereas NN39-171 and 10-1074 bNAbs bind predominantly to the V3 loop. Contact residues are shown in the table, with red font indicating the four residues differentiating 5MUT from BG505.

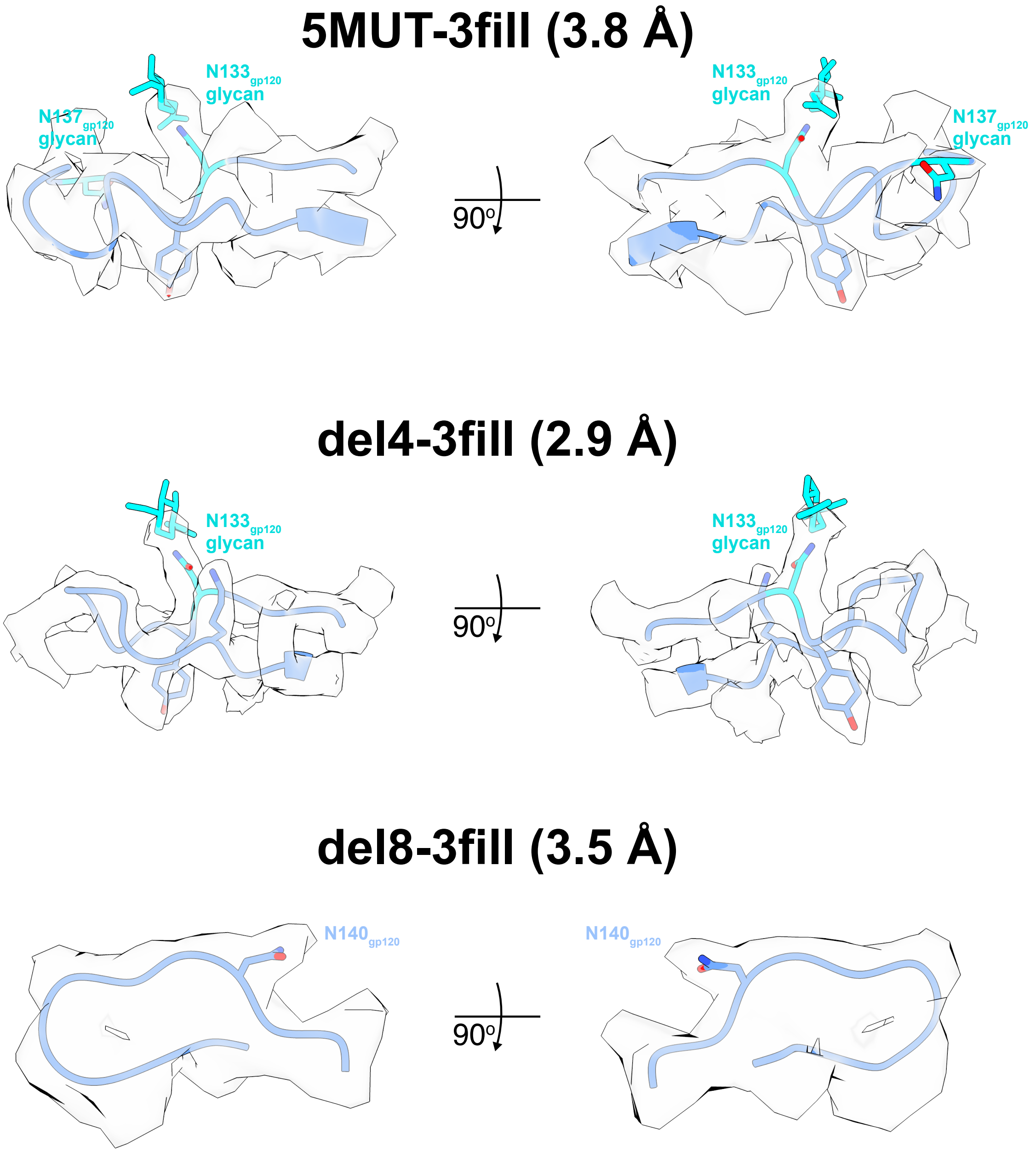


**Fig. S14. 5MUT, del4, and del8 trimers exhibit minimal V1 glycosylation.** Electron density maps showing minimal to no glycosylation within the V1 loops of 5MUT-3fill (top), del4-3fill (middle), and del8-3fill (bottom) SOSIPs.

**A**

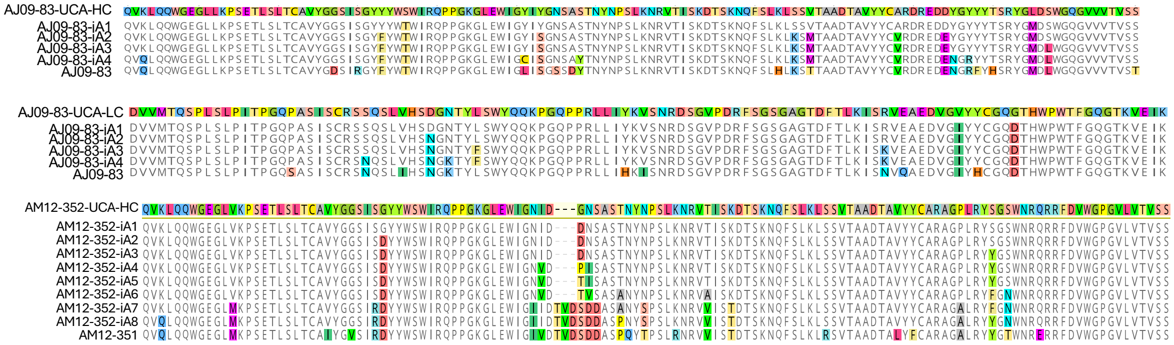


**B**

| mAb ID | BG505. N332 | 5MUT V1 variants | | | | |
|---|---|---|---|---|---|---|
| | | P136S. R143K | 5MUT | del3 | del4 | del8 |
| AM12-352-UCA | 1.09 | 1.07 | 0.91 | 0.74 | 1.00 | 0.95 |
| AM12-352-iA1 | 0.88 | 0.87 | 0.89 | 0.87 | 0.90 | 0.86 |
| AM12-352-iA2 | 0.93 | 0.94 | 1.01 | 1.09 | 0.87 | 0.93 |
| AM12-352-iA3 | 1.10 | 1.06 | 191.0 | 14.78 | 8.58 | 0.98 |
| AM12-352-iA4 | 1.06 | 0.94 | 155.3 | 16.06 | 9.52 | 0.98 |
| AM12-352-iA5 | 0.89 | 0.89 | 131.0 | 12.13 | 7.12 | 0.84 |
| AM12-352-iA6 | 1.01 | 1.00 | 176.5 | 18.59 | 10.43 | 0.91 |
| AM12-352-iA7 | 6.32 | 23.71 | 622.3 | 363.3 | 266.0 | 2.24 |
| AM12-352-iA8 | 51.07 | 120.4 | 921.3 | 591.4 | 356.4 | 15.63 |
| AJ09-83-UCA | 1.70 | 0.99 | 0.95 | 0.95 | 0.95 | 0.96 |
| AJ09-83-iA1 | 0.85 | 0.88 | 0.91 | 0.87 | 0.92 | 0.98 |
| AJ09-83-iA2 | 0.96 | 0.93 | 1.04 | 38.48 | 22.61 | 2.69 |
| AJ09-83-iA3 | 0.98 | 49.51 | 232.0 | 599.4 | 386.7 | 260.1 |
| AJ09-83-iA4 | 47.64 | 108.4 | 251.1 | 549.6 | 366.9 | 536.4 |

| MFI Fold Change over Mock |
|---|
| 0-3 |
| 3-10 |
| 10-100 |
| 100-1000 |

**Fig. S15. Early intermediate antibodies in the AM12-352 and AJ09-83 lineages bind common SHIV.5MUT escape mutants that have shortened V1 loops. (A)** Amino acid alignment of inferred unmutated common ancestor (UCA) and intermediate ancestors (iA1-8) in the AJ09-83 and AM12-352 lineages. Inferred ancestors for the heavy and light chains of the AJ09-83 lineage are shown, while only inferred ancestors for the heavy chain of the AM12-352 lineage are shown. **(B)** Representative heavy and light chain sequences were selected for synthesis and artificially paired. AJ09-83 intermediate heavy chains were paired with intermediate light chains from a similar developmental stage. AM12-352 intermediate heavy chains were paired with the AM12-352-UCA light chain. mRNA-encoded Env trimers were expressed on the surface of 293F cells and binding to UCA and intermediate ancestor mAbs was assessed by flow cytometry. Binding is quantified as fold-change in mean fluorescence intensity (MFI) over mock-transfected cells, and data points with a >3-fold increase are colored as indicated.

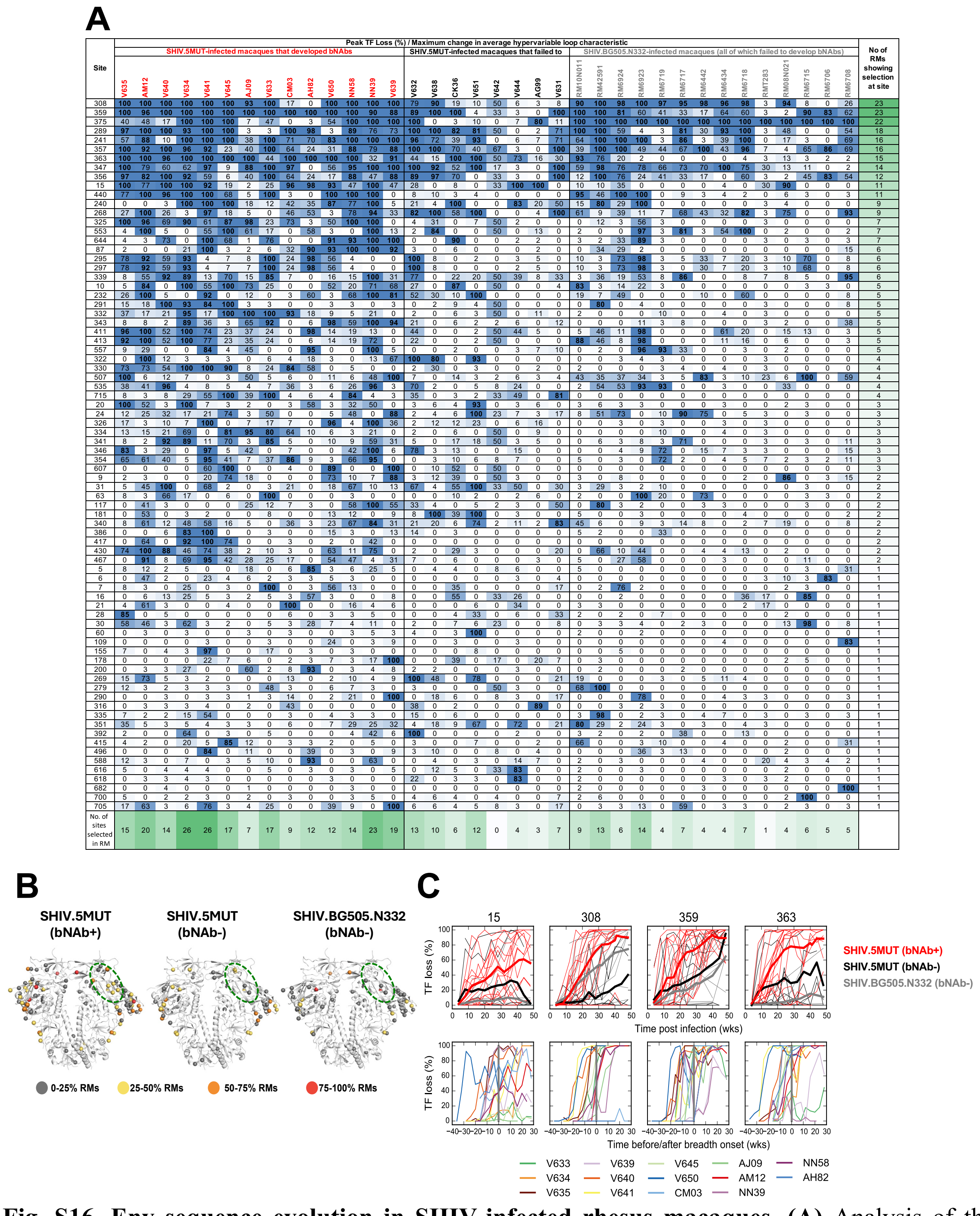


**Fig. S16. Env sequence evolution in SHIV-infected rhesus macaques. (A)** Analysis of the evolving Env quasispecies in all SHIV-infected macaques for sites under immune selection. Three groups, including 14 SHIV.5MUT-infected animals that developed bNAbs, 8 SHIV.5MUT-infected animals that did not develop bNAbs, and 14 SHIV.BG505.N332-infected animals that did not develop bNAbs, were subjected to LASSIE analysis, which identifies residues where mutations have altered the encoded amino acid in at least 80% of sequences at one or more timepoints. All

LASSIE-selected sites are highlighted in blue, with the number indicating the maximum percentage of sequences from any timepoint that exhibit loss of the transmitted founder (TF) residue for each macaque. **(B)** Structural mapping of LASSIE-selected sites in each group onto the BG505 Env (PDB ID: 9EHL). Sites are color-coded as indicated based on recurrence in multiple macaques. The green ellipse demarcates the approximate location of the V3-glycan bNAb epitope. **(C)** Same analysis as shown in **Fig. 6C**, but for sites that were not temporally associated with the acquisition of plasma breadth. Top: Longitudinal mutation frequency (%TF loss) at four sites enriched in SHIV.5MUT-infected macaques that developed bNAbs. Red and black curves denote SHIV.5MUT-infected macaques that did versus did not develop bNAbs, respectively, while grey curves indicate SHIV.BG505.N332-infected animals (all of which failed to develop bNAbs). Thin lines indicate data from individual animals and thick lines represent group averages. Bottom: Mutation frequency (%TF loss) in macaques that developed bNAbs, with the timepoint at which plasma breadth is first detected (defined as at least 1 heterologous virus with reciprocal $ID_{50} > 20$) set to zero weeks (wks). Different colors indicate different macaques. TF, transmitted founder.

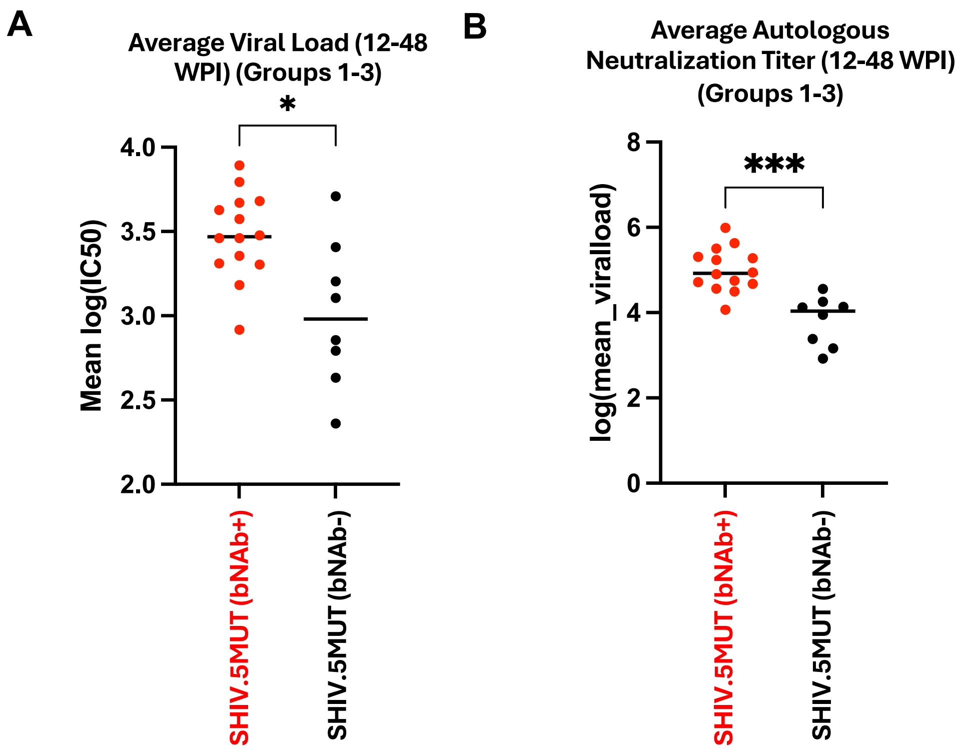


**Fig. S17. bNAb development in SHIV.5MUT-infected macaques is associated with higher viral loads and autologous neutralization titers. (A)** Average viral loads in SHIV.5MUT-infected macaques (Groups 1-3) between weeks 12-48 post-infection, quantified as log(mean viral load). **(B)** Average autologous neutralization titers in SHIV.5MUT-infected macaques (Groups 1-3), represented as mean(log($IC_{50}$)). Mann-Whitney U test was used. Not significant (ns) $P > 0.05$; $^{*}P < 0.05$; $^{**}P < 0.01$; $^{***}P < 0.001$.

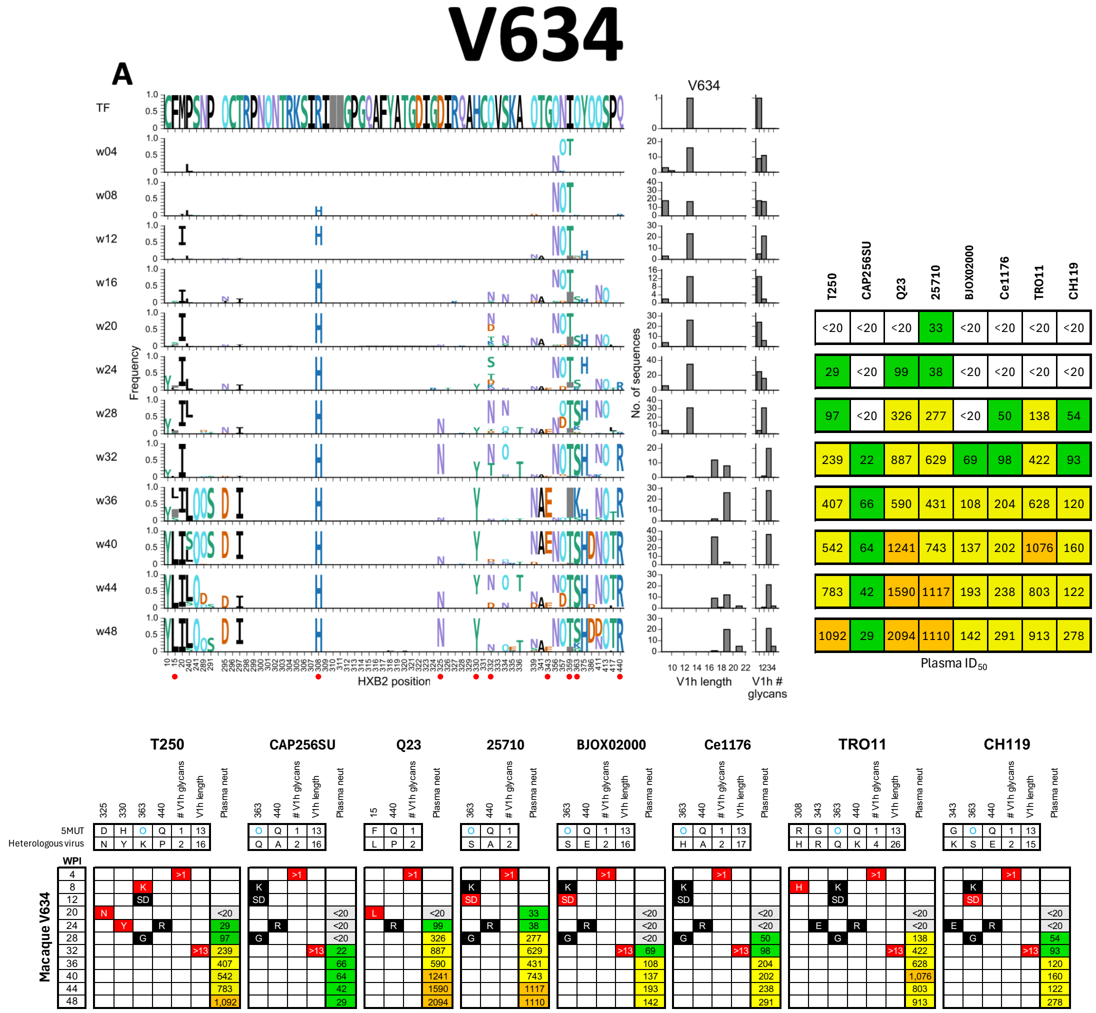


**Fig. S18. Env evolution and development of plasma breadth in rhesus macaque V634.** See **Fig. 6A** and **Fig. 7B** for details regarding figure layout and **Methods** for a description of the analysis. **(A)** Longitudinal patterns of mutations in Env and development of neutralization breadth. **(B)** Integrated information from the three graphs in **(A)**. 20/21 amino acid changes in key sites, 4/6 V1h elongations, and 8/8 V1h additions relevant to a given pseudovirus were detected prior to neutralization breadth being observed in the plasma.

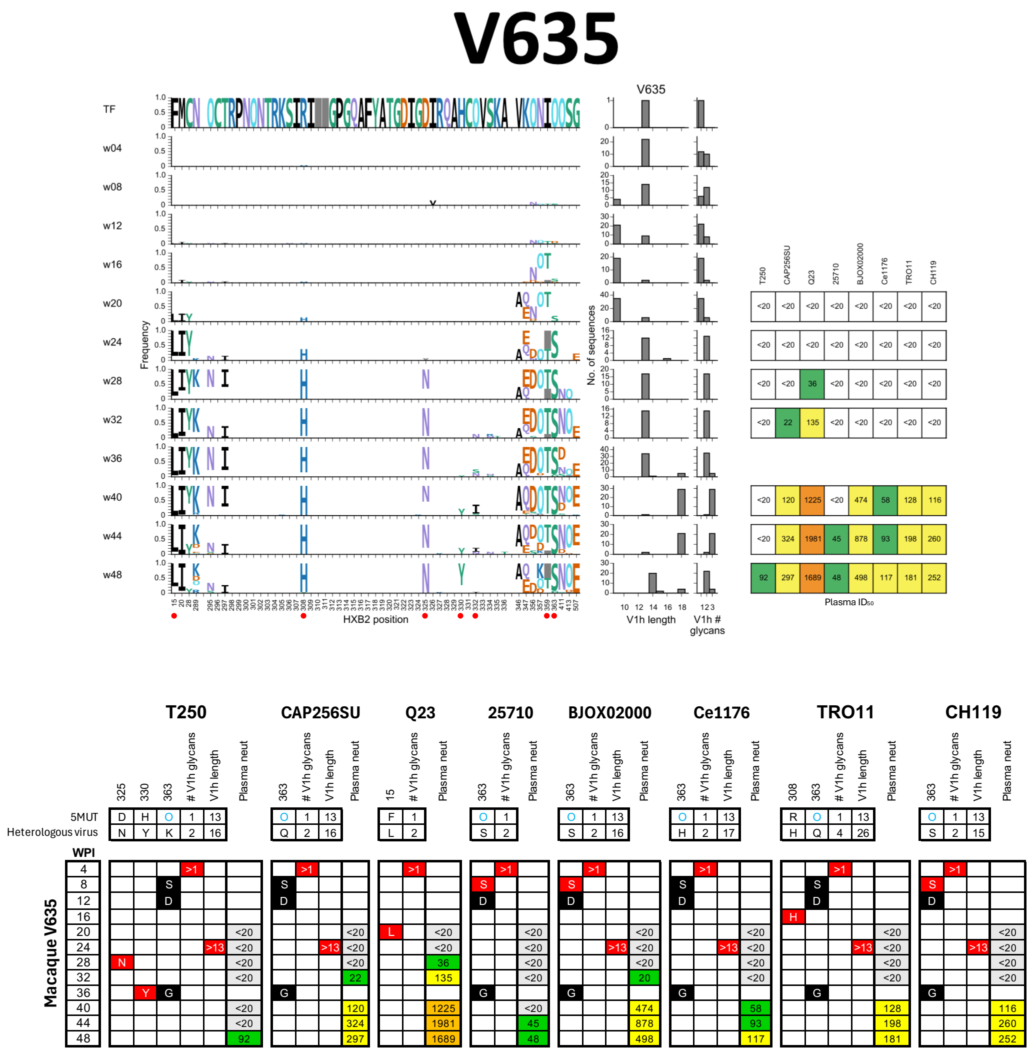


**Fig. S19. Env evolution and development of plasma breadth in rhesus macaque V635.** See **Fig. 6A** and **Fig. 7B** for details regarding figure layout and **Methods** for a description of the analysis. **(A)** Longitudinal patterns of mutations in Env and development of neutralization breadth. **(B)** Integrated information from the three graphs in **(A)**. 11/11 amino acid changes in key sites, 6/6 V1h elongations, and 8/8 V1h glycans additions relevant to a given pseudovirus were detected prior to neutralization breadth being observed in the plasma.

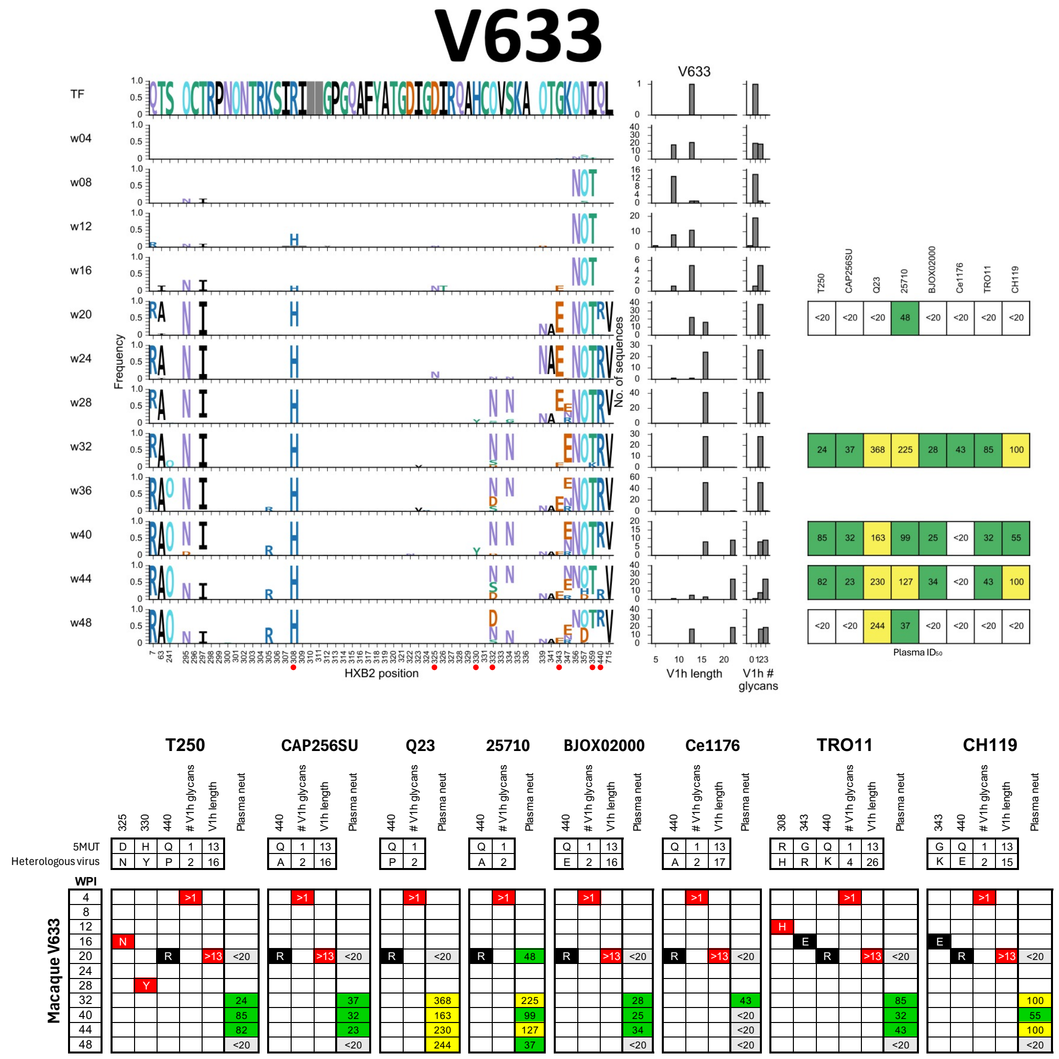


**Fig. S20. Env evolution and development of plasma breadth in rhesus macaque V633.** See **Fig. 6A** and **Fig. 7B** for details regarding figure layout and **Methods** for a description of the analysis. **(A)** Longitudinal patterns of mutations in Env and development of neutralization breadth. **(B)** Integrated information from the three graphs in **(A)**. 13/13 amino acid changes in key sites, 6/6 V1h elongations, and 8/8 V1h glycan additions relevant to a given pseudovirus were detected prior to or concurrent with neutralization breadth being observed in the plasma. The D325N and H330Y variants were rare but present.

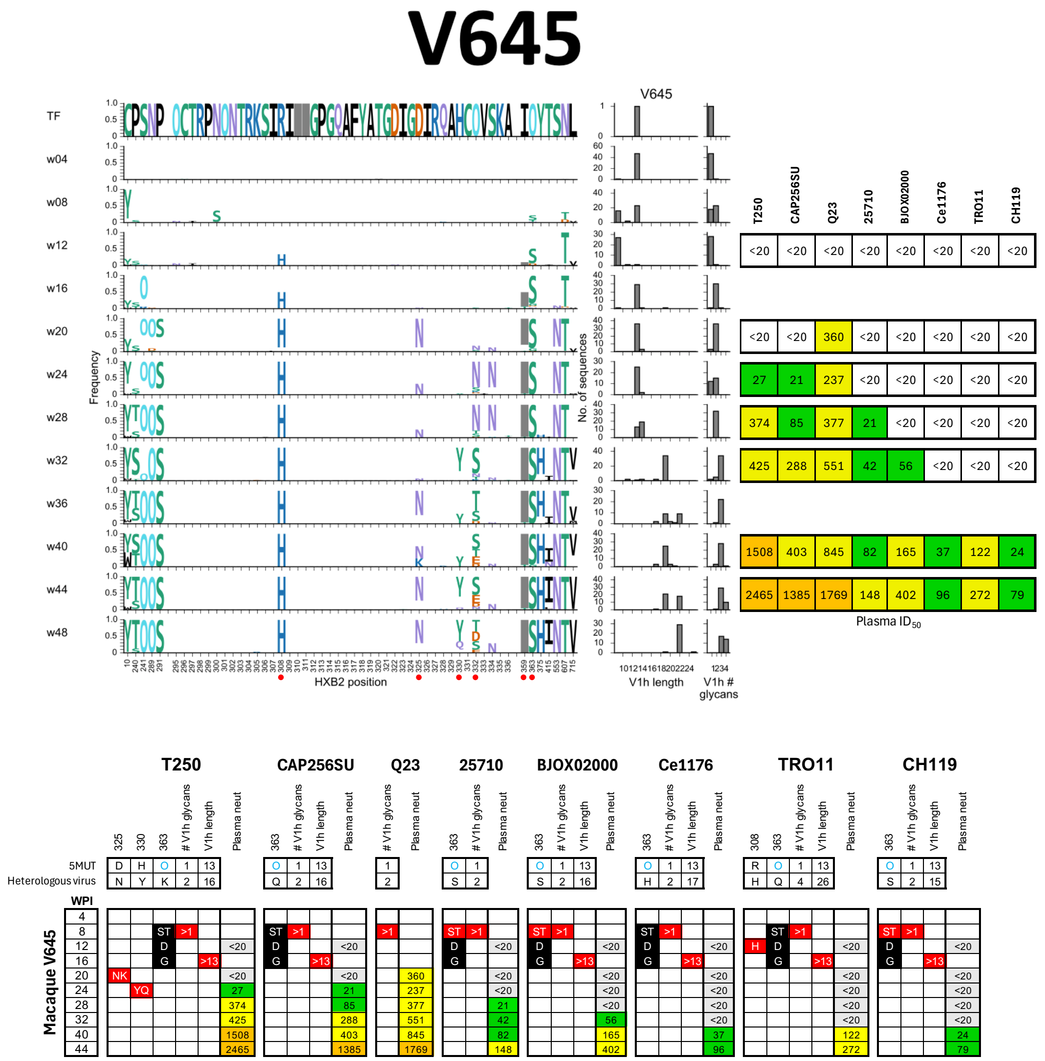


**Fig. S21. Env evolution and development of plasma breadth in rhesus macaque V645.** See **Fig. 6A** and **Fig. 7B** for details regarding figure layout and **Methods** for a description of the analysis. **(A)** Longitudinal patterns of mutations in Env and development of neutralization breadth. **(B)** Integrated information from the three graphs in **(A)**. 10/10 amino acid changes in key sites, 6/6 V1h elongations, and 8/8 V1h glycan additions relevant to a given pseudovirus were detected prior to or concurrent with neutralization breadth being observed in the plasma. O363S was rare at week 8 but common by week 16. Increases in V1h length were initially rare at week 16 but became common by week 32, perhaps contributing to bNAb selection for breadth and potency. Plasma neutralization of Q23 was detected first, perhaps in part because it has a short V1h region and a glycan at O363, much like 5MUT.

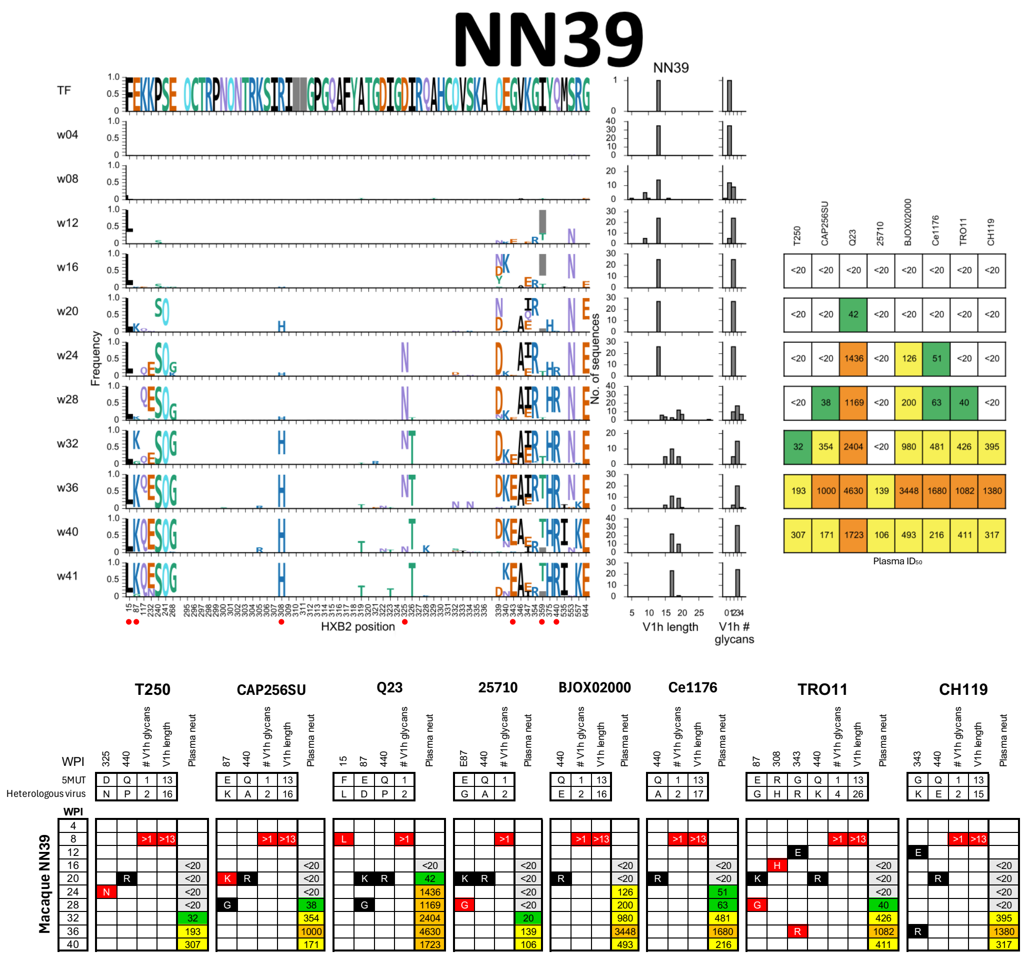


**Fig. S22. Env evolution and development of plasma breadth in rhesus macaque NN39.** See **Fig. 6A** and **Fig. 7B** for details regarding figure layout and **Methods** for a description of the analysis. **(A)** Longitudinal patterns of mutations in Env and development of neutralization breadth. **(B)** Integrated information from the three graphs in **(A)**. 17/17 amino acid changes, 6/6 V1h elongations, and 8/8 V1h glycan additions relevant to a given pseudovirus were detected prior to or concurrent with neutralization breadth being observed in the plasma. To recognize 25710 and TRO11, emergence of the E87G variant may be required, rather than simply the loss of the 5MUT-encoded E87 residue. Similarly, V1 elongation started at week 8 but was rare until week 28, which coincides with the timepoint at which neutralization potency increased substantially against heterologous viruses with longer V1h loops. Plasma neutralization of Q23 was detected earlier at week 20, perhaps in part because it has a short V1h region much like 5MUT.

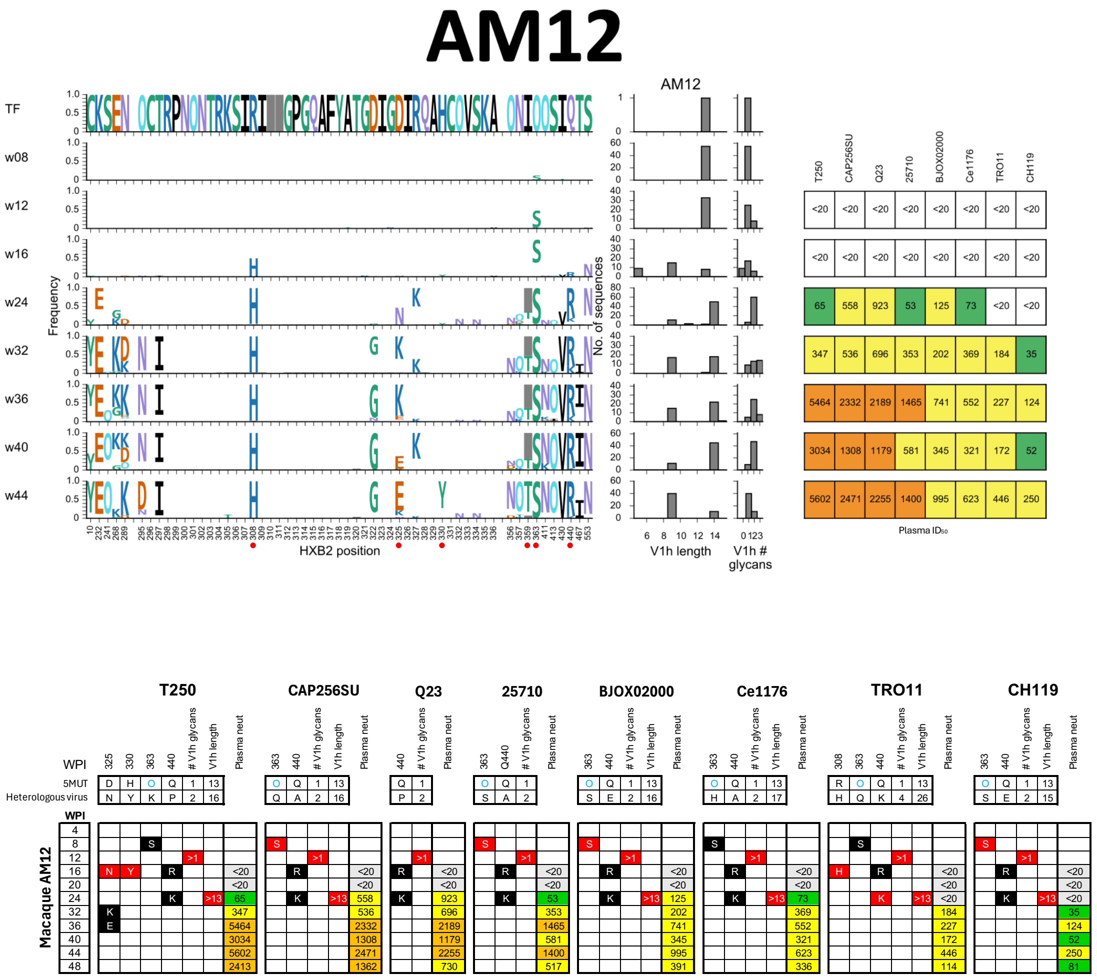


**Fig. S23. Env evolution and development of plasma breadth in rhesus macaque AM12.** See **Fig. 6A** and **Fig. 7B** for details regarding figure layout and **Methods** for a description of the analysis. **(A)** Longitudinal patterns of mutations in Env and development of neutralization breadth. **(B)** Integrated information from the three graphs in **(A)**. 18/18 amino acid changes, 6/6 V1h elongations, and 8/8 V1h glycan additions relevant to a given pseudovirus were detected prior to or concurrent with neutralization breadth being observed in the plasma.

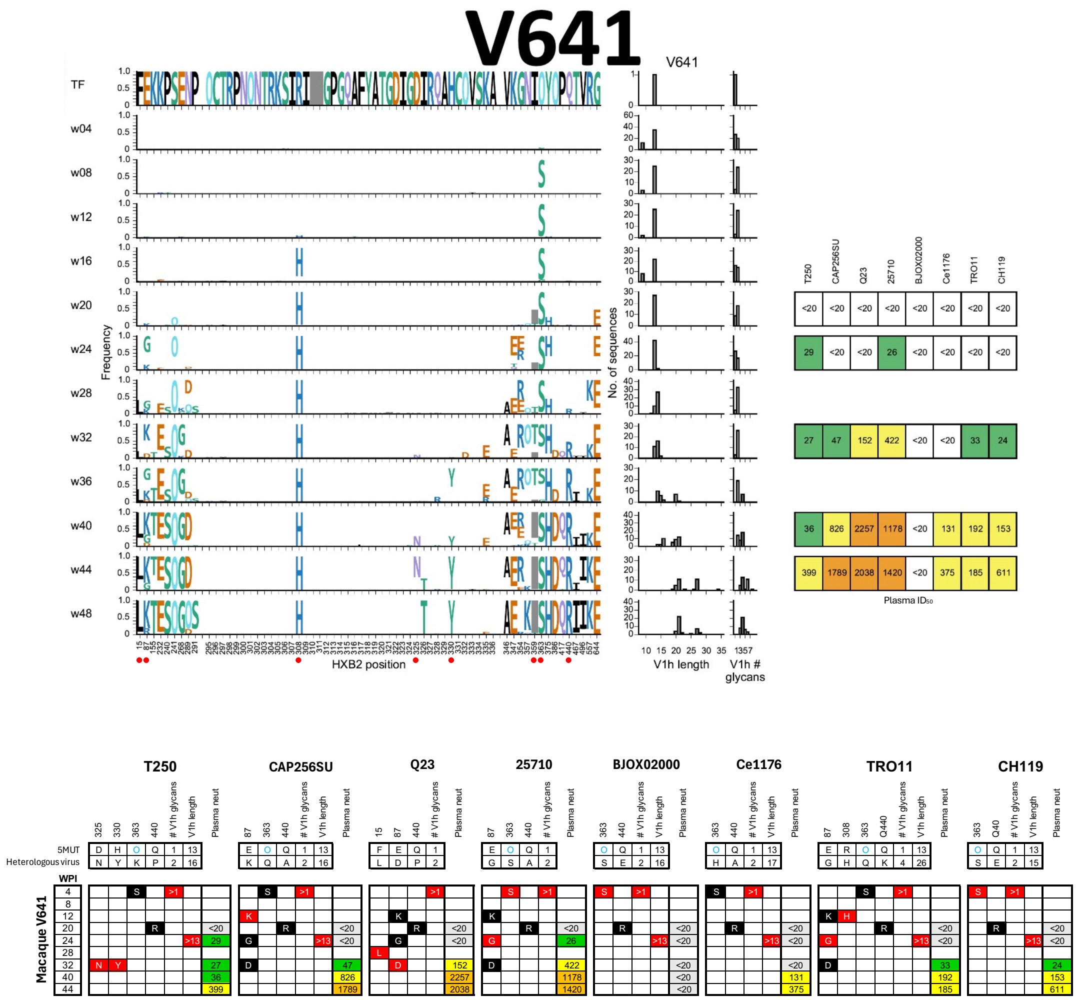


**Fig. S24. Env evolution and development of plasma breadth in rhesus macaque V641.** See **Fig. 6A** and **Fig. 7B** for details regarding figure layout and **Methods** for a description of the analysis. **(A)** Longitudinal patterns of mutations in Env and development of neutralization breadth. **(B)** Integrated information from the three graphs in **(A)**. 19/21 amino acid changes, 5/5 V1h elongations, and 7/7 V1h glycan additions relevant to a given pseudovirus were detected prior to or concurrent with neutralization breadth being observed in the plasma. The two exceptions were D325N and H330Y, which were first detected at week 32 despite T250 being weakly neutralized by the plasma at week 24. This potency increased by over 10-fold following subsequent emergence of the D325N and H330Y variants. For this animal, the specific residue at position 87 may be important for recognizing certain viruses, rather than simply the loss of the 5MUT-encoded E87 residue. Indeed, E87K/D/G variants were sampled in the quasispecies.

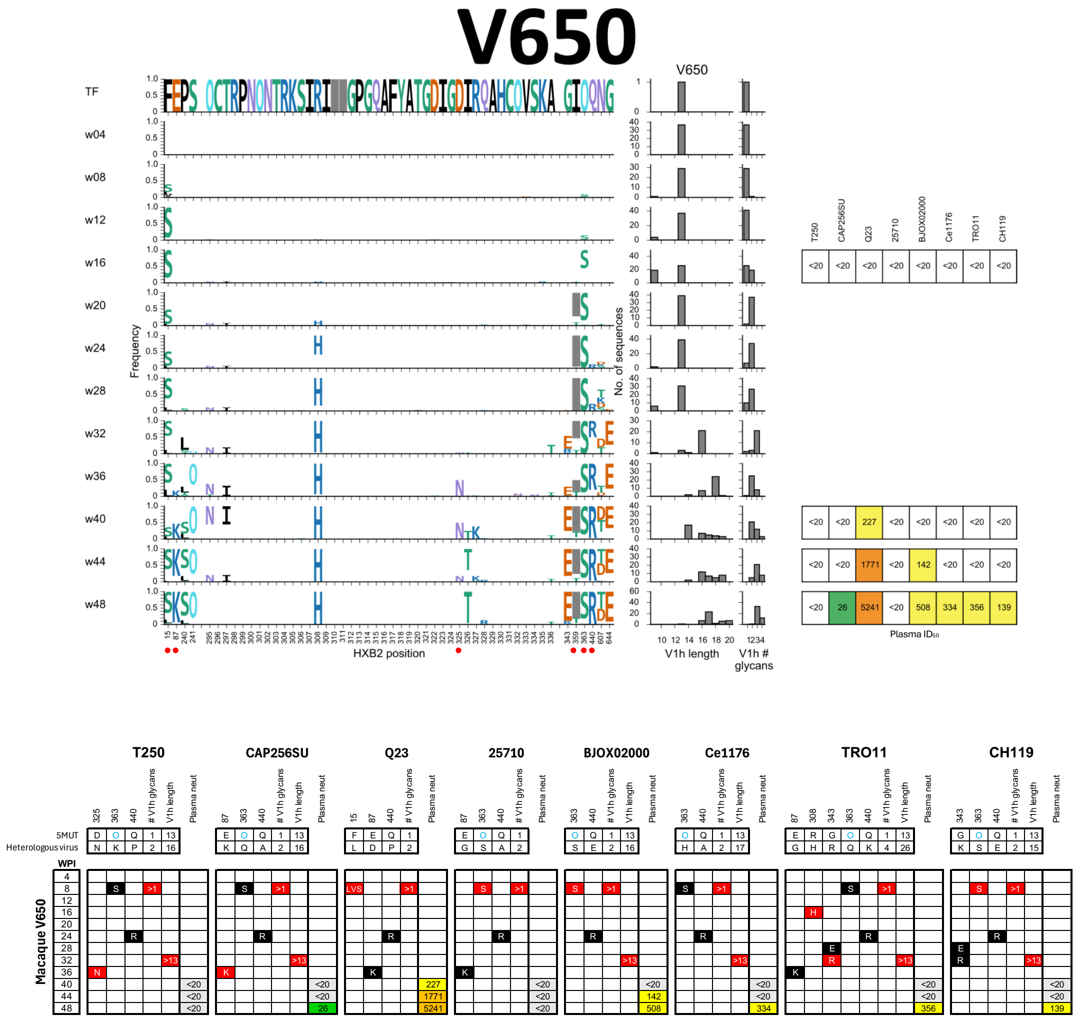

**Fig. S25. Env evolution and development of plasma breadth in rhesus macaque V650.** See **Fig. 6A** and **Fig. 7B** for details regarding figure layout and **Methods** for a description of the analysis. **(A)** Longitudinal patterns of mutations in Env and development of neutralization breadth. **(B)** Integrated information from the three graphs in **(A)**. 18/18 amino acid changes, 5/5 V1h elongations, and 6/6 V1h glycan additions relevant to a given pseudovirus were detected prior to or concurrent with neutralization breadth being observed in the plasma. O363S is very rare at week 8 but becomes common by week 20. V1h-elongated variants are common by week 32, prior to recognition of heterologous viruses with longer V1h lengths.

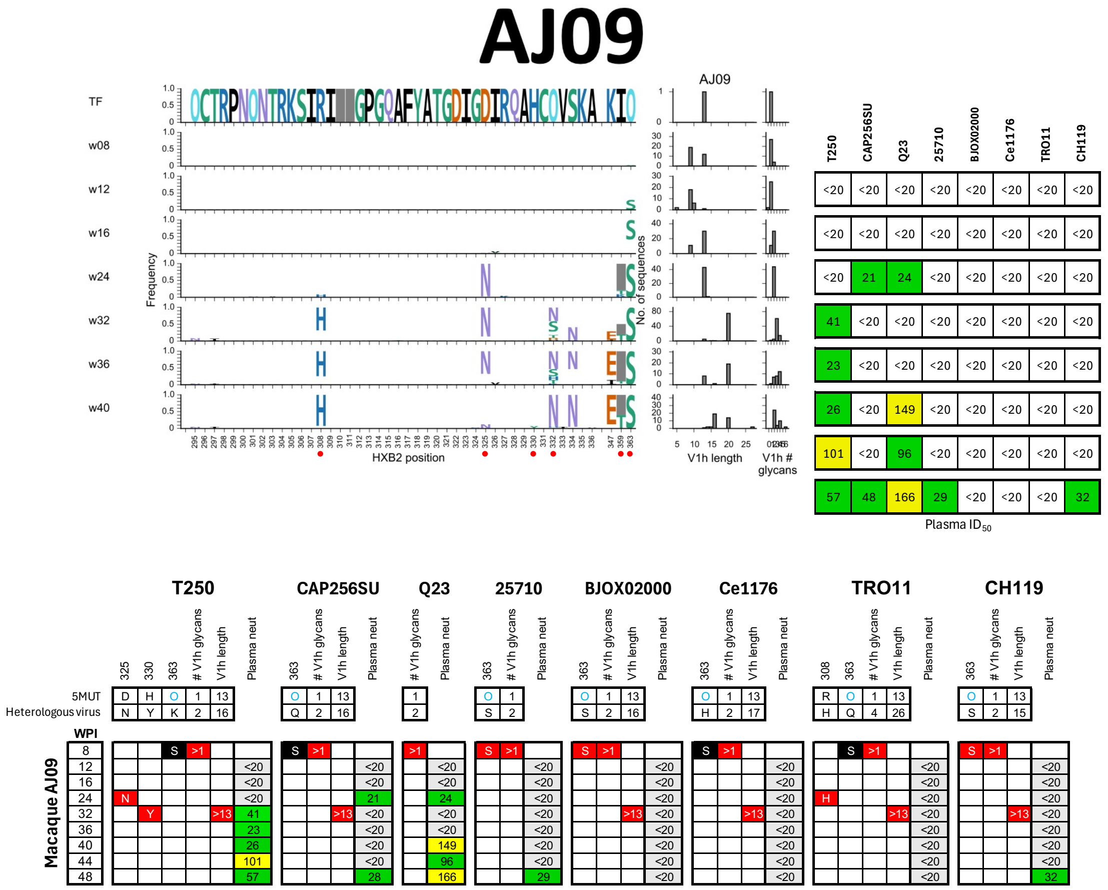


**Fig. S26. Env evolution and development of plasma breadth in rhesus macaque AJ09.** See **Fig. 6A** and **Fig. 7B** for details regarding figure layout and **Methods** for a description of the analysis. **(A)** Longitudinal patterns of mutations in Env and development of neutralization breadth. **(B)** Integrated information from the three graphs in **(A)**. 6/6 amino acid changes, 2/3 V1h elongations, and 5/5 V1h glycan additions relevant to a given pseudovirus were detected prior to or concurrent with neutralization breadth being observed in the plasma. CAP256SU, which has a long V1h region, was weakly neutralized prior to detection of V1h-elongated variants in the quasispecies. The rapid near-complete loss of the critical N332 glycan (O332) at week 32 may drive near-complete escape of the maturing bNAb lineage and contribute to the limited breadth and potency observed in the plasma.

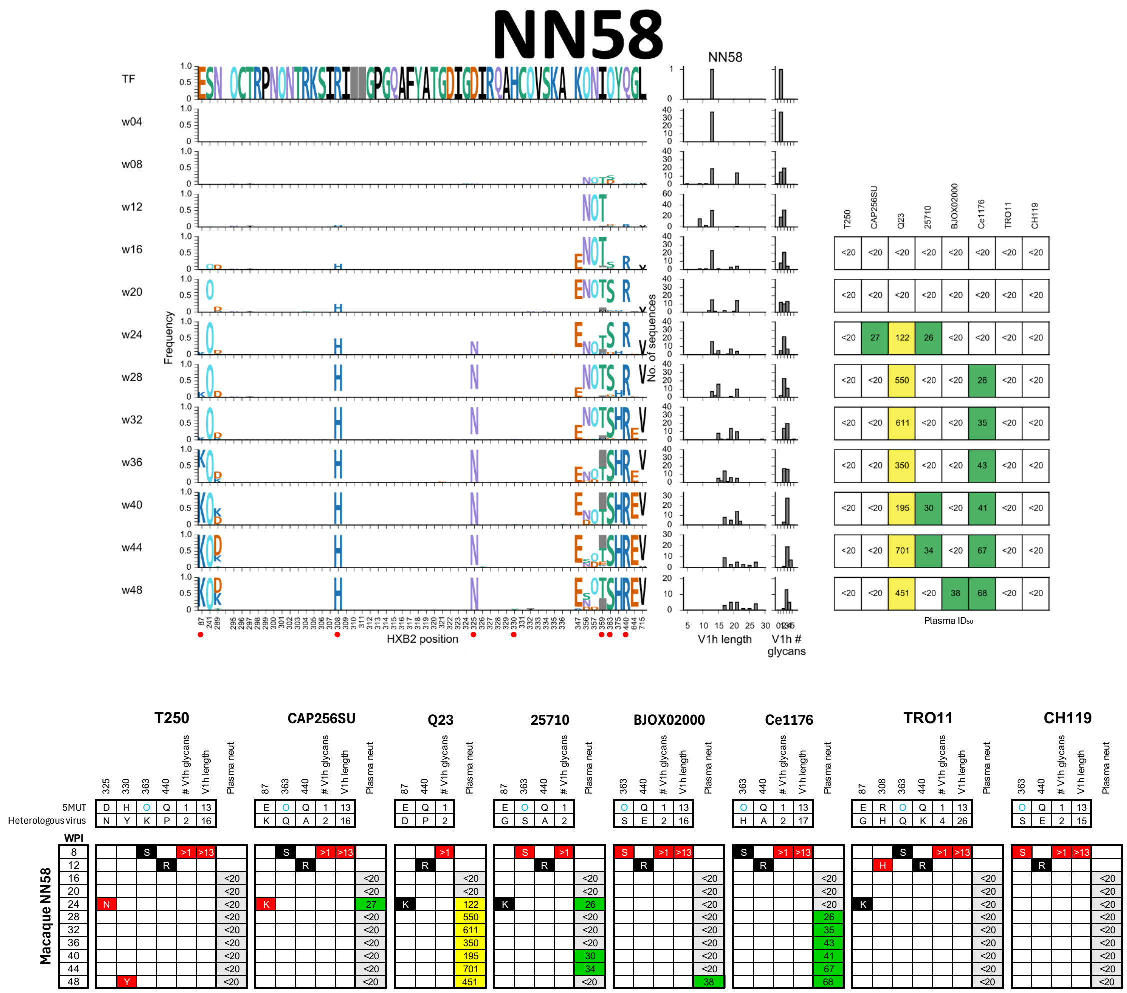


**Fig. S27. Env evolution and development of plasma breadth in rhesus macaque NN58.** See **Fig. 6A** and **Fig. 7B** for details regarding figure layout and **Methods** for a description of the analysis. **(A)** Longitudinal patterns of mutations in Env and development of neutralization breadth. **(B)** Integrated information from the three graphs in **(A)**. 12/12 amino acid changes, 3/3 V1h elongations, and 5/5 V1h glycan additions relevant to a given pseudovirus were detected prior to or concurrent with neutralization breadth being observed in the plasma.

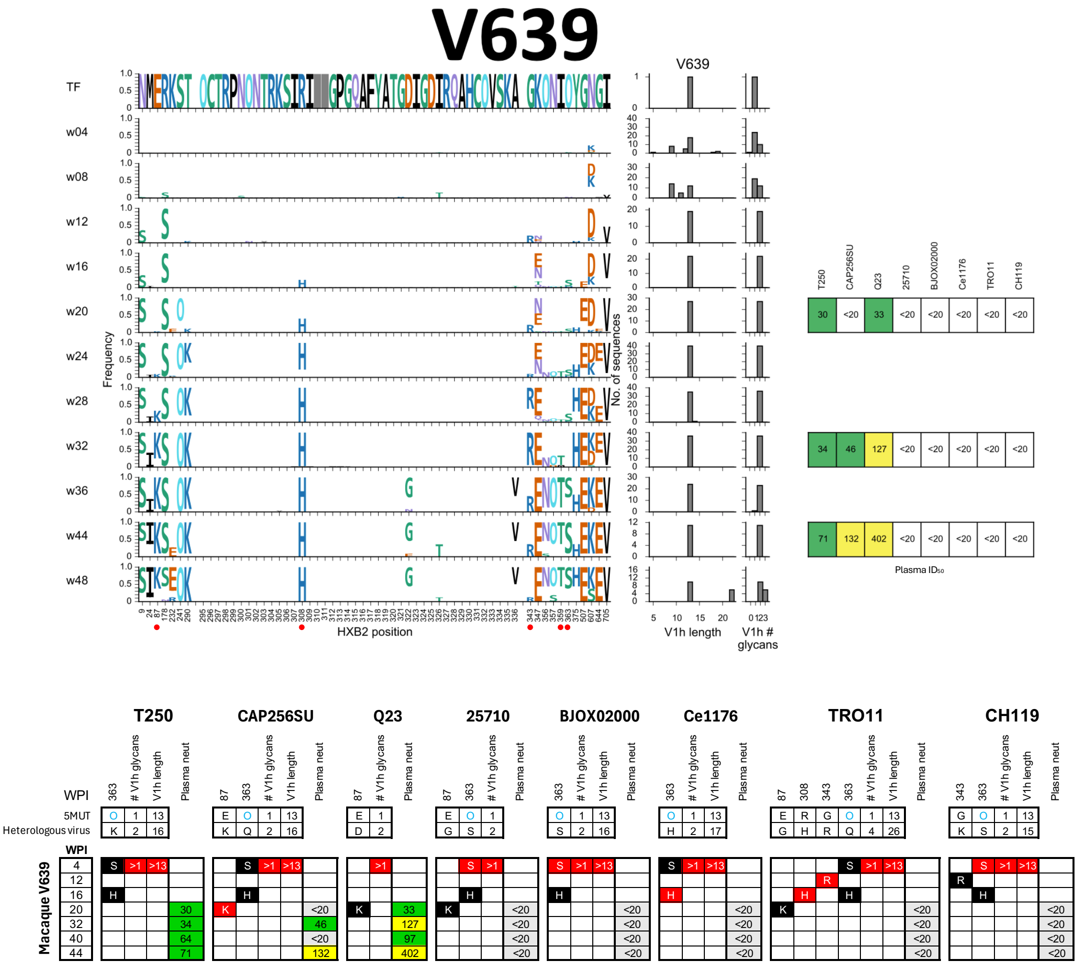


**Fig. S28. Env evolution and development of plasma breadth in rhesus macaque V639.** See **Fig. 6A** and **Fig. 7B** for details regarding figure layout and **Methods** for a description of the analysis. **(A)** Longitudinal patterns of mutations in Env and development of neutralization breadth. **(B)** Integrated information from the three graphs in **(A)**. 4/4 amino acid changes, 2/2 V1h elongations, and 3/3 V1h glycan additions relevant to a given pseudovirus were detected prior to or concurrent with neutralization breadth being observed in the plasma.

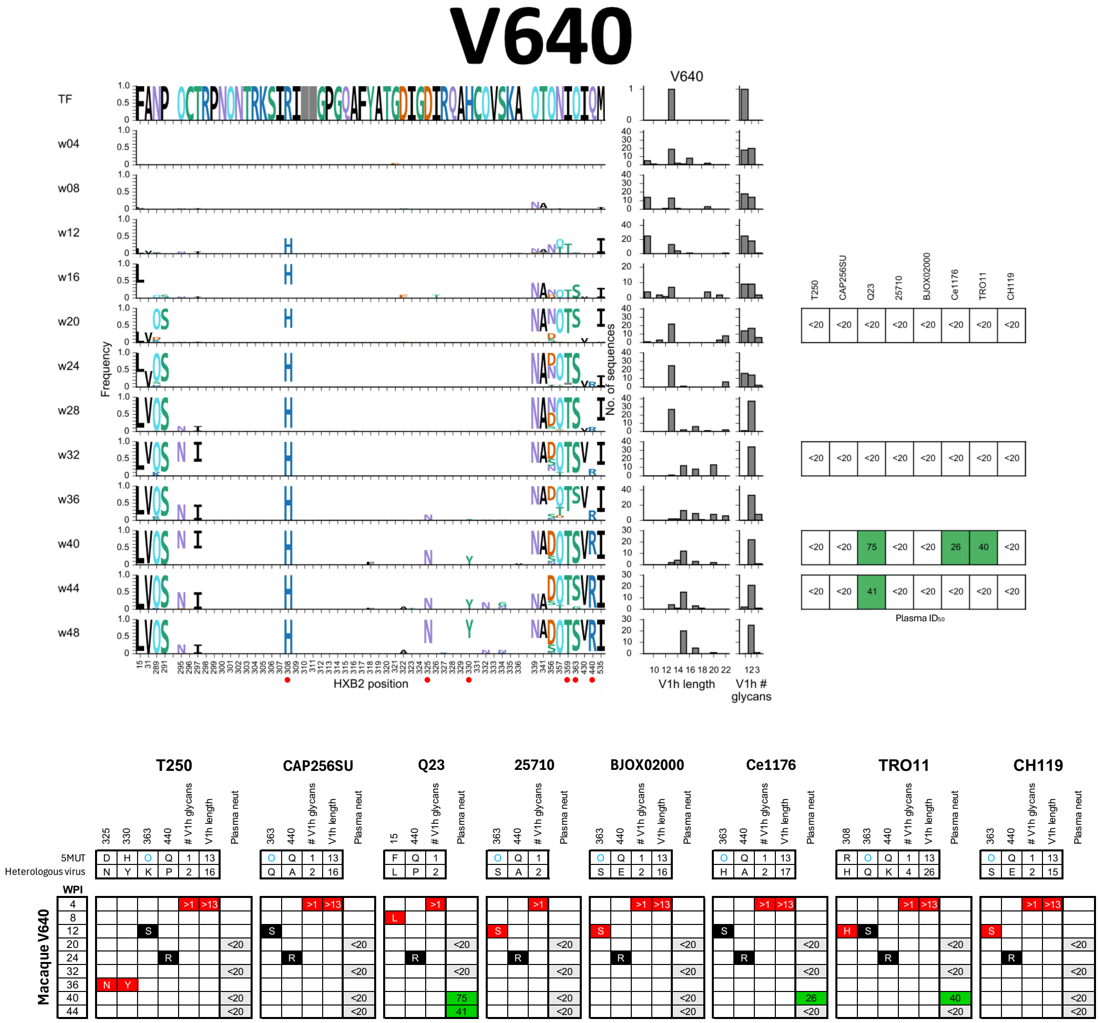


**Fig. S29. Env evolution and development of plasma breadth in rhesus macaque V640.** See **Fig. 6A** and **Fig. 7B** for details regarding figure layout and **Methods** for a description of the analysis. **(A)** Longitudinal patterns of mutations in Env and development of neutralization breadth. **(B)** Integrated information from the three graphs in **(A)**. 7/7 amino acid changes, 2/2 V1h elongations, and 3/3 V1h glycan additions relevant to a given pseudovirus were detected prior to or concurrent with neutralization breadth being observed in the plasma.

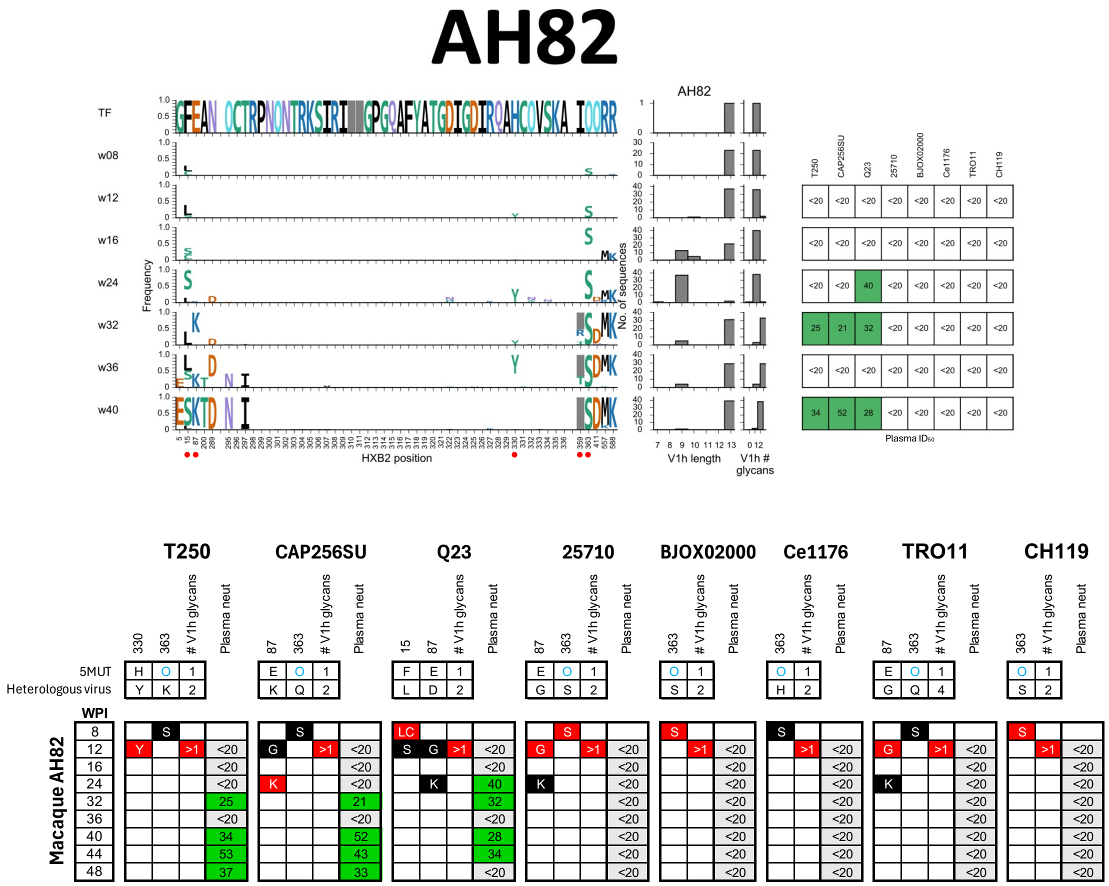


**Fig. S30. Env evolution and development of plasma breadth in rhesus macaque AH82.** See **Fig. 6A** and **Fig. 7B** for details regarding figure layout and **Methods** for a description of the analysis. **(A)** Longitudinal patterns of mutations in Env and development of neutralization breadth. **(B)** Integrated information from the three graphs in **(A)**. 6/6 amino acid changes and 3/3 V1h glycan additions relevant to a given pseudovirus were detected prior to or concurrent with neutralization breadth being observed in the plasma. O363S was very rare at week 8 but became common by week 24. Addition of a second V1h glycan was first observed at week 12, but this variant was very rare until week 32, coincident with detection of plasma neutralization of T250 and CAP256SU, both of which have two V1h glycans. V1h elongation was not observed within the first 48 weeks of infection in this animal and thus was not included in this analysis. V1h length could be contributing to the limited breadth and potency observed in this animal, although other factors may also play a role. Notably, 25710 has a short V1h loop but is not recognized by the plasma.

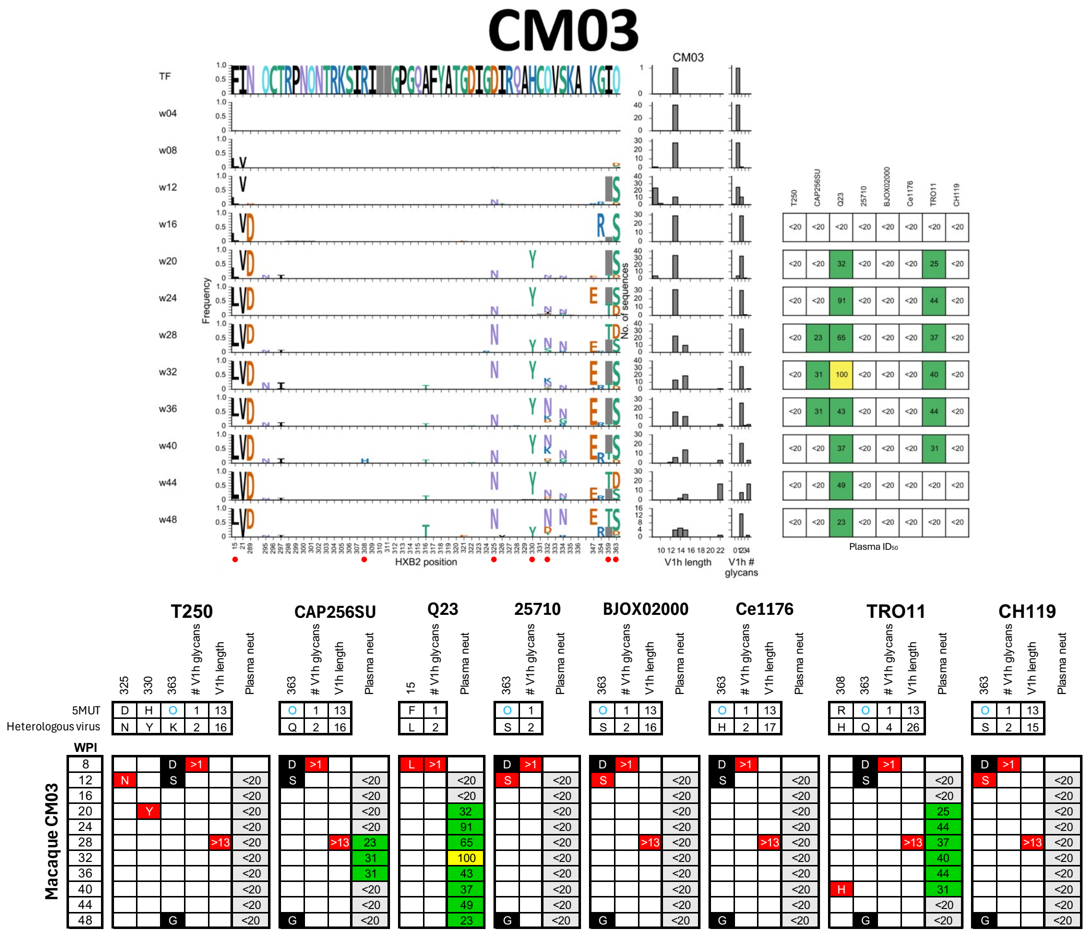


**Fig. S31. Env evolution and development of plasma breadth in rhesus macaque CM03.** See **Fig. 6A** and **Fig. 7B** for details regarding figure layout and **Methods** for a description of the analysis. **(A)** Longitudinal patterns of mutations in Env and development of neutralization breadth. **(B)** Integrated information from the three graphs in **(A)**. 3/4 amino acid changes, 1/2 V1h elongations, and 3/3 V1h glycan additions relevant to a given pseudovirus were detected prior to or concurrent with neutralization breadth being observed in the plasma. TRO11 was neutralized at week 20 even though the R308H variant was not observed until week 40 and elongated V1 variants were not observed until week 28. While R308H seemed to be a resistance mutation for most other bNAb lineages, neither AH82 nor CM03 showed much selection involving this residue during the first year of infection, suggesting that this site might not be relevant to the bNAb lineages in these two animals. V1-elongated variants were detected concurrently with plasma neutralization of CAP256SU.

**Table S1. Rhesus macaque demographic information.**

| Group | RM ID | Sex | Age | CD8 depletion |
|---|---|---|---|---|
| 1 | V630 | Male | 5 | anti-CD8β (25 mg/kg) |
| | V631 | Male | 5 | |
| | V632 | Male | 5 | |
| | V633 | Male | 5 | |
| | V634 | Male | 5 | |
| | V635 | Male | 5 | |
| | V636 | Male | 5 | |
| | V637 | Male | 5 | |
| 2 | V638 | Male | 5 | |
| | V639 | Male | 5 | |
| | V640 | Male | 5 | |
| | V641 | Male | 5 | |
| | V642 | Male | 5 | |
| | V643 | Male | 5 | |
| | V644 | Male | 5 | |
| | V645 | Male | 5 | |
| 3 | V650 | Male | 5 | |
| | V651 | Male | 5 | |
| | V652 | Male | 5 | |
| | V653 | Male | 5 | |
| | AG99 | Female | 21 | None |
| | AH82 | Female | 20 | |
| | AJ09 | Female | 20 | anti-CD8α (25 mg/kg) |
| | AM12 | Female | 21 | |
| | CK36 | Female | 8 | anti-CD8β (25 mg/kg) |
| | CM03 | Female | 8 | |
| | NN39 | Male | 3 | |
| | NN58 | Male | 3 | |

**Table S2. Cryo-EM data collection and processing parameters.**

**Data collection parameters**

| | | |
|---|---|---|
| Microscope | Talos Arctica | Titan Krios |
| Voltage (kV) | 200 | 300 |
| Camera | Gatan K3 | Gatan K3 |
| Magnification | 45000x | 105,000x |
| Frames per movie | 40 | 40 |
| Recording mode | Counting | Counting |
| Dose rate ($e^-$/pixel/s) | 21.234 | 20.453 |
| Total electron dose ($e^-/Å^2$) | 45 | 60 |
| Defocus range (µm) | -0.8 to -1 | -0.8 to -1 |
| Pixel size (Å) (super-resolution) | 0.435 | 0.416 |
| FSC threshold | 0.143 | 0.143 |

**EMDB Maps and Data Processing Parameters**

| Antibody | Antigen | No. of Movies | No. of extracted particles | Symmetry imposed | Map resolution (Å) | PDB | EMDB |
|---|---|---|---|---|---|---|---|
| AJ09-21 | 5MUT-3fill DS | 3483 | 186,421 | C3 | 3.2 | 9YHO | EMD-72969 |
| AJ09-83 | 5MUT-3fill DS | 5560 | 281,473 | C3 | 3.0 | 9YHQ | EMD-72970 |
| AJ09-110 | 5MUT-3fill DS | 4499 | 144,442 | C3 | 3.1 | 9YHR | EMD-72971 |
| AM12-340 | 5MUT-3fill DS | 1065 | 36,140 | C3 | 3.9 | 9YHS | EMD-72972 |
| AM12-347 | 5MUT-3fill DS | 2197 | 115,762 | C3 | 3.6 | 9YHT | EMD-72973 |
| AM12-351 | 5MUT-3fill DS | 1182 | 87,733 | C3 | 3.5 | 9YIB | EMD-72985 |
| AM12-352 | 5MUT-3fill DS | 2291 | 183,947 | C3 | 3.7 | 9YID | EMD-72986 |
| V645-158 | 5MUT-3fill DS | 1822 | 150,394 | C3 | 3.8 | 9YIE | EMD-72987 |
| V634-136 | 5MUT-3fill DS | 3912 | 115,334 | C3 | 3.4 | 9YIF | EMD-72988 |
| V634-136-UCA | del4-3fill DS | 4288 | 235,715 | C1 | 3.8 | 9YIG | EMD-72989 |
| NN39-25 | 5MUT-3fill DS | 2435 | 342,870 | C3 | 4.0 | 9YIH | EMD-72990 |
| NN39-171 | 5MUT-3fill DS | 4889 | 634,261 | C3 | 3.8 | 9YII | EMD-72991 |
| N/A | 5MUT-3fill DS | 1492 | 548,931 | C3 | 3.8 | 9YIJ | EMD-72992 |
| N/A | del4-3fill DS | 7551 | 302,556 | C3 | 2.9 | 9YIK | EMD-72993 |
| N/A | del8-3fill DS | 2303 | 272,897 | C3 | 3.5 | 9YIL | EMD-72994 |

### Table S3. Human V3-glycan bNAb UCA sequences.

| mAb ID | Heavy chain sequence | Light chain sequence | Notes |
|---|---|---|---|
| DH270-UCA3 | QVQLVQSGAEVKKPGASVKVSCKASGYTFTGYYMHWVRQAPGQGLEWMGWINPNSGGTNYAQKFQGRVTMTRDTSISTAYMELSRLRSDDTAVYYCARGGWISLYYDSSGYPNFDYWGQGTLVTVSS | QSALTQPASVSGSPGQSITISCTGTSSDVGSYNLVSWYQQHPGKAPKLMIYEVSKRPSGVSNRFSGSKSGNTASLTISGLQAEDEADYYCCSYAGSSTVIFGGGTKLTVL | Reported in ref (19) |
| BF520-UCA_revised | QVQLVQSGAEVKKPGASVKVSCKASGYTFTGYYMHWVRQAPGQGLEWMGWINPNSGGTNYAQKFQGRVTMTRDTSISTAYMELSRLRSDDTAVYYCARGPFPNYYGSGSYWGGLDYWGQGTLVTVSS | EIVMTQSPATLSVSPGERATLSCRASQSVSSNLAWYQQKPGQAPRLLIYGASTRATGIPARFSGSGSGTEFTLTISSLQSEDFAVYYCQQYNNWPSPFTFGPGTKVDIK | Re-inferred using available data |
| PCDN76-UCA_revised | QVQLQQWGAGLLKPSETLSLTCAVYGGSFSGYYWSWIRQPPGKGLEWIGEINHSGSTNYNPSLKSRVTISVDTSKNQFSLKLSSVTAADTAVYYCARGGRKVYYDFWSGYVNNWFDPWGQGTLVTVSS | EIVLTQSPGTLSLSPGERATLSCRASQSVSSSYLAWYQQKPGQAPRLLIYGASSRATGIPDRFSGSGSGTDFTLTISRLEPEDFAVYYCQQYGSSPTFGQGTKVEIK | Re-inferred using available data |
